# Evolutionary dynamics in the Irano-Anatolian and Caucasus biodiversity hotspots: Evolutionary radiation and its drivers in *Gypsophila* (Caryophyllaceae)

**DOI:** 10.1101/2023.11.24.568494

**Authors:** Hossein Madhani, Richard K. Rabeler, Guenther Heubl, Elizabeth Stacy, Navid Madhani, Shahin Zarre

**Author notes:** Author for correspondence: Hossein Madhani.

## Abstract

- Diversification rates vary through time, with bursts of rapid diversification underscoring species richness across the Tree of Life. Yet the abiotic and biotic factors that correlate with periods of rapid diversification are not well known.
- We explored the diversification dynamics of *Gypsophila* (Caryophylleae: Caryophyllaceae), a large and diverse genus with a high degree of endemism in the Irano-Anatolian and Caucasus biodiversity hotspots. We tested the hypothesis of young and recent diversification of biodiversity in these two hotspots and investigated the biotic and abiotic, and paleoenvironmental cofactors of diversification within the Caryophylleae tribe, with a special focus on *Gypsophila*.
- Our analysis identified multiple shifts in the diversification rate within Caryophylleae, including a newly identified shift in *Gypsophila*, which began approximately 3 million years ago and was triggered by both paleoenvironmental factors and morphological novelties. These novelties were enabled by explosive diversification of woodiness and perenniality and were facilitated through the high adaptability of *Gypsophila* to gypsum soil and the harsh environments of alpine habitats.
- These findings reveal the highly dynamic evolutionary history of both the Caryophylleae clade and *Gypsophila*, consistent with the extensive historical fluctuations in the geology and climate of the Irano-Anatolian and Caucasus biodiversity hotspots. This study significantly improves our understanding of the dynamics of evolution in the Irano-Anatolian and Caucasus biodiversity hotspots and highlights the impact of environmental changes and morphological innovations on diversification rates.

## Introduction

Variation in species richness among lineages and among environments is a long-standing problem in evolutionary biology. Studies of the dynamics of diversification show episodes of increased diversification rate in all major lineages of the Tree of Life, and evolutionary radiation is known to play a major role in the emergence of species-rich clades (Baldwin & Sanderson, 1998; Simões *et al*., 2016; Wiens, 2017; Naciri & Linder, 2020). Historic geological and climatological events like orogenic activities, and climate fluctuations have been shown to change the dynamics of diversification, dispersal, and consequently the patterns of biodiversity (Hughes & Eastwood, 2006; Antonelli *et al*., 2009, 2018; Rahbek *et al*., 2019). Topology-driven isolation in island-like systems, such as mountains (so-called ‘sky islands,’), lakes, and valleys, increases the speciation rate and results in high endemism (Gillespie & Roderick, 2014; Steinbauer *et al*., 2016; Antonelli *et al*., 2018; Rahbek *et al*., 2019). A growing body of research corroborates that the exceptional diversity and species endemic to alpine habitats evolved rapidly and relatively recently due to evolutionary radiation (Hughes & Atchison, 2015). This phenomenon has been reported across several mountain systems, including the Andes Mountains in South America (Hughes & Eastwood, 2006; Nürk *et al*., 2013; Madriñán *et al*., 2013; Lagomarsino *et al*., 2016; Pérez-Escobar *et al*., 2017), Rockies and Sierras mountains in Western North America (Wolfe *et al*., 2006; Tank & Olmstead, 2008; Drummond *et al*., 2012), New Zealand and New Guinea mountains (Winkworth *et al*., 2005; Brown *et al*., 2006; Joly *et al*., 2014), European alpine systems (Comes & Kadereit, 2003; Kadereit *et al*., 2004; Roquet *et al*., 2013), Eastern African highlands (Knox & Palmer, 1995; Gehrke & Linder, 2009; Linder, 2014), alpine habitats of Himalaya mountains and Tibetan Plateau (Liu *et al*., 2006; Jabbour & Renner, 2012; Sun *et al*., 2012; Wen *et al*., 2014; Zhang *et al*., 2014; Favre *et al*., 2015; Xing & Ree, 2017), and Irano-Turanian alpine regions (López-Vinyallonga *et al*., 2009; Moharrek *et al*., 2019).

Biodiversity hotspots (Mittermeier *et al*., 2011; Noss *et al*., 2015), because of their exceptional species richness and unique ecological conditions, are perfect for understanding the dynamics of diversification. Recent studies of the evolutionary dynamics of species-rich lineages in biodiversity hotspots have increased our understanding of the importance of evolutionary radiations and their drivers in shaping the biodiversity of these endangered ecosystems (Table S1). Despite the growing body of research on evolutionary radiations across many of the 36 biodiversity hotspots (Mittermeier *et al*., 2011; Noss *et al*., 2015), the dynamics of diversification in several of these regions remains unexplored. Moreover, the cofactors of rapid diversification and evolutionary radiation are still largely unknown for most biodiversity hotspots (Table S1). Poor understanding of these regions is due largely to insufficient sampling of endemic species and a dearth of resolved phylogenies. Understanding the cofactors of diversification, in particular, may be useful in the management of these important regions in the face of climate change, which accelerates extinction in these endangered regions (Malcolm *et al*., 2006; Fonseca, 2009; Le Roux *et al*., 2019).

### Irano-Anatolian and Caucasus biodiversity hotspots

Home to about 12,400 plant species, including over 30% endemics, the Irano-Anatolian and Caucasus biodiversity hotspots are classified among the 36 global biodiversity hotspots (Mittermeier *et al*., 2011). These two adjacent hotspots are predominantly mountainous areas in South-West (SW) Asia and part of the Irano-Turanian (IT) floristic region, one of the richest floristic regions in the world with a dramatic geological and climatic history (Zohary, 1973; Takhtadzhian *et al*., 1986; Davis *et al*., 1994; Djamali *et al*., 2012a; Manafzadeh *et al*., 2017). Alpine and sub-alpine ecosystems of Turkish-Iranian plateau and Caucasus mountains, and proximity of these high altitudes to vast deserts such as Dasht-e Kavir and Dasht-e Lut, as well as large bodies of water such as Caspian, Black and Mediterranean seas contribute chiefly to the natural landscape of the Irano-Anatolian and Caucasus biodiversity hotspots. Geographically, the area covers the Turkish-Iranian plateau, characterized by several major mountain ranges, i.e. Taurus Mountains, Armenian Highlands, Zagros Mountains, Alborz Mountains, and Kopet-Dag mountain range, all recognized as the main centers of biodiversity in the IT region (Zazanashvili *et al*., 2004; Zazanashvili, 2009; Djamali *et al*., 2012b; Manafzadeh *et al*., 2014, 2017; Parolly, 2015; Noroozi *et al*., 2018).

Two main tectonic activities linked with the India–Asia collision in the East, and the Arabia– Eurasia collision in the West, influenced the geography, topography, and climate history of the IT region from the early Eocene to the current time (Hsü *et al*., 1977; Zachos *et al*., 2001, 2008; van Dam, 2006; Allen & Armstrong, 2008; Ballato *et al*., 2010; Djamali *et al*., 2012a; Manafzadeh *et al*., 2017). The first major tectonic activity in the India–Asia collision started in the early Eocene with the disappearance of the Tethys Ocean and promoted the emergence of the western parts of the IT region by the end of Eocene and early Oligocene (Báldi, 1980; Şengör & Yilmaz, 1981; Rögl, 1997, 1999; Meulenkamp & Sissingh, 2003; Popov *et al*., 2006). By the end of the Oligocene, the India–Asia collision resulted in the formation of the Tibetan plateau, and the subsequent Arabia–Eurasia collision caused the uplift of the Irano-Anatolian plateau, and Caucasus mountains (Şengör & Kidd, 1979; Şengör *et al*., 1985; Pearce *et al*., 1990; Keskin, 2003; Vincent *et al*., 2007; Dilek *et al*., 2010; Yin, 2010; Hatzfeld & Molnar, 2010; Gavillot *et al*., 2010; Mouthereau *et al*., 2012). These geologic events along with climatic fluctuations, such as cooling and aridification episodes during the Eocene-Oligocene transition, warming phases in the late Oligocene and the Middle Miocene, the late Miocene cooling, aridification events in the Pliocene, the Messinian Salinity Crisis, and the Quaternary glacial stages, all had major impacts on the patterns of biodiversity in the area (Hsü *et al*., 1977; Zachos *et al*., 2001, 2008; van Dam, 2006; Allen & Armstrong, 2008; Ballato *et al*., 2010; Djamali *et al*., 2012a; Manafzadeh *et al*., 2017). The orogenic activities and subsequent climate changes promoted a high degree of fragmentation and isolation in the alpine flora, which caused high levels of alpine endemism in these regions (Noroozi *et al*., 2008; Manafzadeh *et al*., 2017).

**Genus *Gypsophila* L.** is one of the largest and most morphologically heterogeneous genera in the family Caryophyllaceae Lam. & DC, including approximately 150 species (Barkoudah, 1962; Hernández-Ledesma *et al*., 2015; Madhani *et al*., 2018) of annual or perennial herbaceous/woody, creeping, or cushion-forming plants. Members of the genus inhabit primarily the mountainous steppes in the Holarctic region, but most of the species are restricted to the high-elevation habitats of western parts of the IT region in the Irano-Anatolian and Caucasus biodiversity hotspots (Barkoudah, 1962; Huber-Morath, 1967; Rechinger, 1988). This high degree of endemism along with the remarkable morphological diversity and resilience of *Gypsophila* members to harsh environments imply the evolutionary adaptation of *Gypsophila* spp. to the alpine habitats of these two biodiversity hotspots. Molecular phylogenetic studies during the last decade resolved the problematic taxonomic boundaries of *Gypsophila* within the tribe Caryophylleae, elucidated phylogenetic relationships and limits of genera in this group (Greenberg & Donoghue, 2011; Pirani *et al*., 2014, 2020; Madhani *et al*., 2018, 2024; Noroozi *et al*., 2020; Fassou *et al*., 2022), and paved the way for exploring the dynamics of diversification and its cofactors inside *Gypsophila* and within Caryophylleae.

Given the young geological age of alpine and sub-alpine regions in the Irano-Anatolian and Caucasus biodiversity hotspots, coupled with the recent geological and climatic changes in this region, we posit that the rich variety of species unique to the mountainous regions of these hotspots results from a recent acceleration in diversification rate. We test this hypothesis, and examine cofactors of diversification, using *Gypsophila* L., a highly diverse alpine genus, almost half of which (∼70 spp) are endemic to alpine and subalpine habitats of the Irano-Anatolian and Caucasus biodiversity hotspots. The goals of this study are: 1) to clarify the evolutionary history within *Gypsophila* using phylogenetic reconstruction and molecular dating, 2) to investigate tempo and rates of diversification within Caryophylleae and *Gypsophila*, 3) to assess the impacts of biotic and abiotic factors on diversification rates in *Gypsophila*, and 4) to reconstruct the ancestral ranges of biogeographic pathways within Caryophylleae and its major clades.

## Materials and Methods

### Taxon sampling, DNA extraction, PCR, and sequencing

About half of all *Gypsophila* species were sampled, including representatives from all previously recognized sections, and representatives of the full breadth of morphological variation within the genus. Species were sampled also from throughout the full geographic range of the genus, minus Australia (i.e., *G. australis* (Schltdl.) A.Gray.). We generated new DNA sequences for 40 species of *Gypsophila* and supplemented these sequences with sequences from Madhani *et al*. (2018) and GenBank for a total of 73 species of *Gypsophila*. We also added some members of all other genera within Caryophylleae in our study with a special focus on *Dianthus*. Altogether we prepared three datasets: the internal transcribed spacer (ITS) region of the ribosomal cistron (consisting of ITS1, the intervening 5.8S gene, and ITS2) for 184 accessions of Caryophylleae including 72 species of *Gypsophila*; the *rps16* intron for 105 accessions from Caryophylleae including 55 species of *Gypsophila*; and the *mat*K gene for 146 accessions from all 11 tribes and main genera of the family Caryophyllaceae including 25 species of *Gypsophila*, 37 species of *Dianthus*, mostly sequenced by Greenberg & Donoghue (2011), and 22 species of *Saponaria*. All DNA extractions, PCR amplification, and sequencing were conducted as described previously in Madhani *et al*. (2018). Information on voucher specimens and GenBank accession numbers are provided in Supporting Information Table S2.

### Sequence alignment and phylogenetic reconstruction

First, we edited sequences in Geneious v.8.0.5 (Kearse *et al*., 2012) and performed the multiple sequence alignments using MAFFT v.7 with default parameters (Katoh & Standley, 2013). The results were manually corrected using Mesquite v.3.7 (Maddison & Maddison, 2021). We used Bayesian inference (BI) and maximum parsimony (MP) to reconstruct the phylogenetic history of the group. The general time-reversible model with gamma-shape rate variation and a proportion of invariable sites (GTR + I + Γ) is estimated as the optimal substitution model for *mat*K and ITS datasets, and GTR + Γ is estimated as the best model for *rps16* using the Akaike information criterion (AIC) implemented in jModelTest v.2.1.6 (Darriba *et al*., 2012). We used MrBayes v.3.2.7a (Ronquist & Huelsenbeck, 2003) with the default condition of three “heated” and one “cold” chain and 40,000,000 generations. Trees were sampled every 1000 generations, and 0.25 of pre-stationarity MCMC samples were discarded as burn-in, which included 10000 samples in each run. The convergence between runs and ESS values were assessed using TRACER v.1.7.2 (Rambaut *et al*., 2018). Finally, we summarized the remaining trees in a 50% majority-rule consensus tree for each dataset. To reconstruct phylogenies by the MP method, we used PAUP* v.4.0a168 (Swofford, 1993) with the following parameters: all characters unordered and equally weighted, heuristic search with random sequence addition, tree-bisection-reconnection branch swapping, 100 random-addition-sequence replicates, MAXTREES option set to 10000, and resulting trees summarized in a majority-rule consensus tree. For bootstrapping we used the maximum likelihood method as implemented in RAxML-HPC2 on XSEDE v.8 (Stamatakis, 2014) with the GTRCAT model and 1000 bootstrap replicates. All phylogenetic analyses were done on the CIPRES server (Miller *et al*., 2010).

### Missing species survey

To account for missing species of *Gypsophila* in our analyses and to obtain more precise estimates of diversification rates associated with individual biotic (life strategy, calyx shape, life form) and abiotic (habitat and elevation) traits, we performed a comprehensive taxonomic survey of all accepted species names of *Gypsophila* (Table S26). This taxonomic survey included four years of fieldwork during the flowering and fruiting seasons of *Gypsophila* in high-elevation habitats of Zagros and Alborz mountains and examination of more than 1000 herbarium sheets. All missing species were assigned to a particular trait group by examining protologues, monographs, and floras, such as Barkoudah (1962), Flora of Turkey (Huber-Morath, 1967), and Flora Iranica (Rechinger, 1988). We also searched major available online resources (Tropicos, http://www. tropicos.org/; JSTOR Global Plants, https://plants.jstor.org; Global Biodiversity Information Facility - GBIF, www.gbif.org; and Integrated Digitized Biocollections - iDigBio, www.idigbio.org), as well as websites of several individual herbaria (BM, BR, E, G, GH, K, KEW, L, LINN, OS, P, US, WU: herbarium abbreviations follow Thiers 2023+). Information on all currently recognized *Gypsophila* species is presented in Table S26.

### Molecular dating

The *mat*K tree was used to infer the age of Caryophylleae. To calibrate the *mat*K tree, we used the only studied fossil of the family, *Caryophylloflora paleogenica* G. J. Jord. & Macphail, which is inferred as the sister to either one or both of the subfamilies, Alsinoideae and Caryophylloideae (Jordan & Macphail, 2003). The age of the strata in which the fossil was found is inferred as middle to late Eocene, 48.6–33.9 Ma (http://www.paleodb.org). The monophyly enforced to Caryophyloideae and Alsinoideae subfamilies with lognormal distribution, mean = 0.0, offset = 33.9, and SD = 1.37 in BEAST v.1.10.4 (Suchard *et al*., 2018), which sets a hard minimum boundary for the age of the group and allows the maximum ages in prior distribution to extend beyond the maximum age of the strata (Frajman *et al*., 2009; Ho & Phillips, 2009). We selected GTR + I + Γ as the best-fitting model for the *mat*K dataset and a relaxed clock for preliminary analysis, as the coefficient of variation frequency histogram did not abut zero, and the standard deviation of the uncorrelated lognormal relaxed clock did not include zero. Thus there was among-branch rate heterogeneity, and a strict molecular clock could be rejected for the *mat*K dataset (Drummond *et al*., 2007). The BEAST analyses were run for both fossilized birth-death and Calibrated Yule models as the tree priors to infer the divergent time of Caryophylleae and the main genera inside it. All input XML files for BEAST analyses were prepared using BEAUti v.1.10.4, and all dating analyses were performed on the CIPRES server (Miller *et al*., 2010).

The inferred age for Caryophylleae from fossil calibration of the *mat*K analysis was used as the secondary calibration point to calibrate the *rps16* and ITS phylogenies. We set a normal distribution with mean=24.68 and SD=3.4 for the Caryophylleae clade, which corresponds to 95% highest posterior density intervals (HPDs) of the age of Caryophylleae (30.90‒17.99 Ma). The preliminary BEAST analyses for ITS and *rps16* using an uncorrelated relaxed lognormal clock led us to reject a strict clock, as the coefficient of variation frequency histogram did not include zero (Drummond *et al*., 2007). We thus used a relaxed clock log-normal for the clock model in all subsequent analyses of the *rps16* and ITS datasets. A Calibrated Yule tree prior was used for both datasets, and the analyses were repeated with a Birth-Death tree prior. Independent runs for each dataset with different tree priors were performed for 40 million generations, sampling every 1000 generations. We assessed the accuracy of chain convergence and adequacy of effective sample size (ESS) by inspection of posterior estimates for different parameters and MCMC sampling of the log file for each run in TRACER v.1.7.2 (Rambaut *et al*., 2018). The first 20% of the samples in each run were discarded and tree files combined by LOGCOMBINER v.1.10.4 (Drummond & Rambaut, 2007), and the maximum clade credibility trees were generated using TREEANNOTATOR v.1.10.4. We used the R package STRAP (Bell & Lloyd, 2015), and ver. ICS2013 (Cohen *et al*., 2013) of the international chronostratigraphic chart to plot the maximum clade credibility trees against stratigraphy.

### Net diversification rate

We measured net diversification rates for major clades inside Caryophylleae based on the maximum clade credibility trees obtained from different analyses of the ITS, *mat*K, and *rps16* datasets. We estimated net diversification rates in stem and crown groups using the Magallón & Sanderson method (Magallón & Sanderson, 2001) implemented in the R package GEIGER (Harmon *et al*., 2008).

### Time-dependent diversification

To infer the diversification dynamics of the Caryophylleae clade, and to identify shifts in diversification rates, we used a time-dependent model implemented in BAMM v.2.5.0 (Rabosky, 2014). We accounted for incomplete taxon sampling by assigning a sampling fraction of major clades in Caryophylleae for each dataset. We generated priors for BAMM using the setBAMMpriors function, implemented in the R package BAMMtools v.2.1.10 (Rabosky *et al*., 2014) based on the maximum clade credibility tree from the BEAST analysis. Four independent MCMC chains were run in BAMM for 40-million generations and sampled every 10000. We set the prior for the expected number of shifts based on Bayes factor calculations (Mitchell & Rabosky, 2017), and used ESS values (> 200) to assess the convergence of the runs using R package coda v.0.19 (Plummer *et al*., 2006). We then used R-package BAMMtools v.2.1.10 (Rabosky *et al*., 2014) to identify the credible sets of speciation-rate shift and single best-shift configuration, plot rate-through-time curves, and calculate speciation, extinction, and net diversification rates through time for Caryophylleae and its major clades. Further, to test whether (and how) diversification rates varied through time (Morlon *et al*., 2011), we fit likelihood models of time-dependent birth-death functions using the *fit_bd* function implemented in RPANDA v.2.1 (Morlon *et al*., 2016). We fitted 12 different birth-death models to the Caryophylleae, *Gypsophila*, *Dianthus*, *Saponaria*, and *Acanthophyllum* phylogenies for the three datasets (Tables S10-S12). Within each of the three datasets, the best model for each clade was chosen via the corrected Akaike Information Criterion (AICc) and Likelihood Ratio Test (LRT), with alpha set at 0.05, and Yule (λ constant and no μ) as the null model.

### Trait-dependent diversification

To test character-associated diversification within *Gypsophila* we used BiSSE (Maddison *et al*., 2007) models implemented in the R package diversitree 0.9-16 (FitzJohn, 2012). We estimated six rate parameters of the BiSSE analysis for five binary traits: 1) life strategy: annual vs. perennial, 2) habitat: non-montane vs. montane, 3) life form: herbaceous vs. woody, 4) calyx shape: tubiform vs. campanulate/turbinate, and 5) elevation: low vs. high. We estimated BiSSE parameters in these five binary traits by eight different models, each constrained for one or more of the six parameters in the BiSSE analyses (see Tables S4-S8). These six parameters include speciation rates of lineages in states 0 and 1 (λ_0_ and λ_1_, respectively), extinction rates of lineages in states 0 and 1 (µ_0_ and µ_1_, respectively), the transition rate from 0 to 1 (q_01_), and transition rate from 1 to 0 (q_10_). To account for incomplete sampling, the proportion of sampled species in state 0 and state 1 were calculated based on the missing-species survey (Table S26) and specified for each trait as the sampling fractions. We used AIC and the likelihood ratio test by anova function in R to find the best-fitting model for each binary trait. The Chi-square values (ChiSq) and their significance (Pr) were calculated by comparing each model to the null model (minimal model), in which all rates for state 0 are equal to the rates for state 1 (λ_0_=λ_1_; µ_0_=µ_1_; q_01_=q_10_). We ran the MCMC analysis for 100000 steps using an exponential prior for the best-fitting model, applied a burn-in for the first 10% of the steps, and examined the posterior probabilities of the six rates of BiSSE analysis for each of the five binary traits.

To test whether high species richness and endemism in Irano-Anatolian and Caucasus hotspots are driven by spacial asymmetries in speciation, extinction, and dispersal rate or not, we used GeoSSE analysis (Goldberg *et al*., 2011), an extension of BiSSE also implemented in the R package diversitree 0.9-16 (FitzJohn, 2012). In this analysis we considered the two hotspots as one region which is A and anywhere outside the two hotspots as B. Therefore, we estimated the seven rate parameters of the GeoSSE analysis, including speciation and extinction rates within the two biodiversity hotspots (λ_A_ and µ_A_, respectively), speciation and extinction rates outside the two biodiversity hotspots (λ_B_ and µ_B_, respectively), speciation rate between A and B (λ_AB_) is the rate at which widespread lineages diverge into one daughter species endemic to A and another endemic to B, dispersal from Irano-Anatolian and Caucasus hotspots to the rest of the world, or range expansion (d_A_), and dispersal rate into the Irano-Anatolian and Caucasus hotspots from outside, or range contraction (d_B_). We defined 14 models by setting different constraints on these parameters and found the best fitting model using AIC and the likelihood ratio test (see Table S9). We ran the MCMC analysis for 100000 steps using an exponential prior for the best-fitting model, applied a burn-in for the first 10% of the steps, and examined the posterior probability of the seven rates of the GeoSSE analysis.

### Paleoenvironment-dependent diversification

To test for a significant association between paleo-environmental changes and diversification rates of Caryophylleae, *Gypsophila,* and *Dianthus* we used a birth-death model with speciation and extinction varying as a function of temperature changes through geological time (Condamine *et al*., 2013) using *fit_env* function implemented in the R package RPANDA (Morlon *et al*., 2016). We fit 12 different birth-death models (Tables S13-S15). To find the value of the paleoenvironmental variable at each time point, based on deep-sea oxygen isotope records (Zachos *et al*., 2008), we interpolated a smooth line during each birth–death modeling process using the R-package PSPLINE (Ramsey & Ripley, 2013). The speciation and/or extinction rates were then estimated as a function of these values along the dated phylogenies according to the parameters of each model and the best-fitting model identified based on AICc and LRT, as was done for the time-dependent analyses (see above).

### Ancestral area reconstruction

To reconstruct the historical distribution of Caryophylleae we used RASP v.4 (Yu *et al*., 2020). We divided the range of Caryophylleae into five areas, based on the extant distribution of the majority of the tribe and the floristic regions it covers. These areas are western Asia (A), central Asia (B), eastern Asia (C), Europe except the Mediterranean region (D), and the Mediterranean region including North Africa (E). We removed the outgroup from the maximum clade-credibility trees generated in BEAST analyses of the three datasets and used them as the input phylogenies in the historical distribution reconstruction analysis by RASP. To compare different historical biogeography models, we fit the data to all six models implemented in the BioGeoBEARS package in R (Matzke, 2014), including DEC, DEC + j, DIVALIKE, DIVALIKE + j, BAYAREALIKE, and BAYAREALIKE + j. The ancestral distributions at all nodes were reconstructed using the Bayesian Binary Method (BBM: Ronquist, 2004; Yu *et al*., 2015), BAYAREALIKE + j (Matzke, 2014), and Dispersal-Extinction-Cladogenesis model (DEC: Ree & Smith, 2008) implemented in RASP v.4. For the BBM analysis, chains were run for 5,000,000 generations, and states were sampled every 1000 generations. F81+G (estimated Felsenstein 1981 + gamma) was used with null root distribution, and the maximum number of possible ancestral areas was set to five (all possible states) for all analyses.

## 3. Results

Detailed information on the three datasets, including numbers of terminals, variable sites, and parsimony informative sites, is presented in Table 1.

**Table 1.** Alignment characteristics of the three genomic regions (loci) used in the present study.

|  | *matK* | *rps16* | ITS |
| --- | --- | --- | --- |
| Total number of terminals | 146 | 105 | 184 |
| No. of Caryophyllae taxa | 101 | 101 | 180 |
| No. of *Gypsophila* taxa | 25 | 55 | 71 |
| No. of *Dianthus* taxa | 38 | 10 | 40 |
| No. of *Saponaria* taxa | 22 | 7 | 24 |
| No. of *Acanthophyllum* taxa | 6 | 10 | 14 |
| No. of *Bolanthus* taxa | 0 | 8 | 15 |
| Alignment length [bp] | 931 | 993 | 698 |
| Conserved characters [bp] | 342 | 554 | 240 |
| Parsimony-uninformative characters [bp] | 152 | 180 | 108 |
| Parsimony-informative characters [bp] | 437 | 259 | 350 |
| Parsimony-informative characters [%] | 46.9 | 36.1 | 50.1 |

### 3.1. Phylogenetic inference

The results of phylogenetic reconstructions using the three datasets and MP (Figs S1-S3), BI (Figs S4-S6), and RAxML (S7-S9) support the classification system for Caryophylleae in Madhani et al. (2018), and later revisions on the status of *Graecobolanthus* (Zografidis *et al*., 2020), boundaries of *Acanthophyllum* (Pirani *et al*., 2020), the newly introduced genus, *Yazdana* A.Pirani & Noroozi, (Noroozi *et al*., 2020), and the newly resurrected genus *Jordania* (Madhani *et al*., 2024). According to the gained phylogenies, *Gypsophila* includes generic names *Ankyropetalum*, *Vaccaria*, *Bolbosaponaria* (p.p. excl. type), *Dichoglottis*, and *Pseudosaponaria*, and one species of *Diaphanoptera* (Barkoudah, 1962; Madhani *et al*., 2018).

The results of phylogenetic analysis on *rps16* and ITS loci indicate two well supported clades within *Gypsophila*. The first clade, here named **Eugypsophila,** includes the majority of species of *Gypsophila*, as well as the type species of the genus, *G. repens* L. (Figs S2, S3, S5, and S6). Most members of the Eugypsophila clade show the morphological characteristic of *Gypsophila* (baby’s-breath), including a perennial or annual herbaceous habit with many-flowered lax thyrse or panicle inflorescences, sepals scariose at margin and connate at base, and a capsule overtopping the calyx. The second clade, here named the **Pulvinate** clade, which is smaller, comprises plants that are highly adapted to alpine habitats with a pulvinate or dense caespitose life form, tiny succulent or spiny leaves, and usually small few-(uni-)flowered raceme-like monochasia (Figs S1-S9). This clade is most clearly evident in the MrBayes analyses (Figs S4-S6). These two clades which are basically sister, include almost all *Gypsophila* species which we call the **core *Gypsophila*** hereafter (Figs S2-S3, S5-S6, S8-S9). Although we included representatives of all sections of *Gypsophila* recognized by Barkoudah (1962), our results did not support recognizing any additional infrageneric groups.

### Divergence time estimation

The results of the divergence time estimation using fossil calibration on the *mat*K dataset are presented in Table 2 and Fig S10. The results suggest that Caryophylleae started to diverge from the early Oligocene to the early Miocene (31.02–18.04 Ma); these results are similar to those reported by Xue *et al*. (2023). The ages inferred for four large genera inside Caryophylleae, including *Dianthus*, *Gypsophila*, *Acanthophyllum*, and *Saponaria* show that the divergence of their common ancestors started during the late Miocene to early Pliocene (Table 2). The estimated age for the crown group of *Gypsophila*, the second largest genus in the tribe with about 150 accepted species, is 5.31 Ma with SD=1.32 (95% HPD interval: 3.02-7.95), which is surprisingly young given the species richness and morphological diversity of the clade (Table 2). The results of secondary calibration on ITS and *rps16* datasets are presented in Figs S11 and S12, and Table S3. The estimates for divergence time by secondary calibration are older and the credible intervals (CIs) are wider compared to fossil calibration, which is not surprising as wider uncertainty and deviation from primary calibration are usually associated with secondary calibrations (Schenk, 2016).

**Table 2.**
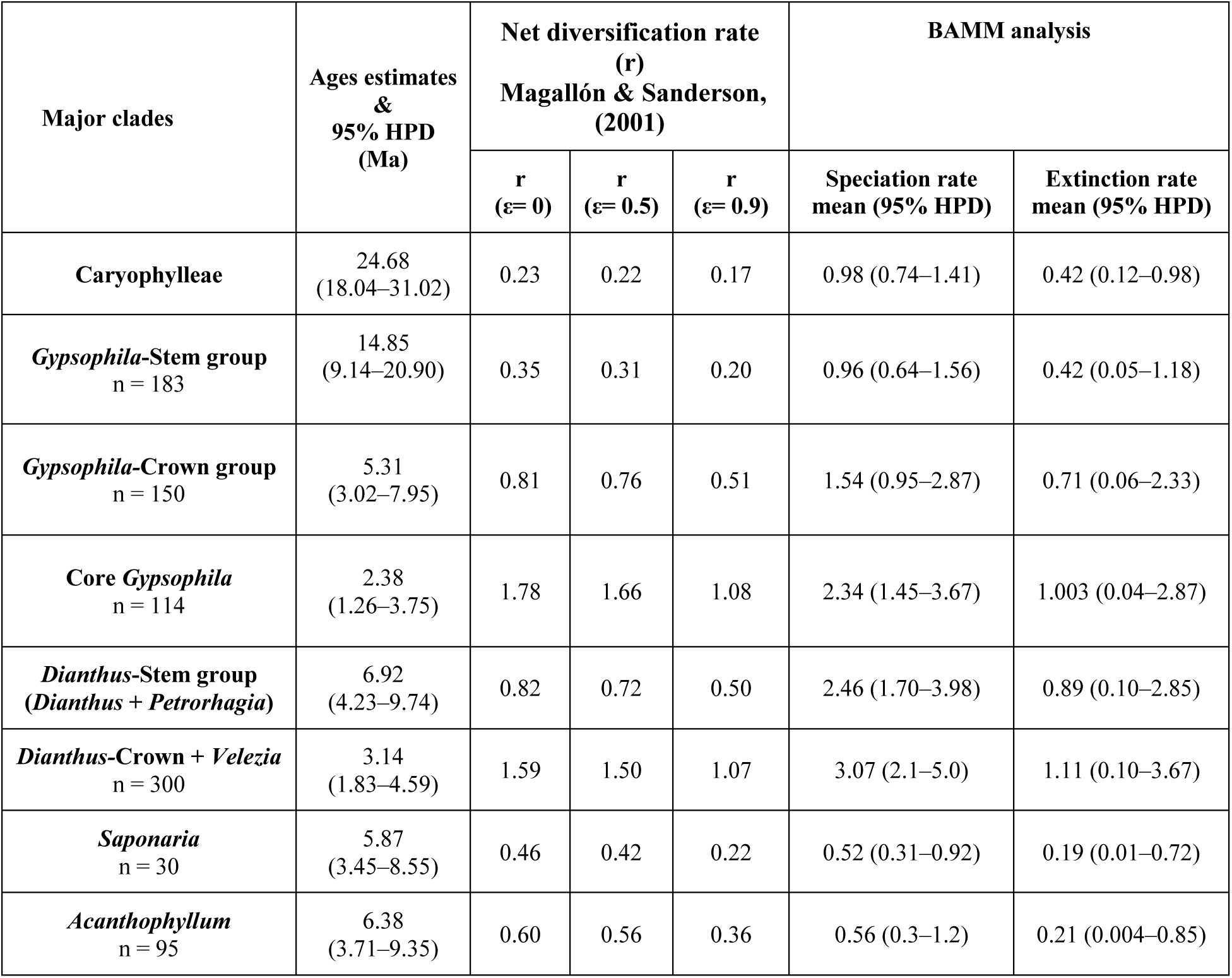
Age estimates for major clades in the Caryophylleae clade using fossil calibration of the *mat*K dataset by BEAST; Net diversification rate estimates using Magallón & Sanderson (2001) method under low (ε= 0), medium (ε= 0.5), and high (ε= 0.9) extinction rates (ε) implemented in geiger package in R; speciation and extinction rates estimated by BAMM.

| Major clades | Ages estimates<br>&<br>95% HPD<br>(Ma) | Net diversification rate<br>(r)<br>Magallón & Sanderson,<br>(2001) |  |  | BAMM analysis |  |
| --- | --- | --- | --- | --- | --- | --- |
| | | r<br>($\epsilon = 0$) | r<br>($\epsilon = 0.5$) | r<br>($\epsilon = 0.9$) | Speciation rate<br>mean (95% HPD) | Extinction rate<br>mean (95% HPD) |
| **Caryophylleae** | 24.68<br>(18.04–31.02) | 0.23 | 0.22 | 0.17 | 0.98 (0.74–1.41) | 0.42 (0.12–0.98) |
| *Gypsophila*-Stem group<br>n = 183 | 14.85<br>(9.14–20.90) | 0.35 | 0.31 | 0.20 | 0.96 (0.64–1.56) | 0.42 (0.05–1.18) |
| *Gypsophila*-Crown group<br>n = 150 | 5.31<br>(3.02–7.95) | 0.81 | 0.76 | 0.51 | 1.54 (0.95–2.87) | 0.71 (0.06–2.33) |
| **Core *Gypsophila***<br>n = 114 | 2.38<br>(1.26–3.75) | 1.78 | 1.66 | 1.08 | 2.34 (1.45–3.67) | 1.003 (0.04–2.87) |
| *Dianthus*-Stem group<br>(*Dianthus* + *Petrorhagia*) | 6.92<br>(4.23–9.74) | 0.82 | 0.72 | 0.50 | 2.46 (1.70–3.98) | 0.89 (0.10–2.85) |
| *Dianthus*-Crown + *Velezia*<br>n = 300 | 3.14<br>(1.83–4.59) | 1.59 | 1.50 | 1.07 | 3.07 (2.1–5.0) | 1.11 (0.10–3.67) |
| *Saponaria*<br>n = 30 | 5.87<br>(3.45–8.55) | 0.46 | 0.42 | 0.22 | 0.52 (0.31–0.92) | 0.19 (0.01–0.72) |
| *Acanthophyllum*<br>n = 95 | 6.38<br>(3.71–9.35) | 0.60 | 0.56 | 0.36 | 0.56 (0.3–1.2) | 0.21 (0.004–0.85) |

### Diversification analyses

The results of time-dependent diversification analysis in BAMM strongly reject a constant diversification rate among the markers used (Bayes factors = 13152, 1142, and 16672 for *mat*K, *rps16*, and ITS datasets, respectively, Tables S20-S22). Instead, at least six distinct credible shifts in speciation rate across the Caryophylleae clade were identified using the three datasets (Figs S13-S15). The best-fitting number of shifts calculated using the stepwise Bayes factor procedure (Mitchell & Rabosky, 2017) shows five, six, and eight shifts for the ITS, *mat*K, and *rps16* datasets, respectively (Tables S23-S25). The best rate shift configurations of the BAMM analysis of all datasets identified at least one shift in the speciation rate of *Dianthus* (consistent with Valente *et al*., 2010) and another shift in *Gypsophila* (Fig S16-S18), and also a shift in *Acanthophyllum* (consistent with Mahmoudi Shamsabad *et al*., 2021) in the *rps16* dataset (Fig S18). The best speciation rate shift configuration within *Gypsophila* was detected at the common ancestor of core *Gypsophila* in the BAMM analyses of all datasets. The rate-through-time plots also indicate that speciation, extinction, and net diversification rates were almost constant following the divergence of the most recent common ancestor (MRCA) of the Caryophylleae clade in the late Oligocene until the early Pliocene (∼ 25-5 Ma, Fig 2b). In the early Pliocene, just before the two identified shifts in speciation occurred in *Gypsophila* and *Dianthus*, all rates started to increase and have continued increasing until the present (Fig. 2b and 2c). Congruent with the shifts identified in these clades, estimates of net diversification rate using the Magallón and Sanderson method also showed extremely high rates of diversification in these large clades within Caryophylleae (Tables 2 and S3). The diversification rates calculated using the Magallón and Sanderson method were 1.08 and 1.78 events *Myr*^-1^ per lineage for core *Gypsophila* under high (e = 0.90) and low (e = 0) extinction fractions, respectively (Table 2), which are among the fastest diversification rates reported for any lineage. Consistent with these calculations, the BAMM analysis also supported very high speciation and extinction rates in all major clades within Caryophylleae (Tables 2 and S3).

**Fig. 1.**
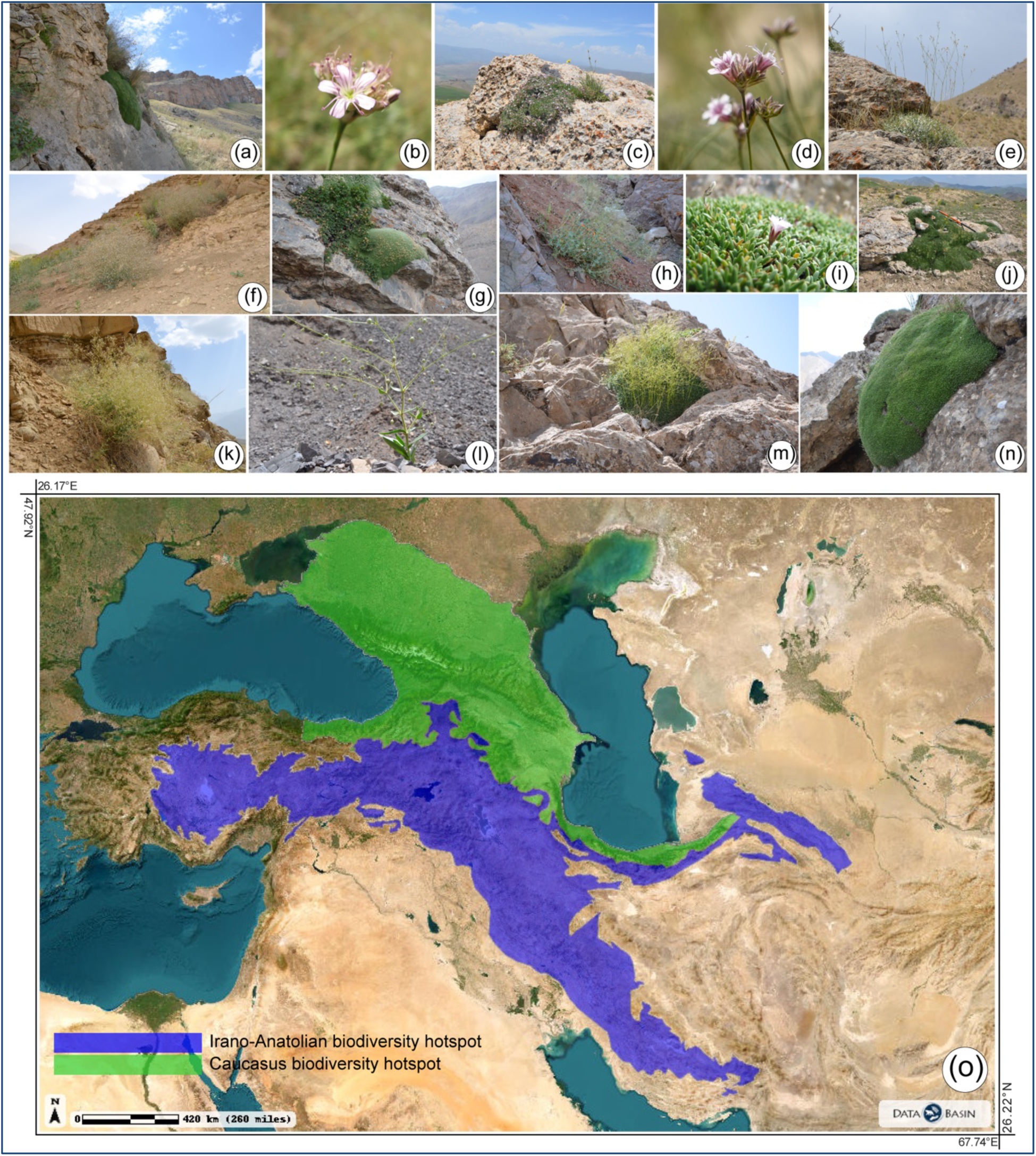
Diversity in *Gypsophila* in its natural habitats in high altitudes of Irano-Anatolian biodiversity hotspot. (a) *Gypsophila pulvinaris*, (b) *Gypsophila caricifolia*, (c) *Gypsophila wilhelminae*, (d and j) *Gypsophila bazorganica*, (e) *Gypsophila graminifolia*, (f) *Gypsophila leioclada*, (g) *Gypsophila aretioides*, (h) *Gypsophila ruscifolia*, (i and n) *Gypsophila saponarioides*, (k) *Gypsophila virgata*, (l) *Gypsophila pilosa*, (m) *Gypsophila acantholimoides* (all photos by H. and N. Madhani), (o) Irano-Anatolian and Caucasus biodiversity hotspots.

**Fig. 2.**
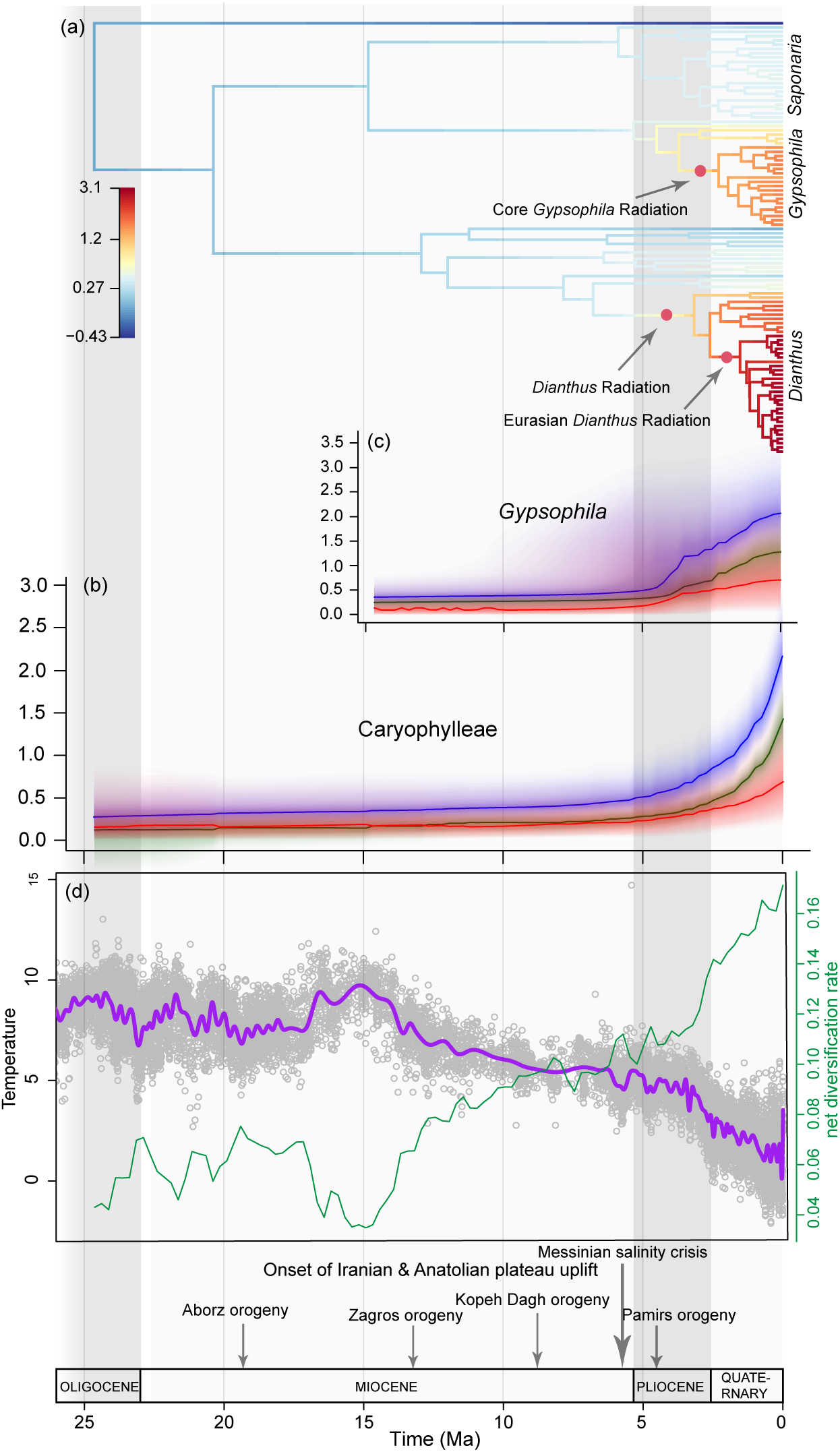
Diversification of Caryophylleae. (a) Net diversification rate plot (phylorate) of the *mat*K dataset showing the best configuration shift identified by BAMM. Evolutionary rate through time for Caryophylleae (b) and *Gypsophila* (c), speciation (blue line), extinction (red line), and net diversification rates (dark green line) shown with color-density shading to indicate 95% confidence intervals. (d) The global temperature through time (purple line) and net diversification rate through time (green line) of the best-fitting model of the temperature-dependent diversification analysis estimated by RPANDA.

Further, of the 12 time-dependent models analyzed for each major clade across all datasets by RPANDA (220 models in total), the best-fitting likelihood model of time-dependent functions indicated constant, yet high, speciation and extinction rates over time across the whole Caryophylleae clade (Table S10). The second (model 6) and third (model 5) best-fitting models, however, both of which were statistically significant, indicated exponential and linear changes, respectively, in speciation rates over time with a constant extinction rate. The best-fitting models for *Dianthus* and *Gypsophila* suggest that speciation rates vary and exhibit an exponential increase over time in all datasets (Tables S10-S12), with the exception of *Gypsophila* in the ITS dataset, where the best-fitting model indicates constant, yet high, speciation and extinction rates over time. Interestingly, the second-best model for this dataset, which also exhibits a significant p-value (4e-04) in the LRT test, shows an exponential increase in speciation rate over time for *Gypsophila* (Tables S12). In the most probable models for *Saponaria* within the ITS and *rps16* datasets, the speciation rate remains constant over time (Tables S11 and S12). In the *Acanthophyllum* clade, the optimal model in the ITS dataset indicates a constant speciation rate throughout all times (Table S12). However, in the *rps16* dataset, the speciation rate aligns with the BAMM results, showing an exponential increase over time (Table S11).

The temperature-dependent analysis rejected the null hypothesis of no influence of temperature on the speciation rate of the Caryophylleae, *Dianthus*, *Gypsophila*, and *Acanthophyllum* clades in most datasets (Tables S13-S15). However, for *Gypsophila* in the *mat*K dataset and *Acanthophyllum* in the ITS dataset, the best models indicate constant speciation and extinction rates unaffected by temperature fluctuations (Table S13). Interestingly, the second-best model for *Gypsophila* within the *mat*K dataset, with a significant p value of 1.7e-04, points towards an exponential relationship between speciation rate and temperature (model 4, Table S13). In contrast, the optimal likelihood models for *Saponaria* showed no association between historical temperature shifts and the diversification rate in either the *rps16* or ITS dataset (Tables S14 and S15).

The best-fitting models derived from the BiSSE analyses for the five binary traits in *Gypsophila* demonstrated higher speciation rates in perennial species compared to annual, in montane species compared to non-montane, in woody species compared to herbaceous, and in species at high elevations compared to those at low elevations (Fig 3a, c-e; Supplementary Tables S4-S6 and S8). According to the best-fitting model for calyx morphology, species with tubiform calyces had higher extinction rates compared to those with campanulate calyces (Fig 3b; Table S7). Lastly, the GeoSSE analysis revealed higher speciation rates within the Irano-Anatolian and Caucasus hotspots, compared to areas outside these two biodiversity hotspots (Fig 3g; Table S9).

**Fig. 3.**
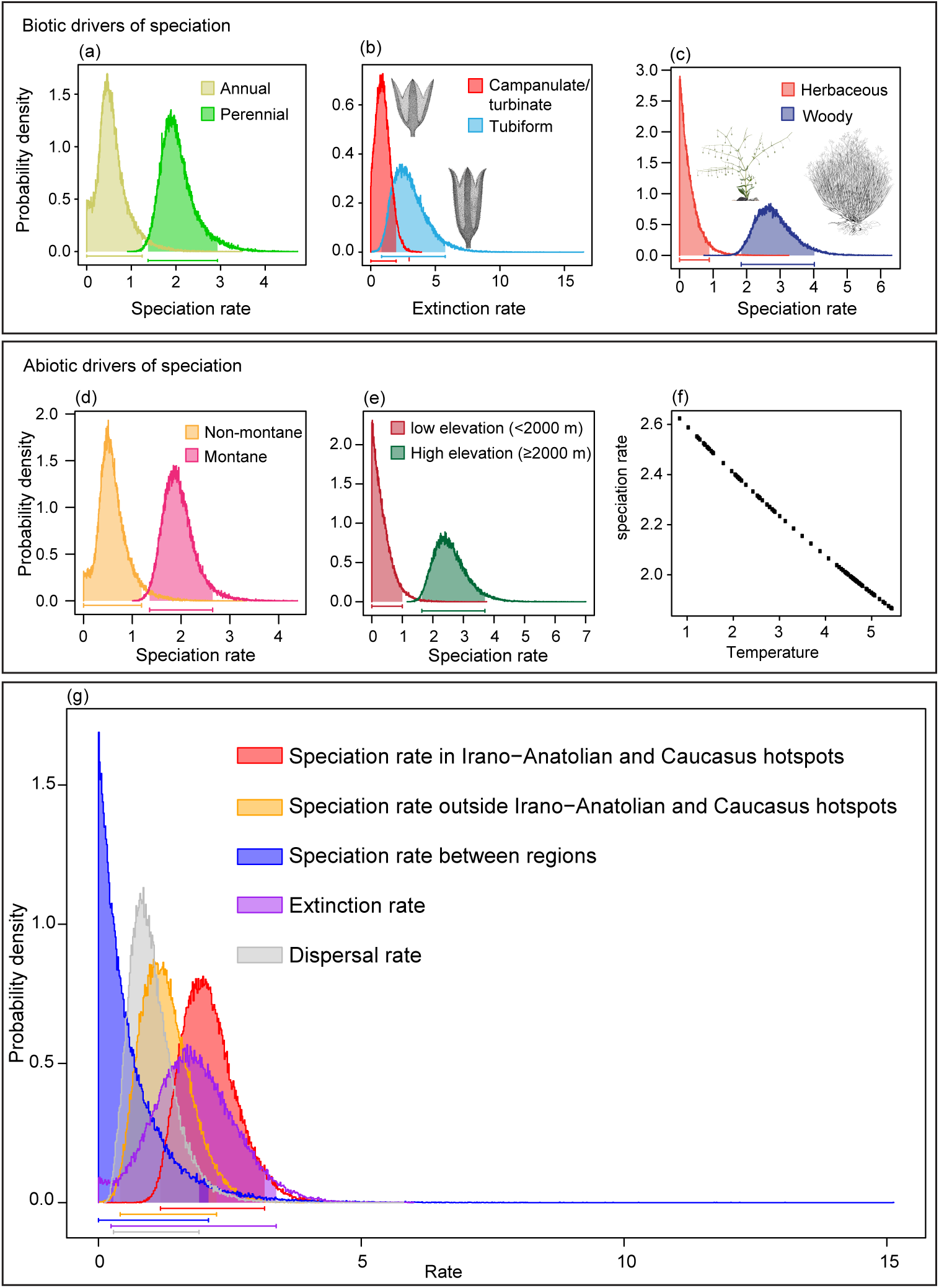
Biotic and abiotic drivers of diversification in *Gypsophila*. Posterior probability plots of speciation/extinction rates of binary traits analyzed by BiSSE: (a) Life strategy (annual vs perennial). (b) Calyx shape (tubiform vs. campanulate/turbinate; illustrated by Shirin Sabeti). (c) Life form (herbaceous vs. woody; illustrated by Shirin Sabeti). (d) Habitat (non-montane vs. montane). (e) Elevation (high elevation vs. low elevation). (f) Speciation rate plot estimated as a function of temperature for the best-fitting model of temperature-dependent analysis of RPANDA for the *Gypsophila* clade. (g) Posterior probability plot of speciation, extinction and dispersal rates per area estimated by GeoSSE.

### Ancestral biogeographic reconstruction

For the biogeographic reconstruction analyses, the BAYAREALIKE + j model was identified as the best-fitting model for all datasets (Table S16); however, the BBM model resulted in higher probabilities for the estimated ancestral ranges and probably better explains the geographical history of the group (Tables S17-S19). Based on the BBM analyses of the ITS and *rps16* datasets (Tables S18 and S19), as well as the BAYAREALIKE + j estimates for ITS (Table S19), western Asia and the Mediterranean region (AE) were inferred as the ancestral area for *Gypsophila* (Fig 4, Tables S17-S19), while the analysis of BBM and BAYAREALIKE + j for *mat*K and *rps16*, respectively, estimated western Asia (80.12%, Table S18) as the ancestral distribution of the MRCA of the genus (Fig 4, Tables S17-S19). Overall, our analysis suggests that dispersal and expansion were more frequent than vicariance in explaining the current geographical distribution of *Gypsophila* (Tables S18 and S19) and Caryophylleae (Table S17).

**Fig. 4.**
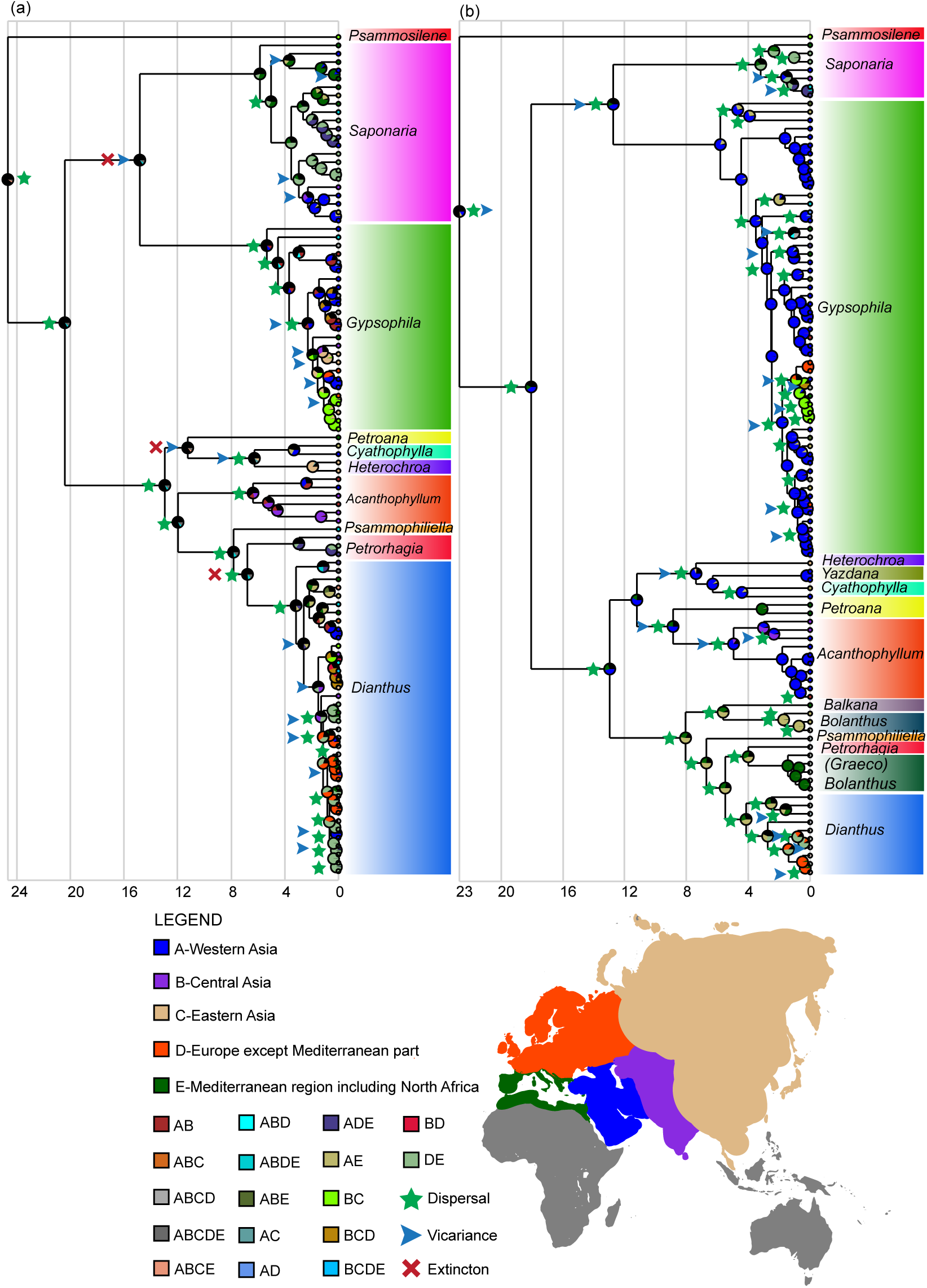
Ancestral area reconstruction of Caryophylleae. The reconstructed ancestral area of the best-fitting model (BAYAREALIKE + j) performed in RASP for the *mat*K (a) and *rps16* (b) datasets. The position of identified dispersal, vicariance, and extinction events are indicated by green stars, blue arrowheads, and red crosses, respectively.

## Discussion

The Irano-Anatolian and Caucasus biodiversity hotspots in the IT floristic region are known for their highly endemic alpine and subalpine flora which correlates with a high degree of topological isolation caused by the recent uplift in the mountain systems in these regions, such as the Turkish-Iranian plateau and its major mountainous components, i.e., Zagros, Alborz, Caucasus, and Kopet-Dag Mountains. Despite the degree of endemism and highly dynamic climatological history of this region, few studies have attempted to characterize the exceptional diversity of endemic taxa in the IT region in terms of molecular clock, biogeography, and dispersal patterns (Manafzadeh *et al*., 2014, 2017; Noroozi *et al*., 2018; Moharrek *et al*., 2019; Ghane-Ameleh *et al*., 2021). The current study is the first to address the issue of mega-diversity in the endemic taxa of Irano-Anatolian and Caucasus biodiversity hotspots through phylogeny, molecular clock, and statistical analyses of diversification.

### Evolutionary radiation in *Gypsophila*

Our study suggests that the majority of the diversity within *Gypsophila* results from recent diversification. Within the large and diverse clade of the Carnation tribe (Caryophylleae), at least three significant, distinct shifts upward in the diversification rate were identified using BAMM analysis, two in *Dianthus* and one in *Gypsophila* (Fig 2 and S13-S18). These recent shifts are consistent with the young ages estimated for both clades based on molecular clock analyses (Tables 2 and S3). The two shifts in diversification rate within the *Dianthus* confirm the evolutionary radiation and estimates for the age of this genus and its Eurasian clade reported by Valente et al (2010). The shift in diversification rate detected in the core *Gypsophila* by BAMM analysis (Fig 2 and S13-S18) is aligned with the surprisingly young age of the crown group of *Gypsophila* (5.31 Ma in the fossil calibration, see Tables 2 and S3), and the secondary calibration of the *rps16* datasets (Table S3). These findings are supported by the best-fitting time-dependent functions selected in RPANDA, which show an exponential increase in speciation rates of *Gypsophila* over time (Tables S10-S12). Our calculation for the net diversification and speciation rates in core *Gypsophila* illuminates one of the fastest evolutionary radiations in the Tree of Life (95% HPD interval of speciation rate: 1.45–3.67 *Myr*^-1^ per lineage, Table 2).

### The impact of orogeny and climate change on species diversification in *Gypsophila*

The beginning of the rise in speciation, extinction, and diversification rates within *Gypsophila*, as depicted in the rate-through-time plot (Fig 2c), coincided with the final phases of the uplift of the major mountain systems in the Irano-Anatolian and Caucasus biodiversity hotspots (Fig 2d). During this period, new alpine environments emerged, creating diverse ecological opportunities that facilitated diversification. Through its impact on aspects such as productivity, the divergence of climatic niches, body size, and metabolic rates, temperature plays a crucial role in shaping biological processes (Condamine *et al*., 2019). This is particularly significant when considering the influence of temperature fluctuations on the rate of speciation. Our paleotemperature-dependency analysis provides evidence of influences of past climate changes on diversification rates across Caryophylleae and its major clades (Tables S13-S15). One interesting finding of this study is the negative correlation between speciation rate in *Gypsophila* and temperature in both the *rps16* and ITS datasets (Tables S14-S15; Fig 3f). The best-fitting paleo-environmental-dependent model in the *mat*K dataset, however, suggested no correlation between temperature and either speciation or extinction rates. Interestingly, the second-best model in this analysis, which was also statistically significant, supported a negative relationship between speciation rate and past temperature changes (Model 4, Table S13). These findings suggest that the diversity within *Gypsophila* and other major clades within Caryophyllaceae was driven by global cooling from the Miocene to the present. An association between the recent global cooling (during the last 15 million years) and increased diversification rate has been suggested also in the terrestrial orchid subfamily Orchidoideae (Thompson *et al*., 2023), and in many subclades within the large rosid clade (Sun *et al*., 2020).

### Sky Islands of Western Asia and Alpine Radiation in *Gypsophila*

Our trait-dependent analyses indicate a significant influence of morphological and ecological characteristics of lineages on diversification rates (Fig 3; Tables S4-S8). Echoing the patterns observed in other alpine radiations, such as *Lupinus* L. (Drummond *et al*., 2012), our study identified a notably higher diversification rate among perennial species compared to their annual counterparts, with perennials experiencing a five-fold increase in speciation rate (Fig 3a; Table S5). Additionally, we observed a significantly higher speciation rate in woody species over herbaceous ones (Fig 3c; Table S6). These findings support the hypothesis that *Gypsophila* species have adapted rapidly to the harsh alpine habitats of the Irano-Anatolian and Caucasus hotspots— environments unfavorable to ancestral annual and herbaceous species. Adaptation to the alpine environment resulted in the rapid diversification of woody forms. It is well-documented that islands and island-like systems foster the evolution of woody species from herbaceous ancestors (Darwin, 1859, p. 392; Carlquist, 1965; Lens *et al*., 2013), with this shift towards woodiness observed in numerous members of oceanic island genera such as *Aeonium* Webb & Berthel. (Mes & Hart, 1996), *Echium* Tourn. ex L. (Böhle *et al*., 1996), *Micromeria* Benth., *Sonchus* L., and *Sideritis* L. in the Canary islands (Bramwell, 1974; Lens *et al*., 2013), silverswords *Argyroxiphium* DC. (Baldwin *et al*., 1991), and *Viola* L. *(Ballard & Sytsma, 2000)* in the Hawaiian islands, as well as in several alpine radiations including New World *Lupinus* (Drummond *et al*., 2012), Arabideae DC. (Karl & Koch, 2013), Delphinieae Schrödinger (Jabbour & Renner, 2012), *Androsace* L. (Roquet *et al*., 2013), North American *Castilleja* Mutis ex L.f. (Tank & Olmstead, 2008), and Gentianinae G. Don (Favre *et al*., 2010), among others.

Various hypotheses have been proposed to explain the shift towards woodiness, or ‘insular woodiness,’ in islands and similar systems, such as the competitive advantage of woody life forms over herbaceous species (Darwin, 1859; Tilman, 1982; Givnish, 1995), increased longevity and outcrossing (Wallace, 1878; Böhle *et al*., 1996), release from seasonality allowing year-round growth (Carlquist, 1974, 2012), and the absence of large herbivores (Carlquist, 1974). However, we propose, following Hughes and Atchison (2015), that the explosive diversification of life forms in both island and alpine radiations, facilitated by shifts towards perenniality and woodiness, presents a strong case for ecological release driven by ecological opportunity (Gavrilets & Losos, 2009; Yoder *et al*., 2010). The significant variety in life forms of *Gypsophila* species and the elevated diversification rate of perennial and woody life forms suggest an explosive diversification, enabled by woodiness and perenniality and propelled by ecological opportunities following the uplift of the Irano-Anatolian and Caucasus Mountains. This diversification encompasses a range of life forms from ruderal annual herbaceous species (e.g., *G. pilosa* Huds.), to large woody forms such as subshrubs (e.g., *G. eriocalyx* Boiss.), low shrubs (e.g., *G. nabelekii* Schischkin), caespitose (e.g., *G. repens* L.), and cushion-shaped species (e.g., *G. aretioides* Boiss.). Remarkably, some cushion-shaped (pulvinate) species of *Gypsophila*, like *G. aretioides* and *G. imbricata* Rupr., have roots that can reach diameters of one meter and weigh up to 150 kg (herbarium sheet in the Leningrad Herbarium) (Barkoudah, 1962). This aligns with our finding of accelerated speciation in *Gypsophila* in montane regions, which is more than 3x the speciation rate of this genus in non-montane regions (Figure 3d; Table S4). Correspondingly, as seen in other examples of mountain radiations, such as the Andean bellflower (Lagomarsino *et al*. 2016), high-elevation habitats exhibit significantly higher speciation rates compared to low-elevation habitats (Figure 3e; Table S8).

In line with observations on other evolutionary radiations in the IT region, such as *Acantholimon* (Moharrek *et al.,* 2019), our results lead us to conclude that the evolutionary radiation within *Gypsophila* represents an instance of alpine radiation, closely linked with the diversification of habits or life forms in response to the challenging montane environments of the Irano-Anatolian and Caucasus biodiversity hotspots. This conclusion is supported by our geographic analysis of the diversification rate within *Gypsophila*, highlighting a higher speciation rate within the Irano-Anatolian and Caucasus biodiversity hotspots compared to that across the full range of the genus (Figure 3g; Table S9). The higher diversification rate at high elevations was likely due also to the lower extinction rates among species with a campanulate/turbinate shape, which is found predominantly in high-elevation species, compared to the elevated extinction rate among species with a tubiform calyx, commonly seen in annual and low-elevation *Gypsophila* species.

### The Messinian salinity crisis and the origin of Gypsophily in *Gypsophila*

Gypsophily, which can be defined as the ability of plants to tolerate and thrive in calcareous-rich soil containing high concentrations of gypsum (CaSO4·2H2O) (Moore *et al*., 2014; Escudero *et al*., 2015), represents a significant ecological adaptation that has enabled several families in Caryophyllales, such as Caryophyllaceae, Chenopodiaceae, and Plumbaginaceae, to colonize gypsum-rich soils in habitats of southwestern Asia (Moore *et al*., 2014; Escudero *et al*., 2015). These regions, along with the Mediterranean, the Horn of Africa, and southwestern North America, are home to some of the largest gypsum surfaces and host a diverse array of gypsum-endemic plant taxa (Moore *et al*., 2014; Escudero *et al*., 2015). Some authors suggest that lineages within Caryophyllales are ‘preadapted’ to cope with the challenges of gypsum soils (Duvigneaud & Denaeyer-De Smet, 1973), for instance, through the accumulation of organic sulfur compounds as a preadaptation to herbivory (Parsons, 1976). However, we propose that the widespread occurrence of gypsophily in the flora of southwestern Asia and the Mediterranean is rooted in paleoenvironmental changes and crises that have transformed this region into one of the driest and hottest in the world, with extensive gypsum surfaces. This includes dramatic tectonic activities that brought an end to the ancient Paratethys, a complex network of inland seaways, brackish lakes, and wetlands, along with other ancient bodies of water that arose from the Tethys Ocean during the Oligocene. The remnants of the Tethys Ocean, which maintained the connection between the Indo-Tethys and Mediterranean-Tethys such as the expansive Paratethys, the Mediterranean, Carpathian, Qom, and Arabian basins, as well as the southern Tethyan seaways and Bitlis seaway, transformed into multiple evaporative basins or shallow seaways and then completely closed during the Miocene (Báldi 1980; Rögl 1997; Sun *et al*. 2021; Palcu *et al*. 2023). These transformations, which connected Afro-Arabia to Eurasia (Rögl, 1999), coincided with the first animal migrations over the ‘Gomphotherium Landbridge’ in the early Miocene (Harzhauser *et al*., 2007). Not only did these transformations alter the global oceanic circulation pattern and influence the climate of the entire planet (Straume *et al*., 2020), but they also led to the creation of the West Asian landmass, which includes the habitats of the Irano-Anatolian and Caucasus biodiversity hotspots (Rögl, 1999), and resulted in the deposition of vast evaporative deposits, thereby creating one of the planet’s largest areas of surface gypsum.

Our ancestral biogeographic reconstruction suggests that the Mediterranean region and western Asia served as the ancestral range for *Gypsophila* during the late Miocene and early Pliocene. This period was heavily impacted by the Messinian Salinity Crisis (MSC), a series of cycles of partial or nearly complete desiccation of the Mediterranean Sea from 5.96 to 5.33 million years ago, caused by reduced water inflow from the Atlantic Ocean (Hsü *et al*., 1973, 1977). This event was likely due to competing tectonics and erosion at the Gibraltar arc (Garcia-Castellanos & Villaseñor, 2011), leading to massive deposition of salt and gypsum across the Mediterranean region (Hsü *et al*., 1977; Krijgsman *et al*., 1999). The MSC may have acted as an evolutionary catalyst, fostering gypsophily in the common ancestor of *Gypsophila*. The subsequent advent and diversification of *Gypsophila* in these regions, followed by its dispersion to the young high-altitude habitats of western Asia and the Caucasus, are likely tied to the species’ acquisition of limestone-dwelling abilities during the MSC. As tolerance to toxic concentrations of heavy metals and a low Ca:Mg ratio in serpentine habitats (Brooks, 1987) may also be linked to the gypsophily syndrome (Escudero *et al*., 2015). This capability may have facilitated the geographic expansion and rapid dispersal of ancestral *Gypsophila* populations from their origins in West Asia and the Mediterranean region to gypsum-rich habitats of the Irano-Turanian regions and Western Mediterranean and Europe. This would have been followed by niche shift and allopatric speciation (de Luis *et al*., 2018), intertwined with other historical phenomena such as tectonic activities, orogeny, and Quaternary climate oscillations, leading to the rapid radiation of *Gypsophila* into more than 150 species and perhaps hundreds of subspecies, varieties, and demes. This shift in the diversification rate of *Gypsophila* coincides with other examples of evolutionary radiation in the Mediterranean and Western Asia, such as *Limonium* (Lledó *et al*., 2005), *Cistus* (Guzmán *et al*., 2009), *Cousinia* (López-Vinyallonga *et al*., 2009), *Dianthus* (Valente *et al*., 2010)*, Onosma* (Cecchi *et al*., 2011), *Tragopogon* (Bell *et al*., 2012), and *Knautia* sect. *Trichera* (Frajman *et al*., 2016). In these genera, diversification was likely triggered by climatic and topographic changes in the Mediterranean following the MSC in the late Miocene; this period marked a transition from the Miocene’s warm and moist climate to a distinct seasonal pattern characterized by dry summers and cold, wet winters (Valente *et al*., 2010; Bell *et al*., 2012; Frajman *et al*., 2016).

In conclusion, our study provides evidence for one of the most rapid species radiations on record and valuable insights into the role of paleogeological and paleoenvironmental events in evolutionary dynamics and in shaping patterns of biodiversity. Our results underscore the impact of the recent uplift of mountain systems in the Irano-Anatolian and Caucasus biodiversity hotspots and subsequent global cooling on the spectacular radiation and spread of the alpine-specialist genus, *Gypsophila*. Our study also highlights the ecological and morphological correlates of diversification, emphasizing the adaptation of *Gypsophila* species to alpine conditions through shifts towards perenniality and woodiness. Our findings on the connection between gypsophily and the Messinian Salinity Crisis, followed by the dispersion and niche diversification of *Gypsophila* in gypsum-rich habitats, suggest a complex interplay between environmental changes and evolutionary adaptations. Further research is needed to investigate the specific environmental factors and underlying genomics architecture that have driven the diversification of *Gypsophila* and to explore its potential role as a model system for understanding the evolution of plant diversity in these biodiversity hotspots. Additionally, testing these diversification correlates on other evolutionary radiations in the Irano-Anatolian and Caucasus biodiversity hotspots will help determine the relative importance of paleogeological-environmental events versus morphological innovations in shaping biodiversity within these regions.

## Acknowledgments

Our sincere appreciation goes to the curators of the herbaria B, G, LE, M, MSB, TUH, W, and WU for their invaluable support and for providing us the opportunity to examine specimens of this plant group and to collect samples for DNA extraction.

## Author contributions

HM: Specimen study, plant collection, laboratory procedures, phylogenetic and statistical analyses, manuscript preparation. RR: Nomenclatural research, manuscript revision, comments on phylogenetic trees. GH: Providing laboratory and technical facilities, providing some plant materials. ES: Manuscript revision. NM: plant collection, manuscript preparation, statistical analyses. SZ: Specimen study, plant collection, manuscript revision, providing sequences. — HM; RR; GH; ES; NM; SZ)

## Notes

### Competing Interest Statement

The authors have declared no competing interest.

### Summary of Updates

In this version, the entire manuscript has been refined and improved, with the discussion section expanded in particular.

## References

Allen MB, Armstrong HA. 2008. Arabia–Eurasia collision and the forcing of mid-Cenozoic global cooling. Palaeogeography, palaeoclimatology, palaeoecology 265: 52–58.

Antonelli A, Kissling WD, Flantua SGA, Bermúdez MA, Mulch A, Muellner-Riehl AN, Kreft H, Linder HP, Badgley C, Fjeldså J, et al. 2018. Geological and climatic influences on mountain biodiversity. Nature geoscience 11: 718–725.

Antonelli A, Nylander JAA, Persson C, Sanmartín I. 2009. Tracing the impact of the Andean uplift on Neotropical plant evolution. Proceedings of the National Academy of Sciences of the United States of America 106: 9749–9754.

Báldi T. 1980. The early history of the Paratethys. Bulletin of the Hungarian Geological Society.

Baldwin BG, Kyhos DW, Dvorak J, Carr GD. 1991. Chloroplast DNA evidence for a North American origin of the Hawaiian silversword alliance (Asteraceae). Proceedings of the National Academy of Sciences of the United States of America 88: 1840–1843.

Baldwin BG, Sanderson MJ. 1998. Age and rate of diversification of the Hawaiian silversword alliance (Compositae). Proceedings of the National Academy of Sciences of the United States of America 95: 9402–9406.

Ballard HE Jr, Sytsma KJ. 2000. Evolution and biogeography of the woody Hawaiian violets (Viola, Violaceae): arctic origins, herbaceous ancestry and bird dispersal. Evolution; international journal of organic evolution 54: 1521–1532.

Ballato P, Mulch A, Landgraf A, Strecker MR, Dalconi MC, Friedrich A, Tabatabaei SH. 2010. Middle to late Miocene Middle Eastern climate from stable oxygen and carbon isotope data, southern Alborz mountains, N Iran. Earth and planetary science letters 300: 125–138.

Barkoudah. 1962. Sion of Gypsophyla, Bolanthus, Ankyropetalum and Phryna. Wentia.

Bell MA, Lloyd GT. 2015. strap: an R package for plotting phylogenies against stratigraphy and assessing their stratigraphic congruence. Palaeontology 58: 379–389.

Bell CD, Mavrodiev EV, Soltis PS, Calaminus AK, Albach DC, Cellinese N, Garcia-Jacas N, Soltis DE. 2012. Rapid diversification of Tragopogon and ecological associates in Eurasia. Journal of evolutionary biology 25: 2470–2480.

Böhle UR, Hilger HH, Martin WF. 1996. Island colonization and evolution of the insular woody habit in Echium L. (Boraginaceae). Proceedings of the National Academy of Sciences of the United States of America 93: 11740–11745.

Bramwell D. 1974. Wild flowers of the Canary Islands. (No Title).

Brooks R. 1987. Serpentine and its vegetation. A multidisciplinary approach.

Brown GK, Nelson G, Ladiges PY. 2006. Historical biogeography of Rhododendron section Vireya and the Malesian Archipelago. Journal of biogeography 33: 1929–1944.

Carlquist SJ. 1965. Island life: a natural history of the islands of the world. (No Title).

Carlquist S. 1974. Island biology–Columbia University Press. New York, USA.

Carlquist S. 2012. How wood evolves: a new synthesis. Botany 90: 901–940.

Cecchi L, Coppi A, Selvi F. 2011. Evolutionary dynamics of serpentine adaptation in Onosma (Boraginaceae) as revealed by ITS sequence data. Osterreichische botanische Zeitschrift 297: 185–199.

Cohen KM, Finney SC, Gibbard PL, Fan J-X. 2013. The ICS international chronostratigraphic chart. Episodes 36: 199–204.

Comes HP, Kadereit JW. 2003. Spatial and temporal patterns in the evolution of the flora of the European Alpine System. Taxon 52: 451–462.

Condamine FL, Rolland J, Morlon H. 2013. Macroevolutionary perspectives to environmental change. Ecology letters 16 Suppl 1: 72–85.

Condamine FL, Rolland J, Morlon H. 2019. Assessing the causes of diversification slowdowns: temperature-dependent and diversity-dependent models receive equivalent support. Ecology letters 22: 1900–1912.

van Dam JA. 2006. Geographic and temporal patterns in the late Neogene (12–3 Ma) aridification of Europe: The use of small mammals as paleoprecipitation proxies. Palaeogeography, palaeoclimatology, palaeoecology 238: 190–218.

Darriba D, Taboada GL, Doallo R, Posada D. 2012. jModelTest 2: more models, new heuristics and parallel computing. Nature methods 9: 772.

Darwin CR. 1859. The Origin ofThe Species by Means of Natural Selection.

Davis SD, Heywood VH, Hamilton AC. 1994. Centres of Plant Diversity: Asia, Australasia, and the Pacific. World Conservation Union.

Dilek Y, Imamverdiyev N, Altunkaynak Ş. 2010. Geochemistry and tectonics of Cenozoic volcanism in the Lesser Caucasus (Azerbaijan) and the peri-Arabian region: collision-induced mantle dynamics and its magmatic fingerprint. International geology review 52: 536–578.

Djamali M, Baumel A, Brewer S, Jackson ST, Kadereit JW, López-Vinyallonga S, Mehregan I, Shabanian E, Simakova A. 2012a. Ecological implications of Cousinia Cass. (Asteraceae) persistence through the last two glacial–interglacial cycles in the continental Middle East for the Irano-Turanian flora. Review of palaeobotany and palynology 172: 10–20.

Djamali M, Brewer S, Breckle SW, Jackson ST. 2012b. Climatic determinism in phytogeographic regionalization: A test from the Irano-Turanian region, SW and Central Asia. *Flora - Morphology, Distribution*, Functional Ecology of Plants 207: 237–249.

Drummond CS, Eastwood RJ, Miotto STS, Hughes CE. 2012. Multiple continental radiations and correlates of diversification in Lupinus (Leguminosae): testing for key innovation with incomplete taxon sampling. Systematic biology 61: 443–460.

Drummond AJ, Ho SYW, Rawlence N, Rambaut A. 2007. A rough guide to BEAST 1.4.

Drummond AJ, Rambaut A. 2007. BEAST: Bayesian evolutionary analysis by sampling trees. BMC evolutionary biology 7: 214.

Duvigneaud P, Denaeyer-De Smet S. 1973. Considérations sur l’écologie de la nutrition minérale des tapis végétaux naturels. Oecologia Plantarum 8: 219–246.

Escudero A, Palacio S, Maestre FT, Luzuriaga AL. 2015. Plant life on gypsum: a review of its multiple facets. Biological reviews of the Cambridge Philosophical Society 90: 1–18.

Fassou G, Korotkova N, Nersesyan A, Koch MA, Dimopoulos P, Borsch T. 2022. Taxonomy of Dianthus (Caryophyllaceae) - overall phylogenetic relationships and assessment of species diversity based on a first comprehensive checklist of the genus. PhytoKeys 196: 91–214.

Favre A, Päckert M, Pauls SU, Jähnig SC, Uhl D, Michalak I, Muellner-Riehl AN. 2015. The role of the uplift of the Qinghai-Tibetan Plateau for the evolution of Tibetan biotas. Biological reviews of the Cambridge Philosophical Society 90: 236–253.

Favre A, Yuan Y-M, Küpfer P, Alvarez N. 2010. Phylogeny of subtribe Gentianinae (Gentianaceae): Biogeographic inferences despite limitations in temporal calibration points. Taxon 59: 1701–1711.

FitzJohn RG. 2012. Diversitree: comparative phylogenetic analyses of diversification in R. Methods in ecology and evolution / British Ecological Society.

Fonseca CR. 2009. The silent mass extinction of insect herbivores in biodiversity hotspots. Conservation biology: the journal of the Society for Conservation Biology 23: 1507–1515.

Frajman B, Eggens F, Oxelman B. 2009. Hybrid origins and homoploid reticulate evolution within Heliosperma (Sileneae, Caryophyllaceae)--a multigene phylogenetic approach with relative dating. Systematic biology 58: 328–345.

Frajman B, Rešetnik I, Niketić M, Ehrendorfer F, Schönswetter P. 2016. Patterns of rapid diversification in heteroploid Knautia sect. Trichera (Caprifoliaceae, Dipsacoideae), one of the most intricate taxa of the European flora. BMC evolutionary biology 16: 204.

Garcia-Castellanos D, Villaseñor A. 2011. Messinian salinity crisis regulated by competing tectonics and erosion at the Gibraltar arc. Nature 480: 359–363.

Gavillot Y, Axen GJ, Stockli DF, Horton BK, Fakhari MD. 2010. Timing of thrust activity in the High Zagros fold-thrust belt, Iran, from (U-Th)/He thermochronometry. Tectonics 29.

Gavrilets S, Losos JB. 2009. Adaptive radiation: contrasting theory with data. Science 323: 732–737.

Gehrke B, Linder HP. 2009. The scramble for Africa: pan-temperate elements on the African high mountains. Proceedings. Biological sciences / The Royal Society 276: 2657–2665.

Ghane-Ameleh S, Khosravi M, Saberi-Pirooz R, Ebrahimi E, Aghbolaghi MA, Ahmadzadeh F. 2021. Mid-Pleistocene Transition as a trigger for diversification in the Irano-Anatolian region: Evidence revealed by phylogeography and distribution pattern of the eastern three-lined lizard. Global Ecology and Conservation 31: e01839.

Gillespie RG, Roderick GK. 2014. Evolution: Geology and climate drive diversification. Nature 509: 297–298.

Givnish TJ. 1995. 1 - Plant Stems: Biomechanical Adaptation for Energy Capture and Influence on Species Distributions. In: Gartner BL, ed. Plant Stems. San Diego: Academic Press, 3–49.

Goldberg EE, Lancaster LT, Ree RH. 2011. Phylogenetic inference of reciprocal effects between geographic range evolution and diversification. Systematic biology 60: 451–465.

Greenberg AK, Donoghue MJ. 2011. Molecular systematics and character evolution in Caryophyllaceae. Taxon 60: 1637–1652.

Guzmán B, Lledó MD, Vargas P. 2009. Adaptive radiation in mediterranean cistus (cistaceae). PloS one 4: e6362.

Harmon LJ, Weir JT, Brock CD, Glor RE, Challenger W. 2008. GEIGER: investigating evolutionary radiations. Bioinformatics 24: 129–131.

Harzhauser M, Kroh A, Mandic O, Piller WE, Göhlich U, Reuter M, Berning B. 2007. Biogeographic responses to geodynamics: A key study all around the Oligo–Miocene Tethyan Seaway. Zoologischer Anzeiger - A Journal of Comparative Zoology 246: 241–256.

Hatzfeld D, Molnar P. 2010. Comparisons of the kinematics and deep structures of the Zagros and Himalaya and of the Iranian and Tibetan plateaus and geodynamic implications. Reviews of geophysics 48.

Hernández-Ledesma P, Berendsohn WG, Borsch T, Von Mering S, Akhani H, Arias S, Castañeda-Noa I, Eggli U, Eriksson R, Flores-Olvera H, et al. 2015. A taxonomic backbone for the global synthesis of species diversity in the angiosperm order Caryophyllales. Willdenowia 45: 281–383.

Ho SYW, Phillips MJ. 2009. Accounting for calibration uncertainty in phylogenetic estimation of evolutionary divergence times. Systematic biology 58: 367–380.

Hsü KJ, Montadert L, Bernoulli D, Cita MB, Erickson A, Garrison RE, Kidd RB, Mèlierés F, Müller C, Wright R. 1977. History of the Mediterranean salinity crisis. Nature 267: 399–403.

Hsü KJ, Ryan WBF, Cita MB. 1973. Late Miocene Desiccation of the Mediterranean. Nature 242: 240– 244.

Huber-Morath. A. 1967. Gypsophila Reeve H, Davis DL. 1967. Flora of Turkey and the East Aegean Islands, Vol. 2. Edinburgh University Press.

Hughes CE, Atchison GW. 2015. The ubiquity of alpine plant radiations: from the Andes to the Hengduan Mountains. The New phytologist 207: 275–282.

Hughes C, Eastwood R. 2006. Island radiation on a continental scale: exceptional rates of plant diversification after uplift of the Andes. Proceedings of the National Academy of Sciences of the United States of America 103: 10334–10339.

Jabbour F, Renner SS. 2012. A phylogeny of Delphinieae (Ranunculaceae) shows that Aconitum is nested within Delphinium and that Late Miocene transitions to long life cycles in the Himalayas and Southwest China coincide with bursts in diversification. Molecular phylogenetics and evolution 62: 928– 942.

Joly S, Heenan PB, Lockhart PJ. 2014. Species radiation by niche shifts in New Zealand’s rockcresses (Pachycladon, Brassicaceae). Systematic biology 63: 192–202.

Jordan GJ, Macphail MK. 2003. A Middle-Late Eocene inflorescence of Caryophyllaceae from Tasmania, Australia. American journal of botany 90: 761–768.

Kadereit JW, Griebeler EM, Comes HP. 2004. Quaternary diversification in European alpine plants: pattern and process. Philosophical transactions of the Royal Society of London. Series B, Biological sciences 359: 265–274.

Karl R, Koch MA. 2013. A world-wide perspective on crucifer speciation and evolution: phylogenetics, biogeography and trait evolution in tribe Arabideae. Annals of botany 112: 983–1001.

Katoh K, Standley DM. 2013. MAFFT multiple sequence alignment software version 7: improvements in performance and usability. Molecular biology and evolution 30: 772–780.

Kearse M, Moir R, Wilson A, Stones-Havas S, Cheung M, Sturrock S, Buxton S, Cooper A, Markowitz S, Duran C, et al. 2012. Geneious Basic: an integrated and extendable desktop software platform for the organization and analysis of sequence data. Bioinformatics 28: 1647–1649.

Keskin M. 2003. Magma generation by slab steepening and breakoff beneath a subduction-accretion complex: An alternative model for collision-related volcanism in Eastern Anatolia, Turkey. Geophysical research letters 30.

Knox EB, Palmer JD. 1995. Chloroplast DNA variation and the recent radiation of the giant senecios (Asteraceae) on the tall mountains of eastern Africa. Proceedings of the National Academy of Sciences of the United States of America 92: 10349–10353.

Krijgsman W, Hilgen FJ, Raffi I, Sierro FJ, Wilson DS. 1999. Chronology, causes and progression of the Messinian salinity crisis. Nature 400: 652–655.

Lagomarsino LP, Condamine FL, Antonelli A, Mulch A, Davis CC. 2016. The abiotic and biotic drivers of rapid diversification in A ndean bellflowers (Campanulaceae). The New phytologist 210: 1430– 1442.

Lens F, Davin N, Smets E, del Arco M. 2013. Insular Woodiness on the Canary Islands: A Remarkable Case of Convergent Evolution. International journal of plant sciences 174: 992–1013.

Le Roux JJ, Hui C, Castillo ML, Iriondo JM, Keet J-H, Khapugin AA, Médail F, Rejmánek M, Theron G, Yannelli FA, et al. 2019. Recent Anthropogenic Plant Extinctions Differ in Biodiversity Hotspots and Coldspots. Current biology: CB 29: 2912–2918.e2.

Linder HP. 2014. The evolution of African plant diversity. Frontiers in Ecology and Evolution 2.

Liu J-Q, Wang Y-J, Wang A-L, Hideaki O, Abbott RJ. 2006. Radiation and diversification within the Ligularia–Cremanthodium–Parasenecio complex (Asteraceae) triggered by uplift of the Qinghai-Tibetan Plateau. Molecular phylogenetics and evolution 38: 31–49.

Lledó MD, Crespo MB, Fay MF, Chase MW. 2005. Molecular phylogenetics of Limonium and related genera (Plumbaginaceae): biogeographical and systematic implications. American journal of botany 92: 1189–1198.

López-Vinyallonga S, Mehregan I, Garcia-Jacas N, Tscherneva O, Susanna A, Kadereit JW. 2009. Phylogeny and evolution of the Arctium-Cousinia complex (Compositae, Cardueae-Carduinae). Taxon 58: 153–171.

de Luis M, Bartolomé C, García Cardo Ó, Martínez Labarga JM, Álvarez-Jiménez J. 2018. Sympatric and allopatric niche shift of endemic Gypsophila (Caryophyllaceae) taxa in the Iberian Peninsula. PloS one 13: e0206043.

Maddison WP, Maddison DR. 2021.Mesquite: a modular system for evolutionary analysis. Version 3.31. 2017.

Maddison WP, Midford PE, Otto SP. 2007. Estimating a binary character’s effect on speciation and extinction. Systematic biology 56: 701–710.

Madhani H, Rabeler R, Pirani A, Oxelman B, Heubl G, Zarre S. 2018. Untangling phylogenetic patterns and taxonomic confusion in tribe Caryophylleae (Caryophyllaceae) with special focus on generic boundaries. Taxon 67: 83–112.

Madhani H, Rabeler RK, Zarre S. 2024. Generic delimitation of *Bolanthus* and resurrection of *Jordania* within Caryophylleae (Caryophyllaceae). Taxon.

Madriñán S, Cortés AJ, Richardson JE. 2013. Páramo is the world’s fastest evolving and coolest biodiversity hotspot. Frontiers in genetics 4: 192.

Magallón S, Sanderson MJ. 2001. Absolute diversification rates in angiosperm clades. Evolution; international journal of organic evolution 55: 1762–1780.

Mahmoudi Shamsabad M, Moharrek F, Assadi M, Nieto Feliner G. 2021. Biogeographic history and diversification patterns in the Irano-Turanian genus Acanthophyllum s.l. (Caryophyllaceae). Plant biosystems 155: 425–435.

Malcolm JR, Liu C, Neilson RP, Hansen L, Hannah L. 2006. Global warming and extinctions of endemic species from biodiversity hotspots. Conservation biology: the journal of the Society for Conservation Biology 20: 538–548.

Manafzadeh S, Salvo G, Conti E. 2014. A tale of migrations from east to west: the Irano-Turanian floristic region as a source of Mediterranean xerophytes. Journal of biogeography 41: 366–379.

Manafzadeh S, Staedler YM, Conti E. 2017. Visions of the past and dreams of the future in the Orient: the Irano-Turanian region from classical botany to evolutionary studies. Biological reviews of the Cambridge Philosophical Society 92: 1365–1388.

Matzke NJ. 2014. Model selection in historical biogeography reveals that founder-event speciation is a crucial process in Island Clades. Systematic biology 63: 951–970.

Mes THM, Hart HT. 1996. The evolution of growth-forms in the Macaronesian genus Aeonium (Crassulaceae) inferred from chloroplast DNA RFLPs and morphology. Molecular ecology 5: 351–363.

Meulenkamp JE, Sissingh W. 2003. Tertiary palaeogeography and tectonostratigraphic evolution of the Northern and Southern Peri-Tethys platforms and the intermediate domains of the African–Eurasian convergent plate boundary zone. Palaeogeography, palaeoclimatology, palaeoecology 196: 209–228.

Miller MA, Pfeiffer W, Schwartz T. 2010. Proceedings of the gateway computing environments workshop (GCE). Creating the CIPRES science.

Mitchell JS, Rabosky DL. 2017. Bayesian model selection with BAMM: effects of the model prior on the inferred number of diversification shifts. Methods in ecology and evolution / British Ecological Society 8: 37–46.

Mittermeier RA, Turner WR, Larsen FW, Brooks TM, Gascon C. 2011. Global Biodiversity Conservation: The Critical Role of Hotspots. In: Zachos FE, Habel JC, eds. Biodiversity Hotspots: Distribution and Protection of Conservation Priority Areas. Berlin, Heidelberg: Springer Berlin Heidelberg, 3–22.

Moharrek F, Sanmartín I, Kazempour-Osaloo S, Nieto Feliner G. 2019. Morphological Innovations and Vast Extensions of Mountain Habitats Triggered Rapid Diversification Within the Species-Rich Irano-Turanian Genus Acantholimon (Plumbaginaceae). Frontiers in genetics 9: 698.

Moore MJ, Mota JF, Douglas NA, Flores-Olvera H, Ochoterena H. 2014. The ecology, assembly, and evolution of gypsophile floras.

Morlon H, Lewitus E, Condamine FL, Manceau M, Clavel J, Drury J. 2016. RPANDA: an R package for macroevolutionary analyses on phylogenetic trees. Methods in ecology and evolution / British Ecological Society 7: 589–597.

Morlon H, Parsons TL, Plotkin JB. 2011. Reconciling molecular phylogenies with the fossil record. Proceedings of the National Academy of Sciences of the United States of America 108: 16327–16332.

Mouthereau F, Lacombe O, Vergés J. 2012. Building the Zagros collisional orogen: Timing, strain distribution and the dynamics of Arabia/Eurasia plate convergence. Tectonophysics 532–535: 27–60.

Naciri Y, Linder HP. 2020. The genetics of evolutionary radiations. Biological reviews of the Cambridge Philosophical Society 95: 1055–1072.

Noroozi J, Akhani H, Breckle S-W. 2008. Biodiversity and phytogeography of the alpine flora of Iran. Biodiversity and conservation 17: 493–521.

Noroozi J, Pirani A, Moazzeni H, Mahmoodi M, Zare G, Noormohammadi A, Barfuss MHJ, Suen M, Schneeweiss GM. 2020. The new locally endemic genus Yazdana (Caryophyllaceae) and patterns of endemism highlight the high conservation priority of the poorly studied Shirkuh Mountains (central Iran). Journal of systematics and evolution 58: 339–353.

Noroozi J, Talebi A, Doostmohammadi M, Rumpf SB, Linder HP, Schneeweiss GM. 2018. Hotspots within a global biodiversity hotspot - areas of endemism are associated with high mountain ranges. Scientific reports 8: 10345.

Noss RF, Platt WJ, Sorrie BA, Weakley AS, Means DB, Costanza J, Peet RK. 2015. How global biodiversity hotspots may go unrecognized: lessons from the North American Coastal Plain. Diversity & distributions 21: 236–244.

Nürk N, Scheriau C, Madriñán S. 2013. Explosive radiation in high Andean Hypericum—rates of diversification among New World lineages. Frontiers in genetics 4.

Palcu DV, Mariș I, de Leeuw A, Melinte-Dobrinescu M, Anton E, Frunzescu D, Popov S, Stoica M, Jovane L, Krijgsman W. 2023. The legacy of the Tethys Ocean: Anoxic seas, evaporitic basins, and megalakes in the Cenozoic of Central Europe. Earth-Science Reviews:104594.

Parolly G. 2015. The high-mountain flora and vegetation of the western and central Taurus mts. (turkey) in the times of climate change. In: Climate Change Impacts on High-Altitude Ecosystems. Cham: Springer International Publishing, 99–133.

Parsons RF. 1976. Gypsophily in Plants-A Review. The American midland naturalist 96: 1–20.

Pearce JA, Bender JF, De Long SE, Kidd WSF, Low PJ, Güner Y, Saroglu F, Yilmaz Y, Moorbath S, Mitchell JG. 1990. Genesis of collision volcanism in Eastern Anatolia, Turkey. Journal of Volcanology and Geothermal Research 44: 189–229.

Pérez-Escobar OA, Chomicki G, Condamine FL, Karremans AP, Bogarín D, Matzke NJ, Silvestro D, Antonelli A. 2017. Recent origin and rapid speciation of Neotropical orchids in the world’s richest plant biodiversity hotspot. The New phytologist 215: 891–905.

Pirani A, Moazzeni H, Zarre S, Rabeler RK, Oxelman B, Pavlenko AV, Kovalchuk A. 2020. Phylogeny ofAcanthophyllums.l. revisited: An update on generic concept and sectional classification. Taxon 69: 500–514.

Pirani A, Zarre S, Pfeil BE, Bertrand YJK, Assadi M, Oxelman B. 2014. Molecular phylogeny of Acanthophyllum (Caryophyllaceae: Caryophylleae), with emphasis on infrageneric classification. Taxon 63: 592–607.

Plummer M, Best N, Cowles K, Vines K. 2006. CODA: convergence diagnosis and output analysis for MCMC. R News 6: 7–11.

Popov SV, Shcherba IG, Ilyina LB, Nevesskaya LA, Paramonova NP, Khondkarian SO, Magyar I. 2006. Late Miocene to Pliocene palaeogeography of the Paratethys and its relation to the Mediterranean. Palaeogeography, palaeoclimatology, palaeoecology 238: 91–106.

Rabosky DL. 2014. Automatic detection of key innovations, rate shifts, and diversity-dependence on phylogenetic trees. PloS one 9: e89543.

Rabosky DL, Grundler M, Anderson C. 2014. BAMM tools: an R package for the analysis of evolutionary dynamics on phylogenetic trees. Methods in ecology and evolution / British Ecological Society.

Rahbek C, Borregaard MK, Antonelli A, Colwell RK, Holt BG, Nogues-Bravo D, Rasmussen CMØ, Richardson K, Rosing MT, Whittaker RJ, et al. 2019. Building mountain biodiversity: Geological and evolutionary processes. Science 365: 1114–1119.

Rambaut A, Drummond AJ, Xie D, Baele G, Suchard MA. 2018. Posterior Summarization in Bayesian Phylogenetics Using Tracer 1.7. Systematic biology 67: 901–904.

Ramsey, Ripley. 2013. pspline: Penalized smoothing splines. R package version.

Rechinger. 1988. Flora Iranica: Caryophyllaceae, vol. 163. Graz: Akademische Druck-u. Verlagsanstalt.

Ree RH, Smith SA. 2008. Maximum likelihood inference of geographic range evolution by dispersal, local extinction, and cladogenesis. Systematic biology 57: 4–14.

Rögl F. 1997. Palaeogeographic Considerations for Mediterranean and Paratethys Seaways (Oligocene to Miocene). Annalen des Naturhistorischen Museums in Wien. Serie A. Mineralogie und Petrographie, Geologie und Palaeontologie, Anthropologie und Praehistorie 99: 279–310.

Rögl F. 1999. Mediterranean and Paratethys. Facts and hypotheses of an Oligocene to Miocene paleogeography (short overview).

Ronquist F. 2004. Bayesian inference of character evolution. Trends in ecology & evolution 19: 475–481.

Ronquist F, Huelsenbeck JP. 2003. MrBayes 3: Bayesian phylogenetic inference under mixed models. Bioinformatics 19: 1572–1574.

Roquet C, Boucher FC, Thuiller W, Lavergne S. 2013. Replicated radiations of the alpine genus Androsace (Primulaceae) driven by range expansion and convergent key innovations. Journal of biogeography 40: 1874–1886.

Schenk JJ. 2016. Consequences of Secondary Calibrations on Divergence Time Estimates. PloS one 11: e0148228.

Şengör AMC, Görür N, Şaroğlu F. 1985.Strike-Slip Faulting and Related Basin Formation in Zones of Tectonic Escape: Turkey as a Case Study.

Şengör AMC, Kidd WSF. 1979. Post-collisional tectonics of the Turkish-Iranian plateau and a comparison with Tibet. Tectonophysics 55: 361–376.

Şengör AMC, Yilmaz Y. 1981. Tethyan evolution of Turkey: A plate tectonic approach. Tectonophysics 75: 181–241.

Simões M, Breitkreuz L, Alvarado M, Baca S, Cooper JC, Heins L, Herzog K, Lieberman BS. 2016. The Evolving Theory of Evolutionary Radiations. Trends in ecology & evolution 31: 27–34.

Stamatakis A. 2014. RAxML version 8: a tool for phylogenetic analysis and post-analysis of large phylogenies. Bioinformatics 30: 1312–1313.

Steinbauer MJ, Field R, Grytnes J-A, Trigas P, Ah-Peng C, Attorre F, Birks HJB, Borges PAV, Cardoso P, Chou C-H, et al. 2016. Topography-driven isolation, speciation and a global increase of endemism with elevation. Global ecology and biogeography: a journal of macroecology 25: 1097–1107.

Straume EO, Gaina C, Medvedev S, Nisancioglu KH. 2020. Global Cenozoic Paleobathymetry with a focus on the Northern Hemisphere Oceanic Gateways. Gondwana Research 86: 126–143.

Suchard MA, Lemey P, Baele G, Ayres DL, Drummond AJ, Rambaut A. 2018. Bayesian phylogenetic and phylodynamic data integration using BEAST 1.10. Virus evolution 4: vey016.

Sun J, Sheykh M, Ahmadi N, Cao M, Zhang Z, Tian S, Sha J, Jian Z, Windley BF, Talebian M. 2021. Permanent closure of the Tethyan Seaway in the northwestern Iranian Plateau driven by cyclic sea-level fluctuations in the late Middle Miocene. Palaeogeography, Palaeoclimatology, Palaeoecology 564:110172.

Sun M, Folk RA, Gitzendanner MA, Soltis PS, Chen Z, Soltis DE, Guralnick RP. 2020. Recent accelerated diversification in rosids occurred outside the tropics. Nature communications 11: 3333.

Sun Y, Wang A, Wan D, Wang Q, Liu J. 2012. Rapid radiation of Rheum (Polygonaceae) and parallel evolution of morphological traits. Molecular phylogenetics and evolution 63: 150–158.

Swofford DL. 1993.PAUP: Phylogenetic analysis using parsimony, ver. 3.1. 1. Illinois Natural History Survey, Champaign.

Takhtadzhian AL, Takhtadzhian LA, Takhtajan A, Crovello TJ. 1986. Floristic regions of the world. University of California press.

Tank DC, Olmstead RG. 2008. From annuals to perennials: phylogeny of subtribe Castillejinae (Orobanchaceae). American journal of botany 95: 608–625.

Thompson JB, Davis KE, Dodd HO, Wills MA, Priest NK. 2023. Speciation across the Earth driven by global cooling in terrestrial orchids. Proceedings of the National Academy of Sciences of the United States of America 120: e2102408120.

Thiers BM. 2023. Index Herbariorum. https://sweetgum.nybg.org/science/ih/

Tilman D. 1982. Resource competition and community structure. Monographs in population biology 17: 1–296.

Valente LM, Savolainen V, Vargas P. 2010. Unparalleled rates of species diversification in Europe. Proceedings. Biological sciences / The Royal Society 277: 1489–1496.

Vincent SJ, Morton AC, Carter A, Gibbs S, Barabadze TG. 2007. Oligocene uplift of the Western Greater Caucasus: an effect of initial Arabia?Eurasia collision. Terra nova 19: 160–166.

Wallace AR. 1878. Tropical Nature, and Other Essays. Macmillan and Company.

Wen J, Zhang J-Q, Nie Z-L, Zhong Y, Sun H. 2014. Evolutionary diversifications of plants on the Qinghai-Tibetan Plateau. Frontiers in genetics 5: 4.

Wiens JJ. 2017. What explains patterns of biodiversity across the Tree of Life?: New research is revealing the causes of the dramatic variation in species numbers across branches of the Tree of Life. BioEssays: news and reviews in molecular, cellular and developmental biology 39.

Winkworth RC, Wagstaff SJ, Glenny D, Lockhart PJ. 2005. Evolution of the New Zealand mountain flora: Origins, diversification and dispersal. Organisms, diversity & evolution 5: 237–247.

Wolfe AD, Randle CP, Datwyler SL, Morawetz JJ, Arguedas N, Diaz J. 2006. Phylogeny, taxonomic affinities, and biogeography of Penstemon (Plantaginaceae) based on ITS and cpDNA sequence data. American journal of botany 93: 1699–1713.

Xing Y, Ree RH. 2017. Uplift-driven diversification in the Hengduan Mountains, a temperate biodiversity hotspot. Proceedings of the National Academy of Sciences of the United States of America 114: E3444–E3451.

Xue B, Song Z, Cai J, Ma Z, Huang J, Li Y, Yao G. 2023. Phylogenetic analysis and temporal diversification of the tribe Alsineae (Caryophyllaceae) with the description of three new genera, Hesperostellaria, Reniostellaria and Torreyostellaria. Frontiers in plant science 14: 1127443.

Yin A. 2010. Cenozoic tectonic evolution of Asia: A preliminary synthesis. Tectonophysics 488: 293– 325.

Yoder JB, Clancey E, Des Roches S, Eastman JM, Gentry L, Godsoe W, Hagey TJ, Jochimsen D, Oswald BP, Robertson J, et al. 2010. Ecological opportunity and the origin of adaptive radiations. Journal of evolutionary biology 23: 1581–1596.

Yu Y, Blair C, He X. 2020. RASP 4: Ancestral State Reconstruction Tool for Multiple Genes and Characters. Molecular biology and evolution 37: 604–606.

Yu Y, Harris AJ, Blair C, He X. 2015. RASP (Reconstruct Ancestral State in Phylogenies): a tool for historical biogeography. Molecular phylogenetics and evolution 87: 46–49.

Zachos JC, Dickens GR, Zeebe RE. 2008. An early Cenozoic perspective on greenhouse warming and carbon-cycle dynamics. Nature 451: 279–283.

Zachos J, Pagani M, Sloan L, Thomas E, Billups K. 2001. Trends, rhythms, and aberrations in global climate 65 Ma to present. Science 292: 686–693.

Zazanashvili. 2009. The Caucasus Hotspot. Status and protection of globally threatened species.

Zazanashvili, Sanadiradze, Bukhnikashvili. 2004. Caucasus. *., Mittermaier, CG, Lamoreux …*.

Zhang J-Q, Meng S-Y, Allen GA, Wen J, Rao G-Y. 2014. Rapid radiation and dispersal out of the Qinghai-Tibetan Plateau of an alpine plant lineage Rhodiola (Crassulaceae). Molecular phylogenetics and evolution 77: 147–158.

Zografidis A, Fragman-Sapir O, Strid A, Dimopoulos P. 2020. Notes on the generic name Graecobolanthus (Caryophylleae, Caryophyllaceae). Taxon 69: 992–997.

Zohary M. 1973. Geobotanical foundations of the Middle East. Fischer.

