## Supplementary Figures and Tables for "Evolutionary dynamics in the Irano-Anatolian and Caucasus biodiversity hotspots: Evolutionary radiation and its drivers in *Gypsophila* (Caryophyllaceae)"

[1]Department of Plant Science, Center of Excellence in Phylogeny of Living Organisms, School of Biology, College of Science, University of Tehran, PO Box 14155-6455 Tehran, Iran; [2]School of Life Sciences, University of Nevada, Las Vegas, Las Vegas, NV 89119, U.S.A.; [3]University of Michigan Herbarium-EEB, 3600 Varsity Drive, Ann Arbor, Michigan 48108-2228, U.S.A.; [4]Biodiversity Research – Systematic Botany, Department of Biology I, Ludwig–Maximilians Universität München, Menzinger Str. 67, D-80638 München, Germany and GeoBio Center LMU; [5]Department of Cell and Molecular Biology, School of Biology, College of Science, University of Tehran, PO Box 14155-6655 Tehran, Iran.

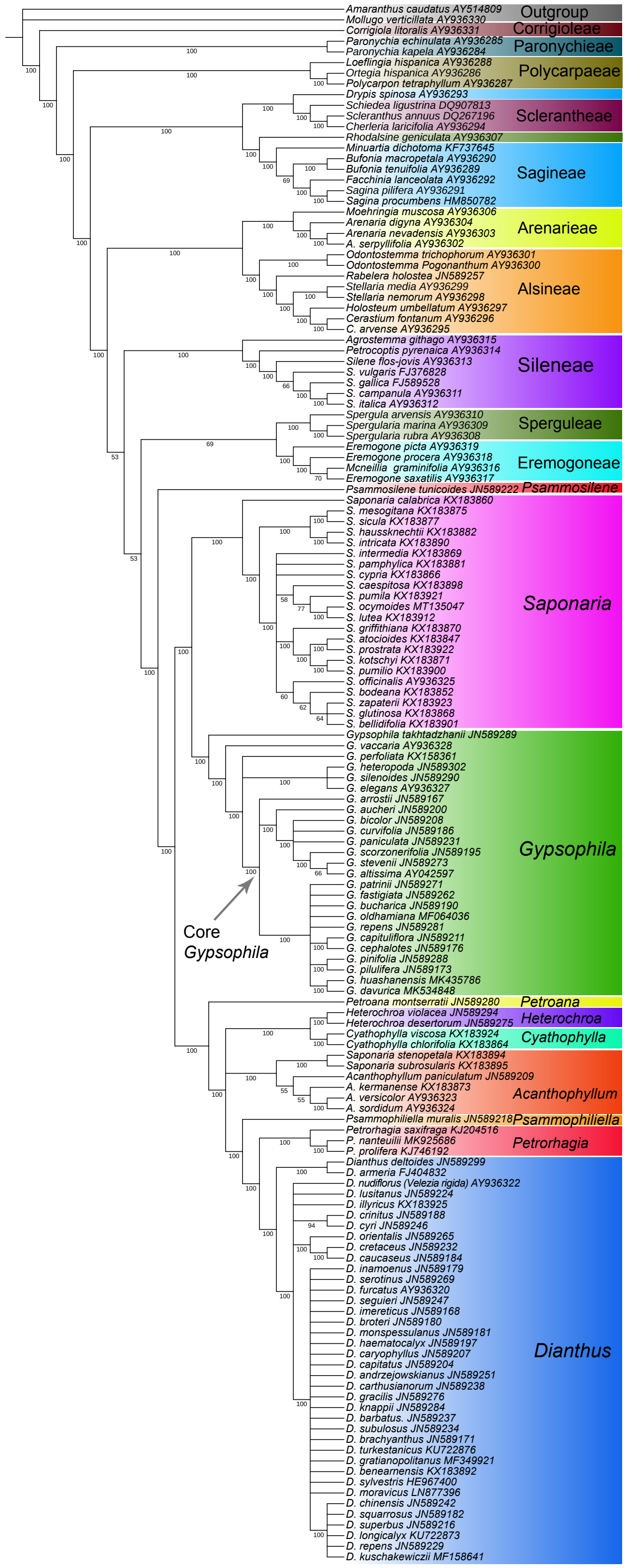

**Fig. S1** Maximum parsimonious tree of the Caryophyllaceae family reconstructed using the *mat*K dataset.

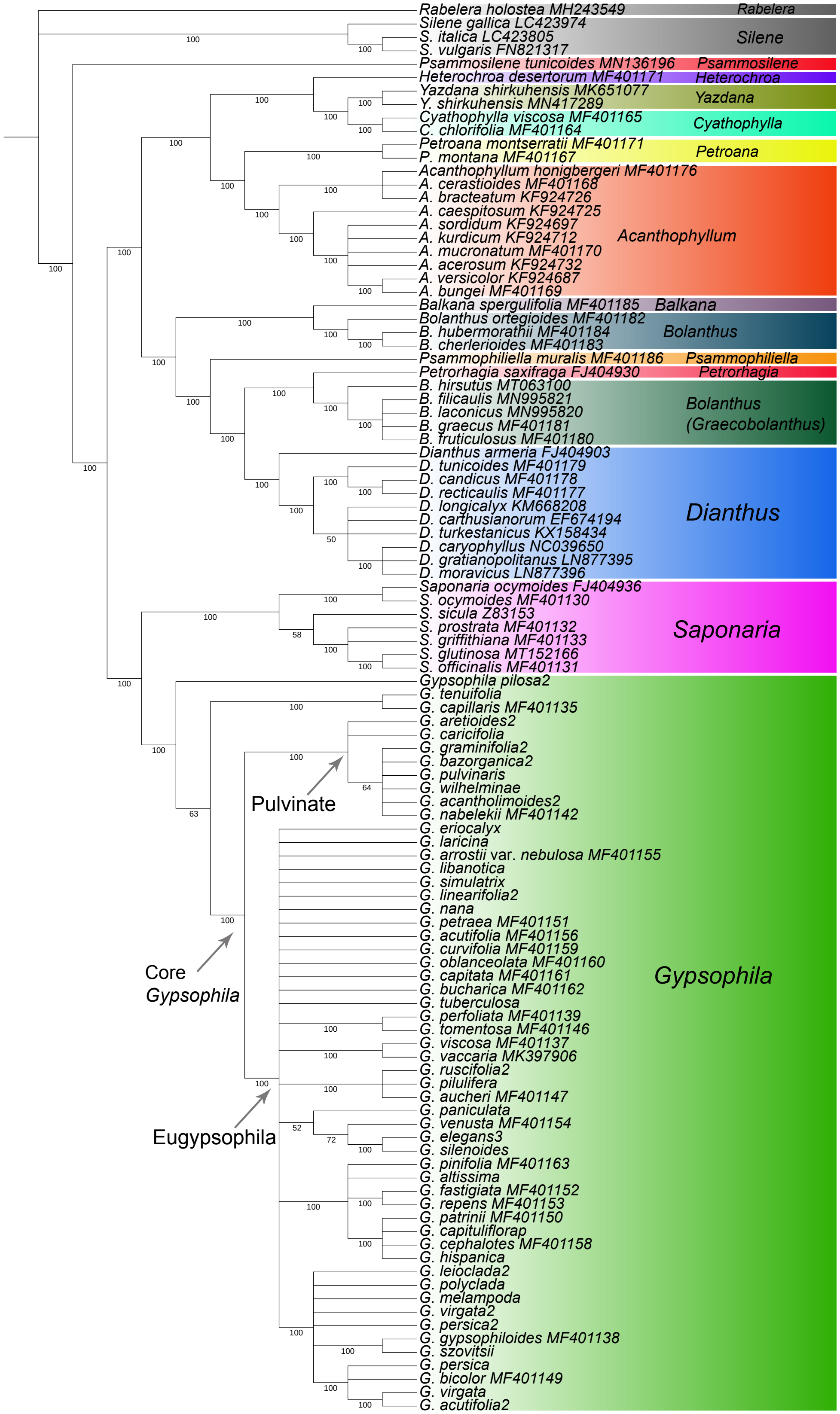

**Fig. S2** Maximum parsimonious tree of the Caryophylleae tribe reconstructed using the *rps16* dataset.

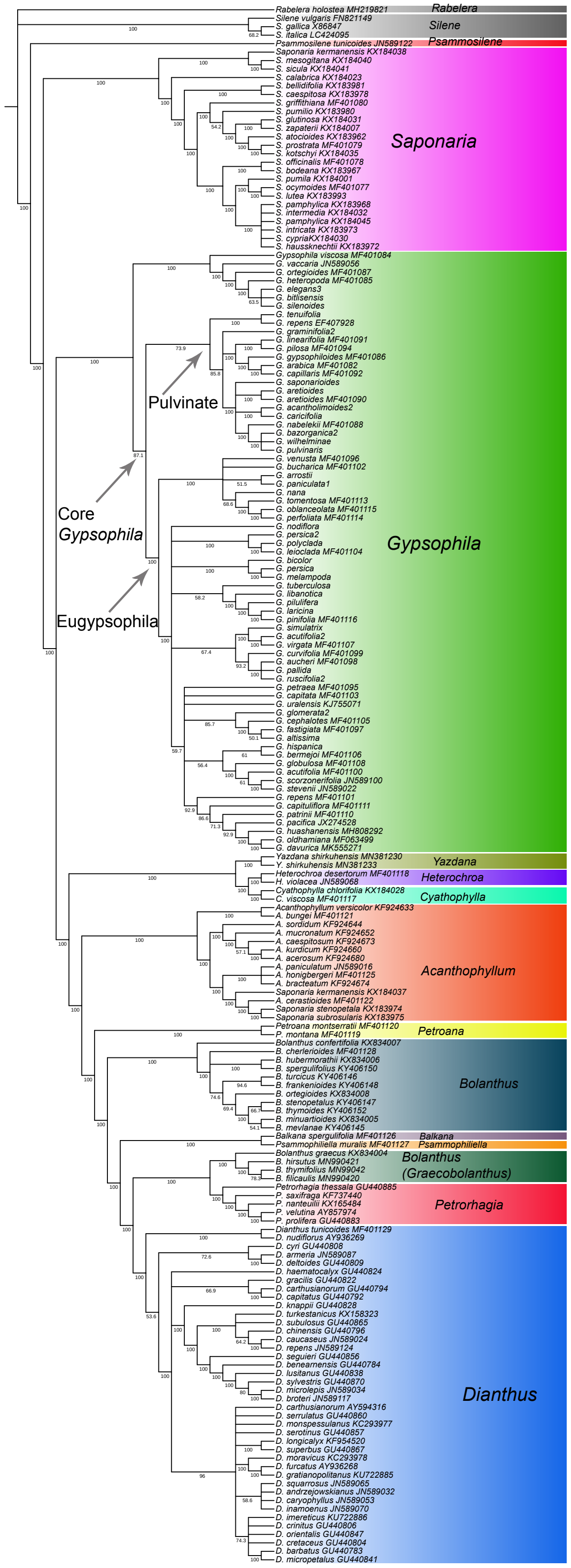

**Fig. S3** Maximum parsimonious tree of the Caryophylleae tribe reconstructed using the ITS dataset.

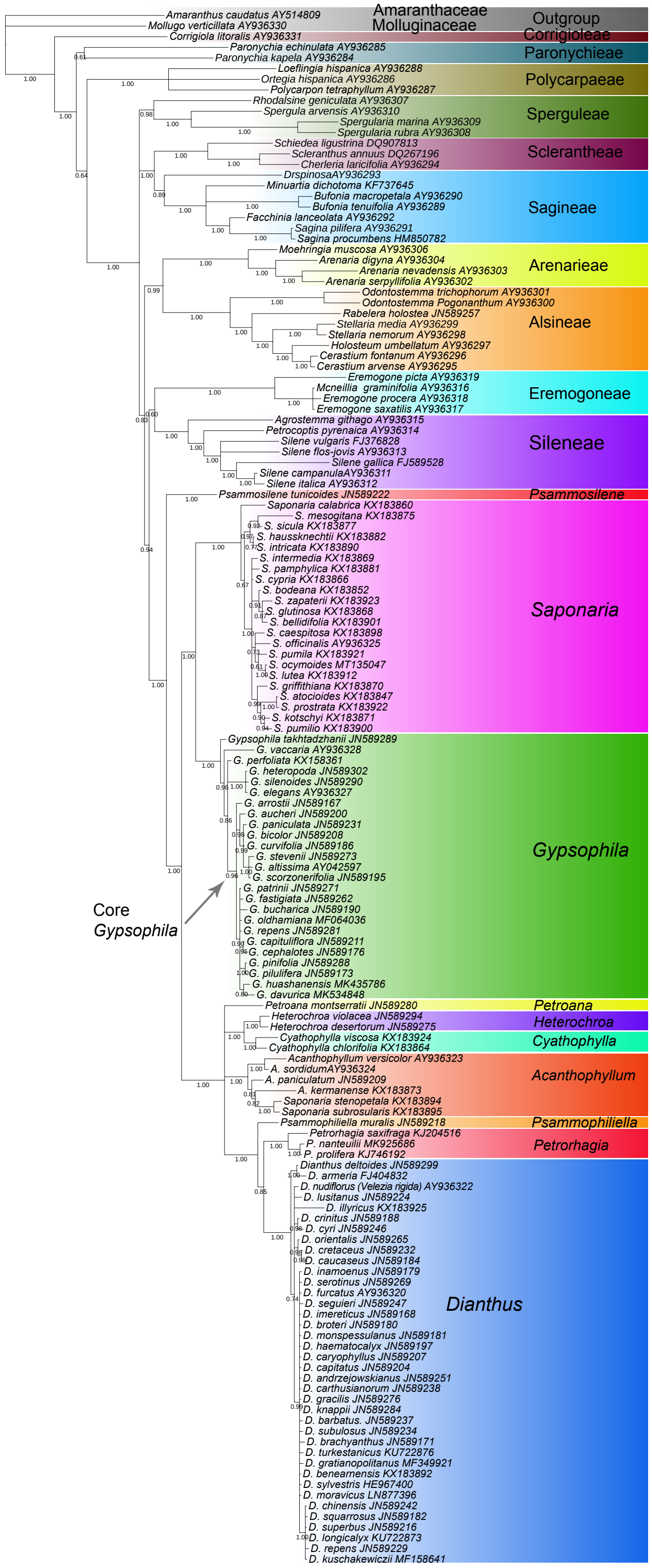

**Fig. S4** Majority-rule consensus tree of the Caryophyllaceae family inferred from Bayesian analysis of the *mat*K dataset.

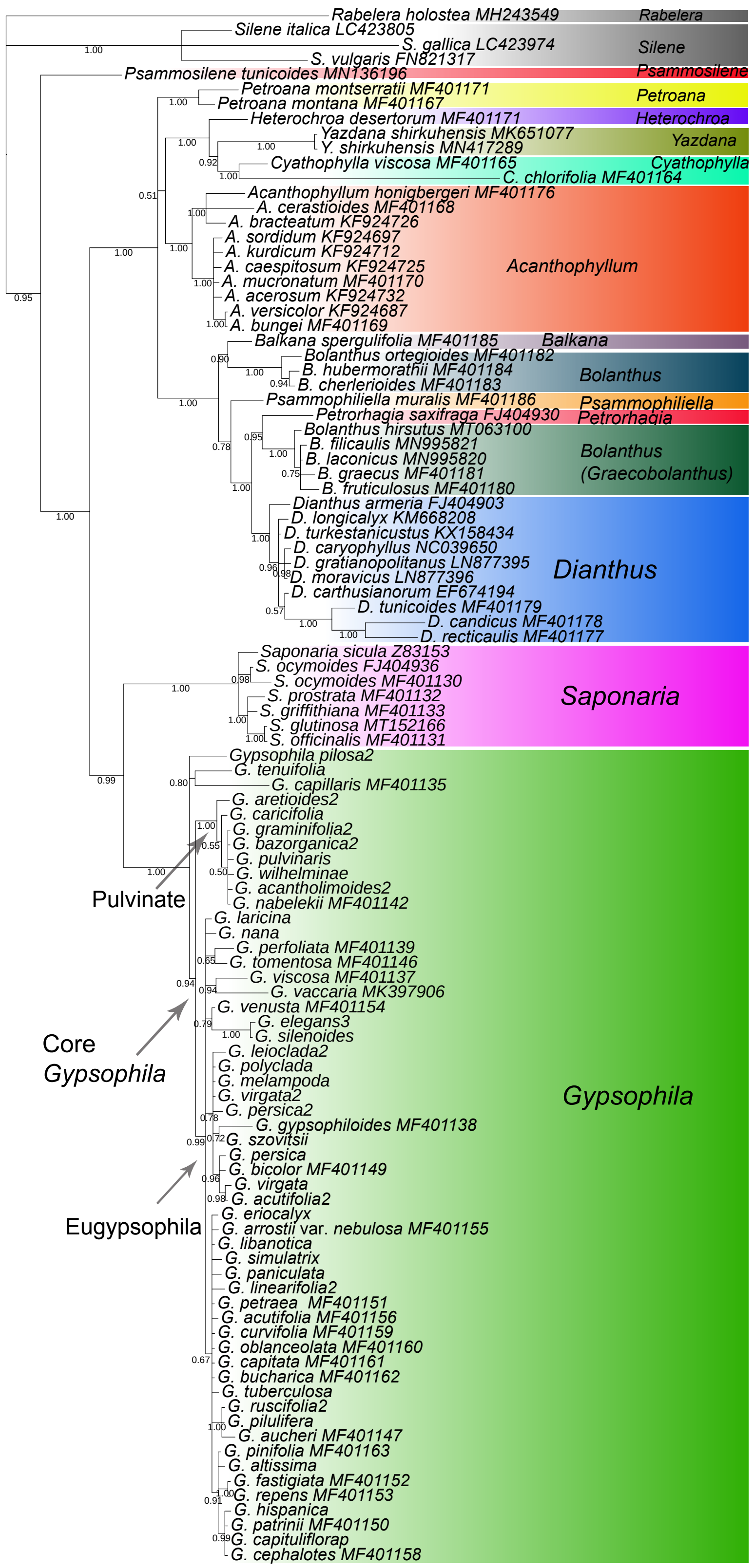

**Fig. S5** Majority-rule consensus tree of the Caryophylleae tribe inferred from Bayesian analysis of the *rps16* dataset.

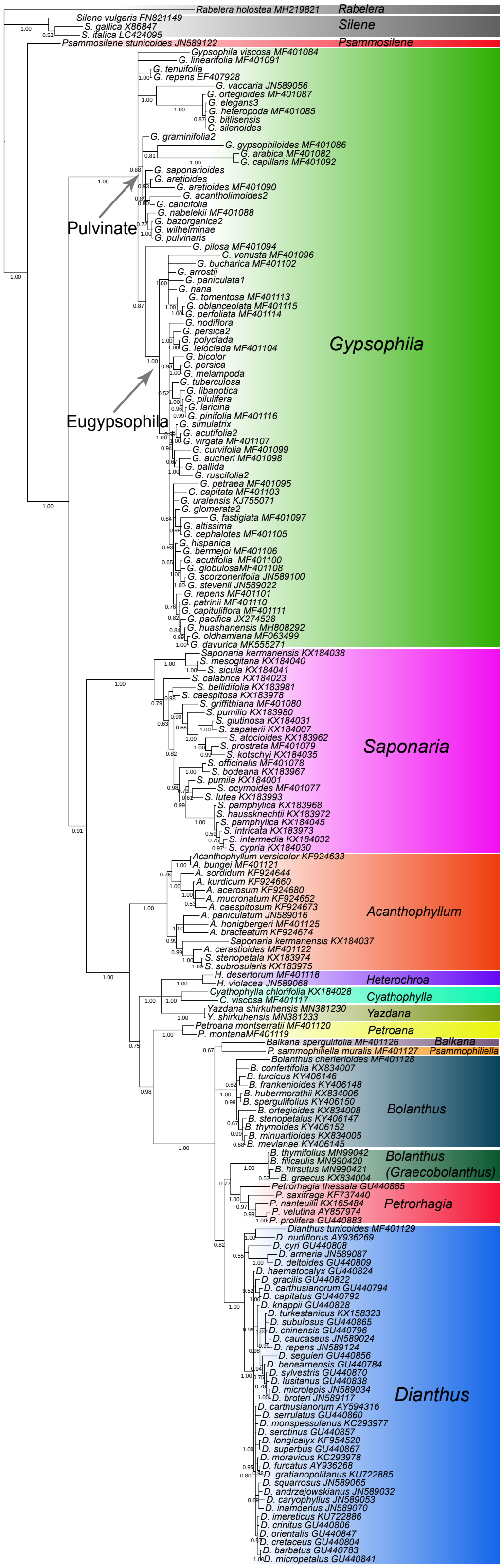

**Fig. S6** Majority-rule consensus tree of the Caryophylleae tribe inferred from Bayesian analysis of the ITS dataset.

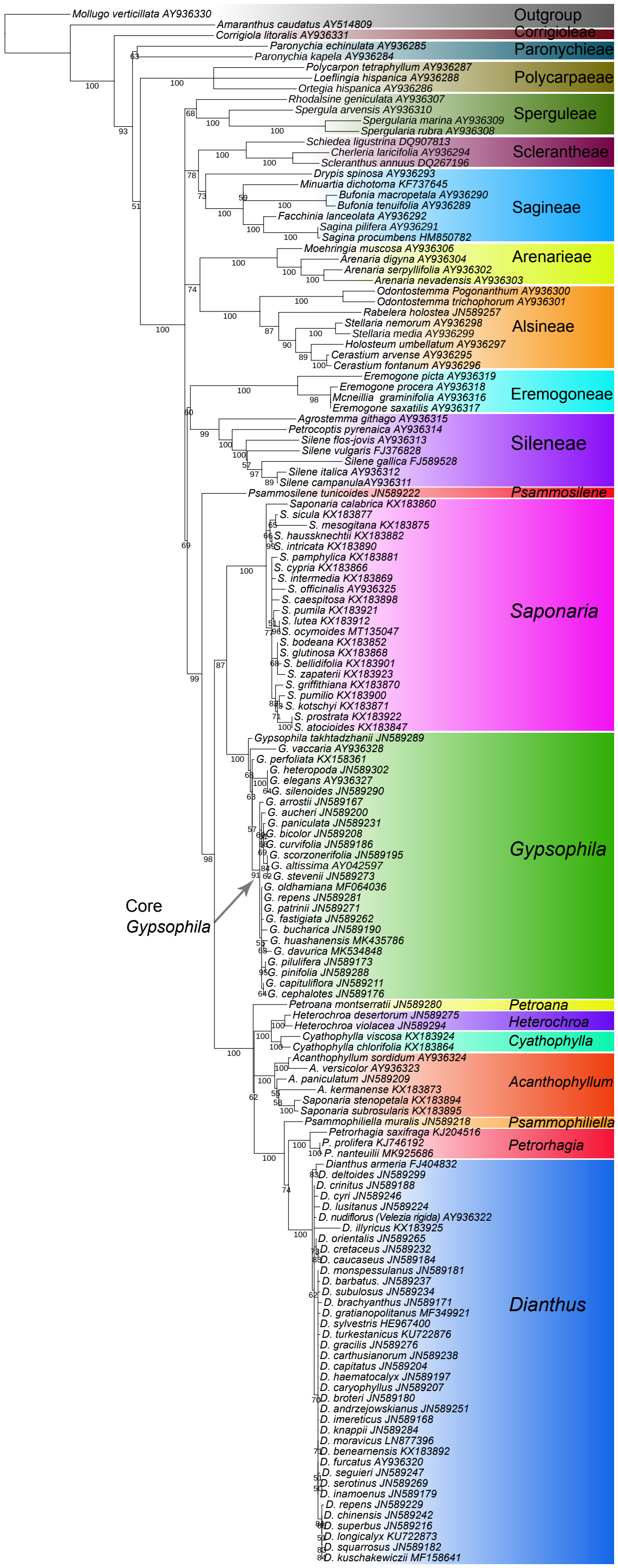

**Fig. S7** Maximum likelihood tree of the Caryophyllaceae family with bootstrap values inferred by RAxML using the *mat*K dataset.

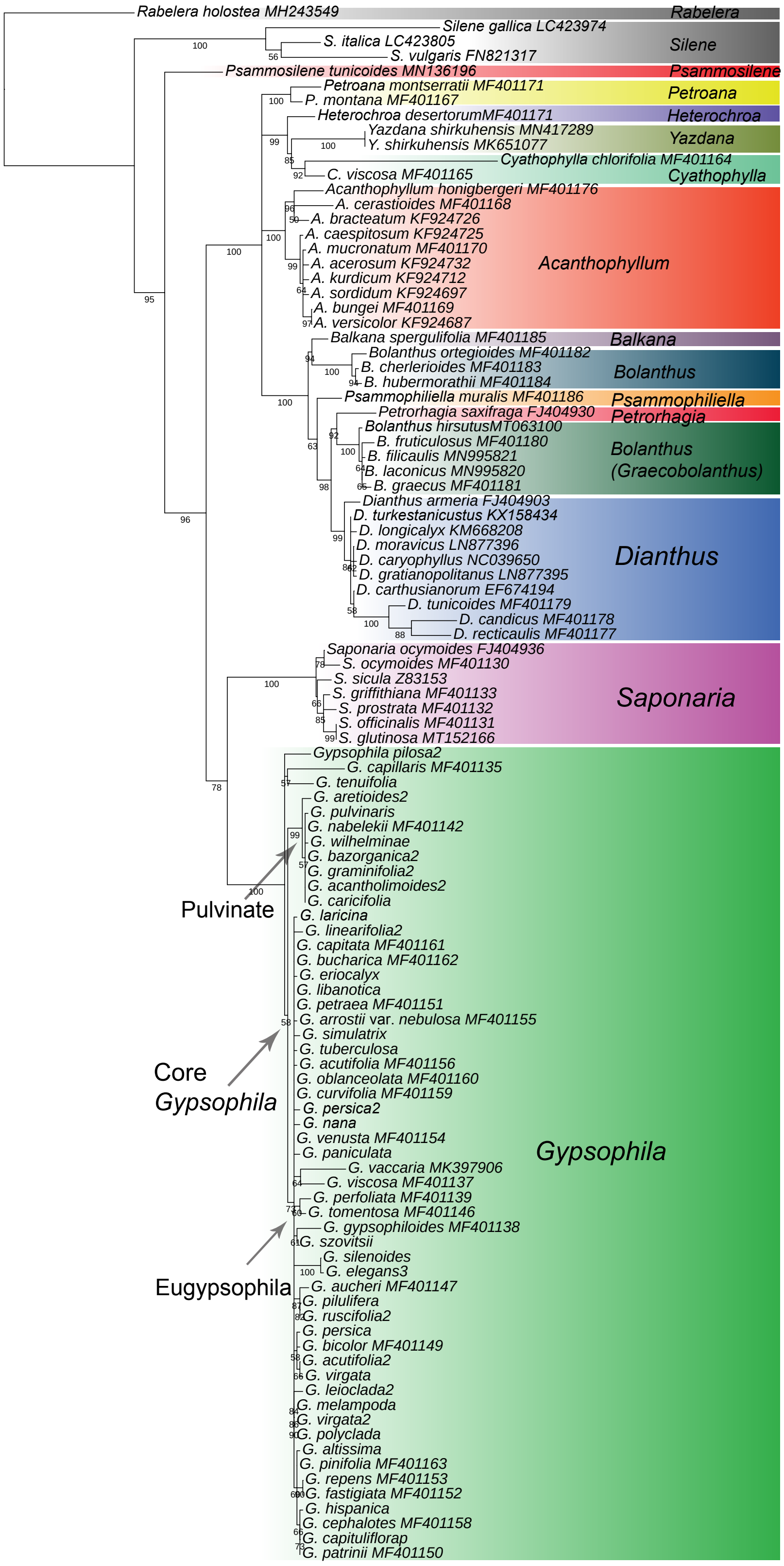

**Fig. S8** Maximum likelihood tree of the Caryophylleae tribe with bootstrap values inferred by RAxML using the *rps16* dataset.

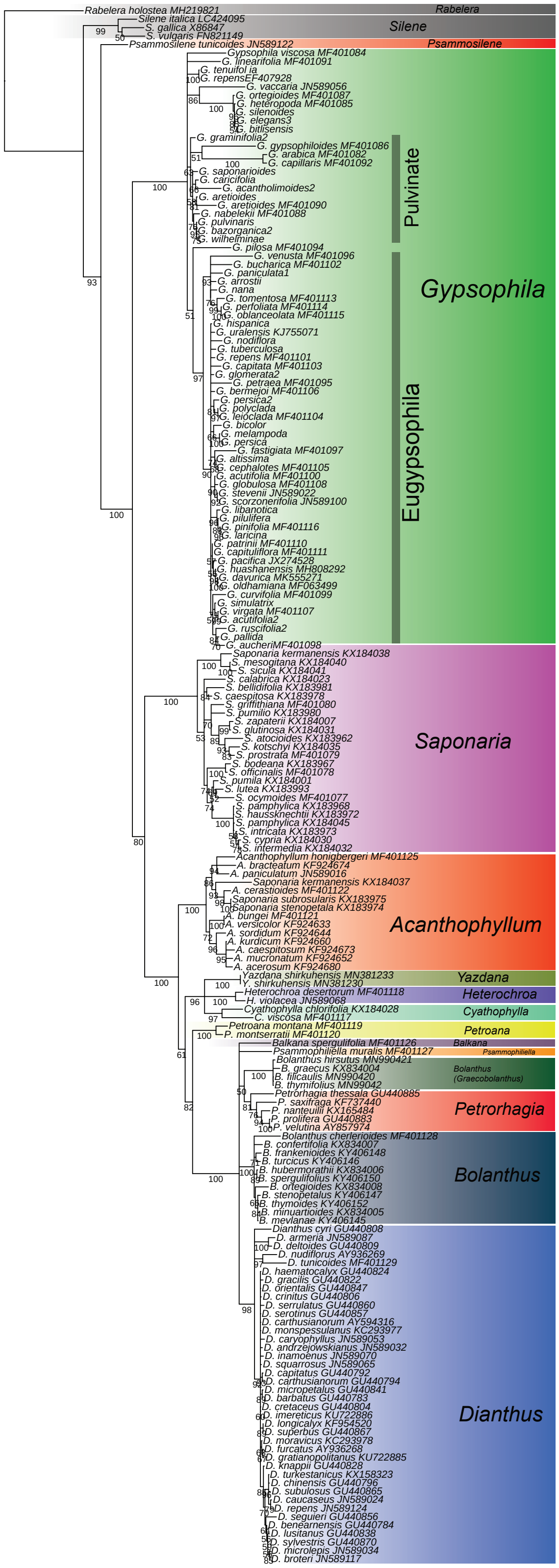

**Fig. S9** Maximum likelihood tree of the Caryophylleae tribe with bootstrap values inferred by RAxML using the ITS dataset.

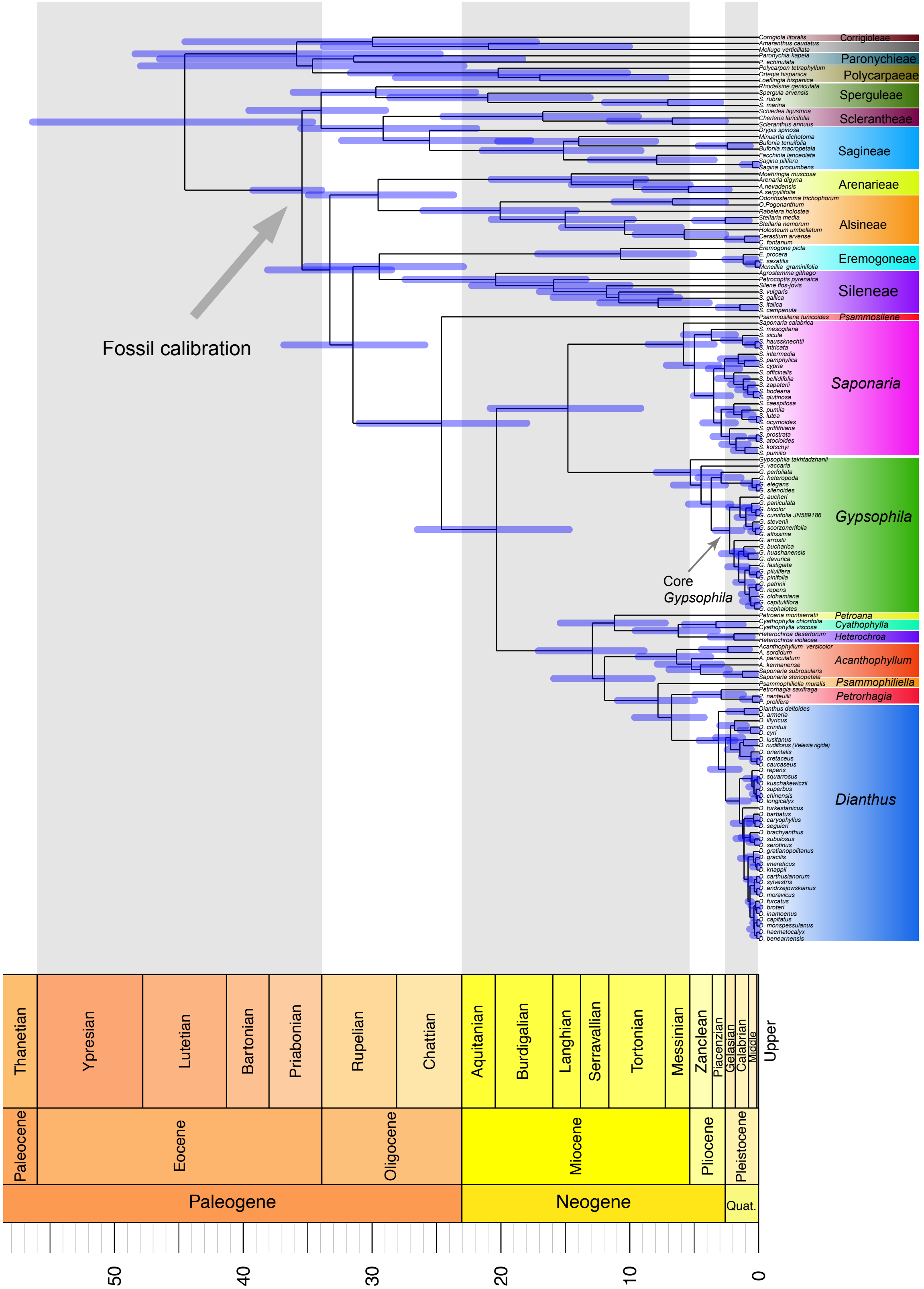

**Fig. S10** Maximum clade credibility tree from the BEAST analysis of the *mat*K dataset. The fossil calibration point is indicated by the grey arrow.

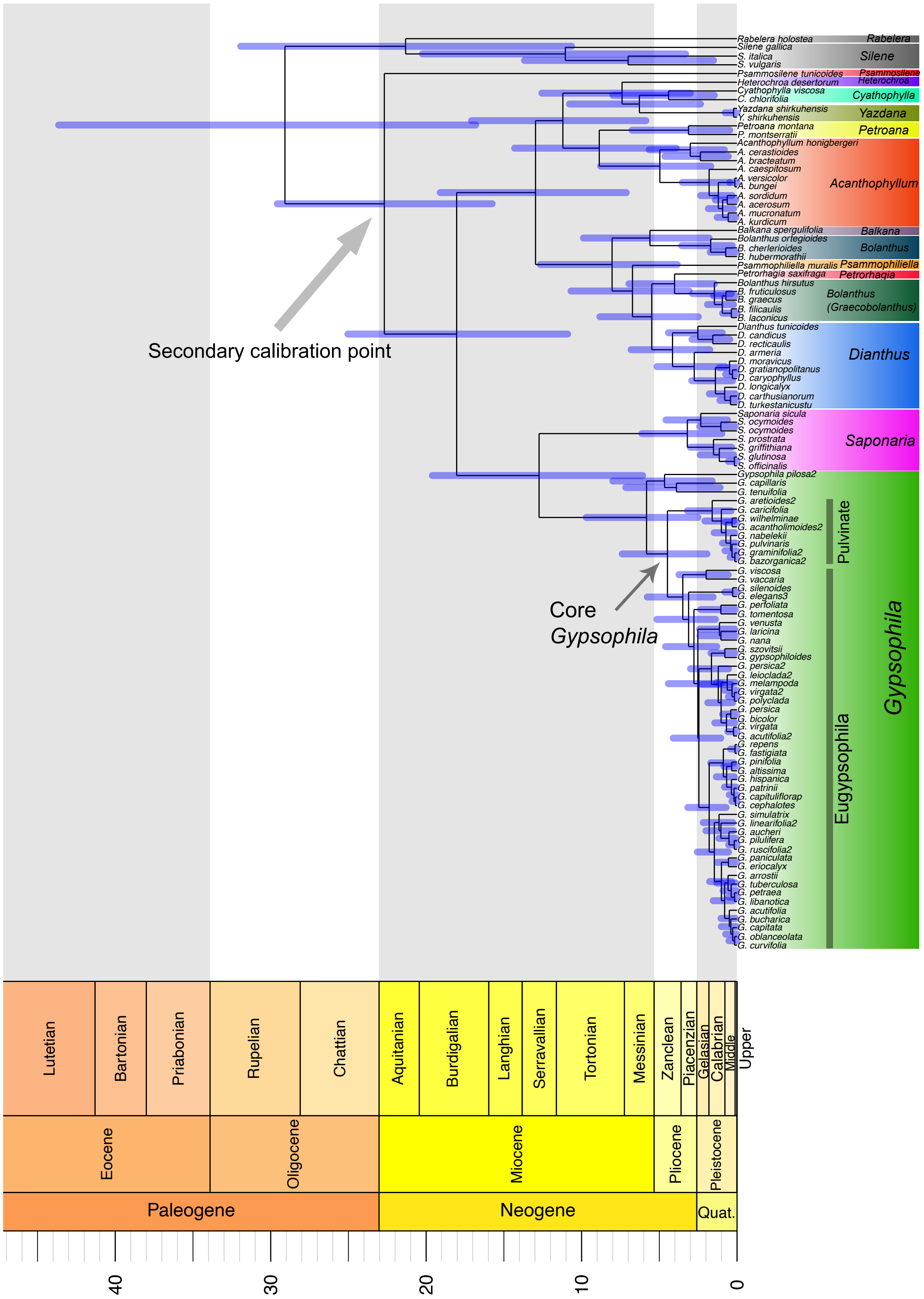

**Fig. S11** Maximum clade credibility tree from the BEAST analysis of the *rps16* dataset. The secondary calibration point is indicated by the grey arrow.

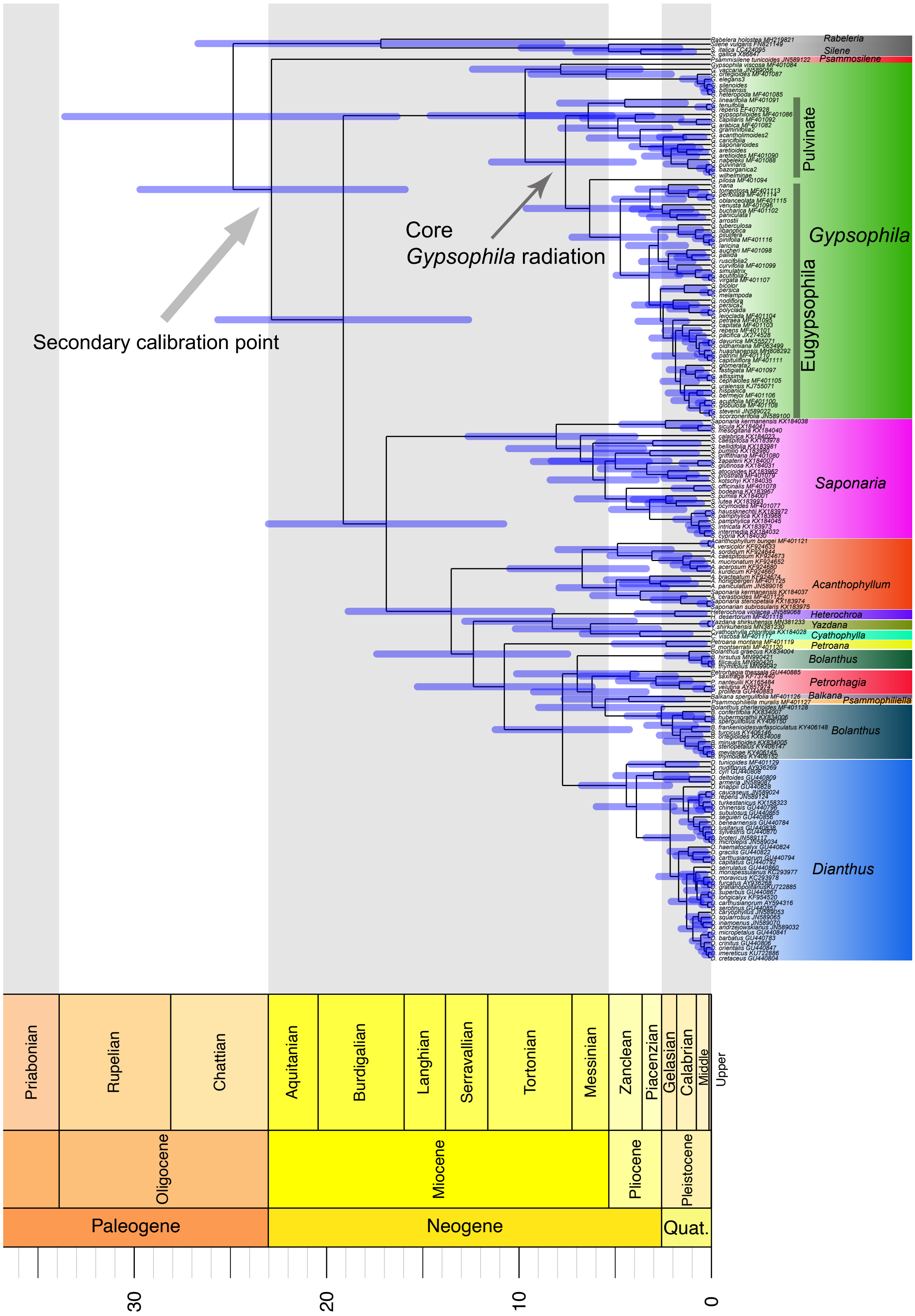

**Fig. S12** Maximum clade credibility tree from the BEAST analysis of the ITS dataset. The secondary calibration point is indicated by the grey arrow.

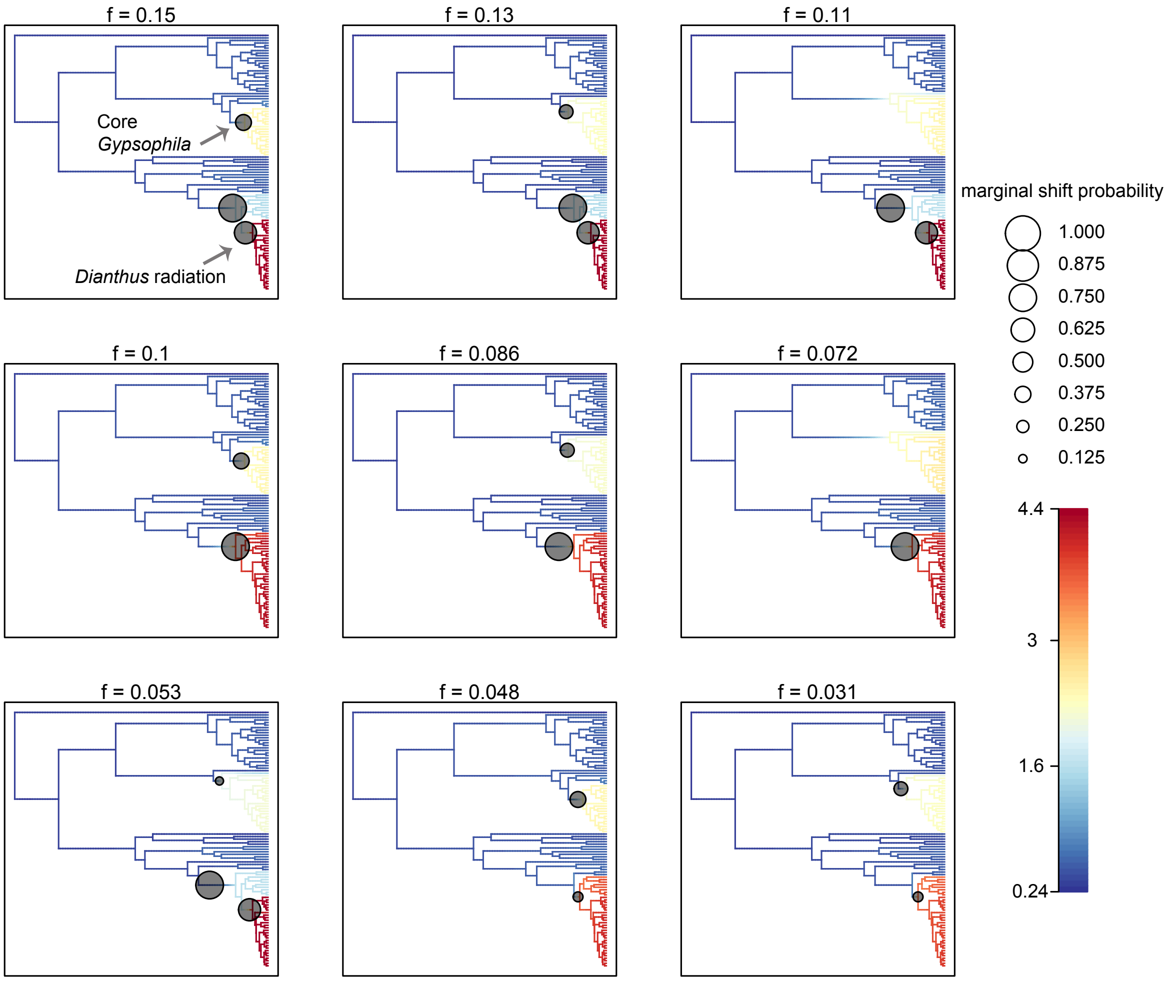

**Fig. S13** Credible set of shift configurations from BAMM analysis of the *mat*K dataset. Significant rate changes are designated by a circle along the corresponding branches at which the shift in speciation rate occurred.

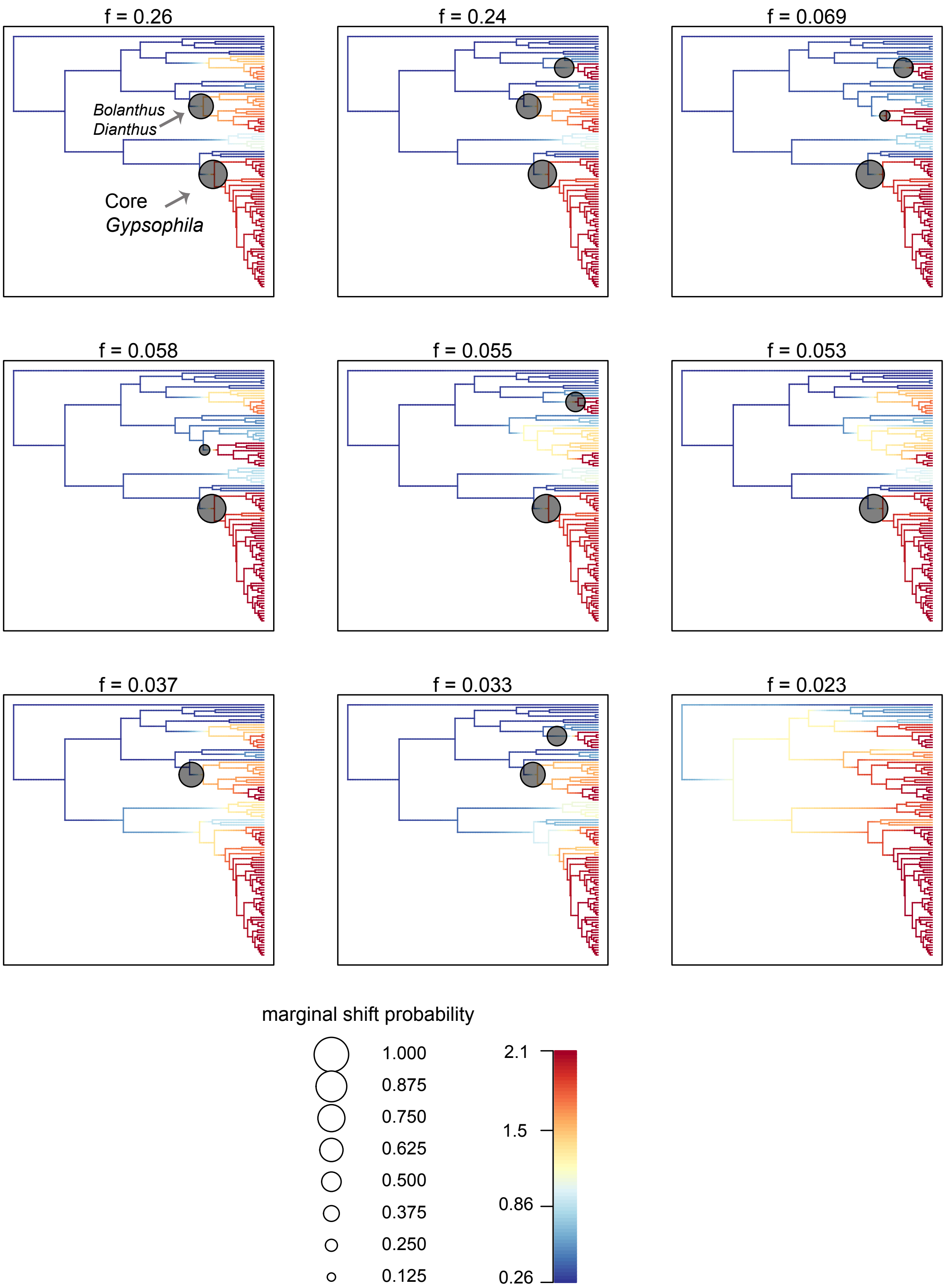

**Fig. S14** Credible set of shift configurations from BAMM analysis of the *rps16* dataset. Significant rate changes are designated by a circle along the corresponding branches at which the shift in speciation rate occurred.

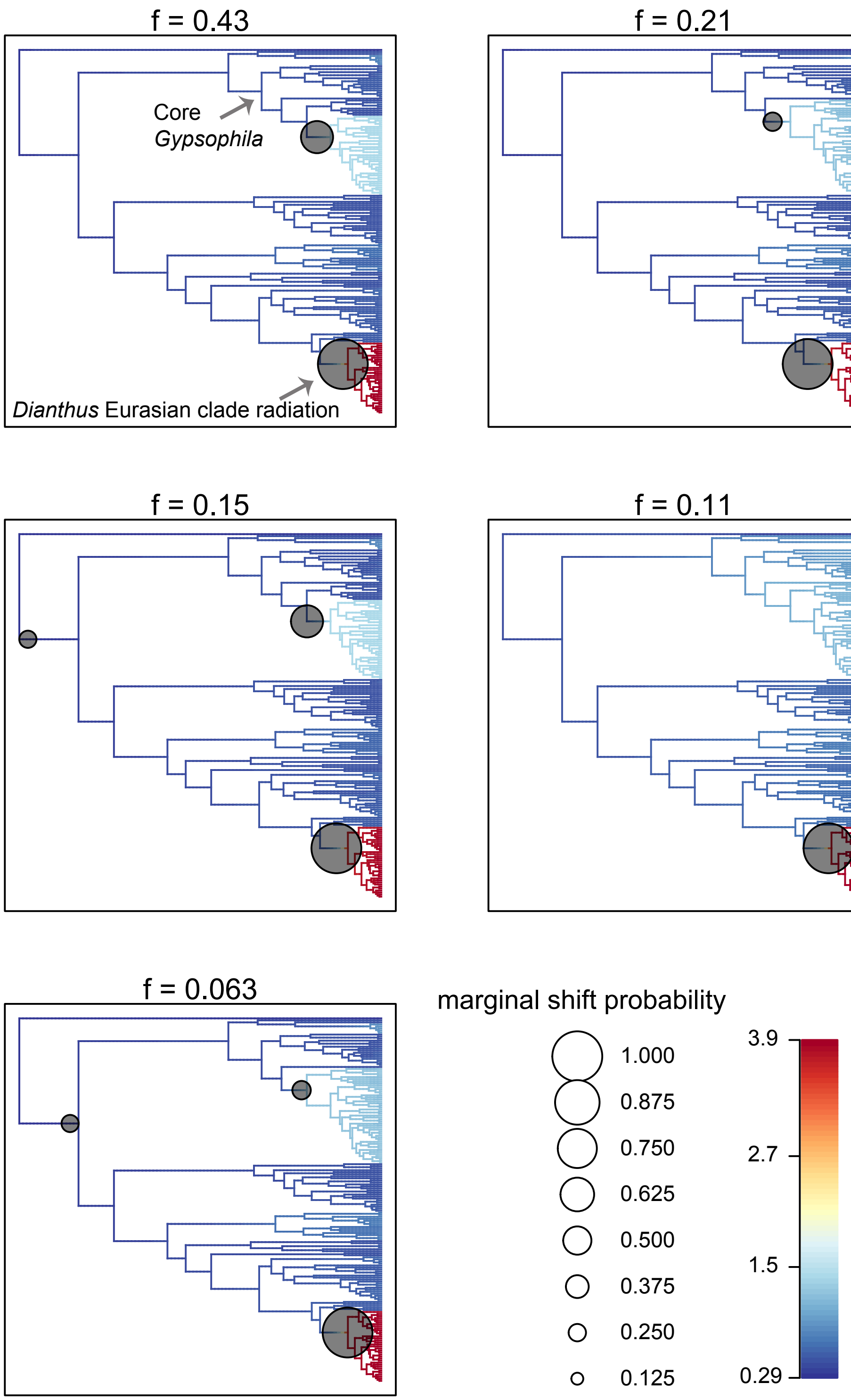

**Fig. S15** Credible set of shift configurations from BAMM analysis of the ITS dataset. Significant rate changes are designated by a circle along the corresponding branches at which the shift in speciation rate occurred.

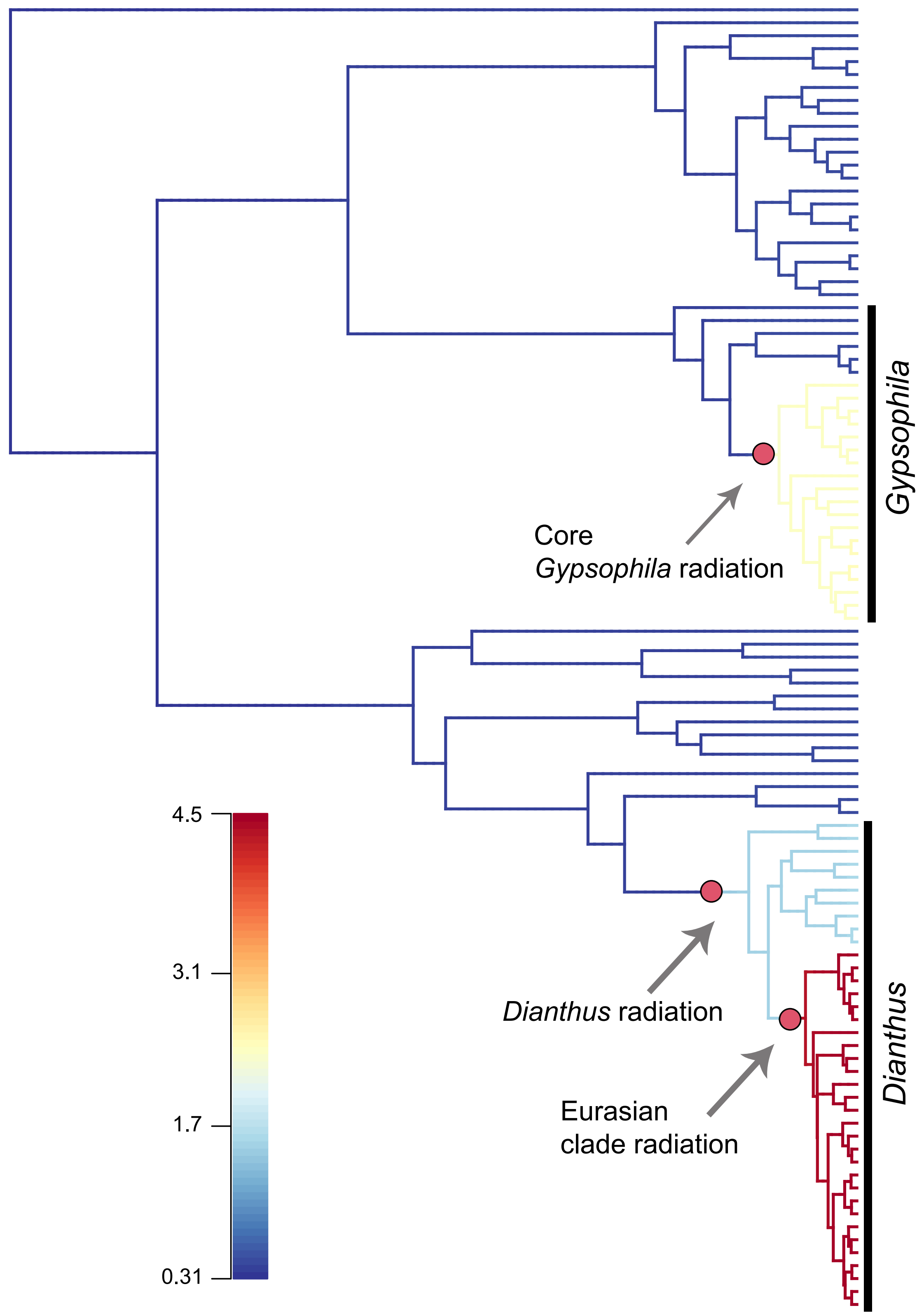

**Fig. S16** Best shift configurations from BAMM analysis of the *mat*K dataset. The best shift configurations in the speciation rates are indicated by a red circle along the corresponding branches.

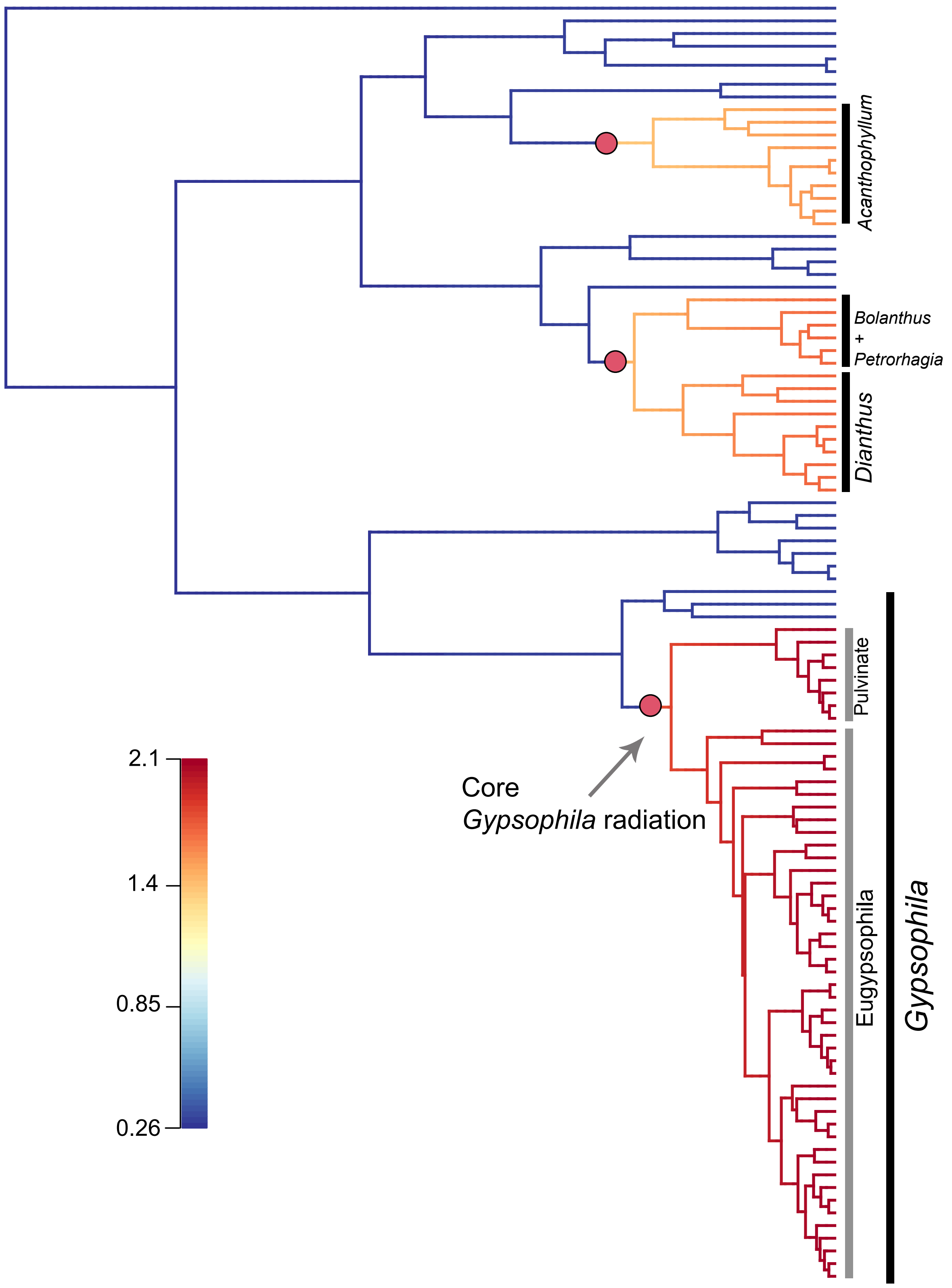

**Fig. S17** Best shift configurations from BAMM analysis of the *rps16* dataset. The best shift configurations in the speciation rates are indicated by a red circle along the corresponding branches.

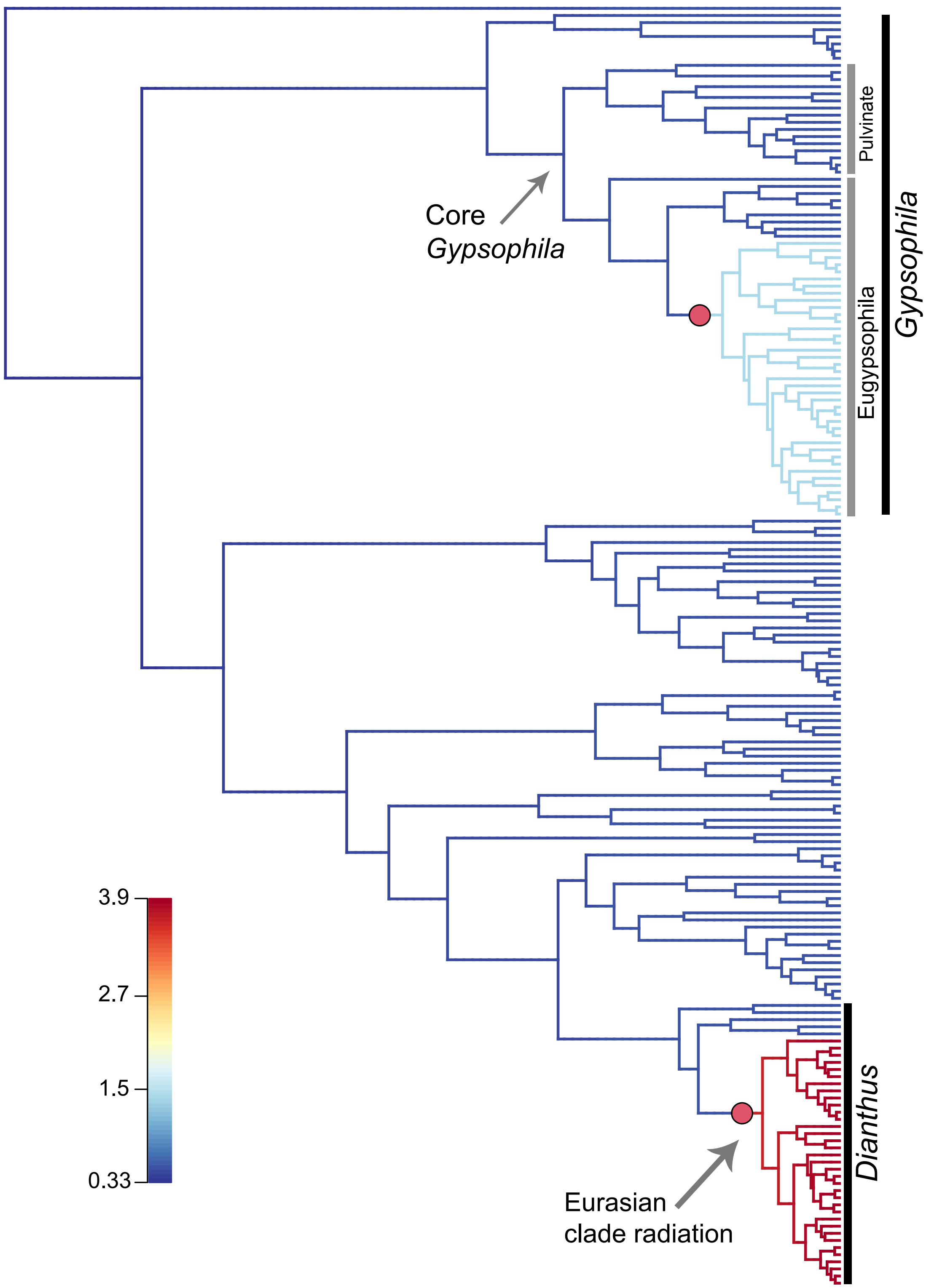

**Fig. S18** Best shift configurations from BAMM analysis of the ITS dataset. The best shift configurations in the speciation rates are indicated by a red circle along the corresponding branches.

**Table S1** Summary of studies on dynamics of diversification (reports of evolutionary radiations) in 36 global biodiversity hotspots.

| Continent | Biodiversity Hotspots | Taxa | Species number | Clade age in MA (95% CI) | Net Diversification Rate events Myr–1 per lineage | Constraints and Methods of Calibration | Diversification rate analyses | References |
|---|---|---|---|---|---|---|---|---|
| **South America** | **(1) Tropical Andes** | Neotropical bellflowers (Campanulaceae: Lobelioideae): Centropogonid clade | ~550 | 5.02 (3.95–6.13) | 1.06 (0.87–1.35) when ε= 0.5 | Fossil calibration using both treePL and BEAST | Magallon & Sanderson approach; BAMM; RPANDA | (Lagomarsino *et al.*, 2016) |
| | | *Lupinus* (Andean clade) | ~85 | 1.47 (1.18–1.76) | 2.49–3.72 | Fossil calibration using R8S | Diversification rates calculated using simple Yule model: R = (ln n1 − ln n0)/t Where, n1 is extant species, n0 is the initial species number (usually 1), and t is time in Myr | (Hughes & Eastwood, 2006) |
| | | High elevation paramo Valerianaceae: *Valeriana* | 53 | 14.58 (3.47–4.98) | 0.80–1.34 | Fossil data, Andes uplift time, respectively, the previous estimate of South American colonization by Valerianaceae Bayesian relaxed clock implemented in MULTIDIVTI | R = (ln n1 − ln n0)/t | (Bell & Donoghue, 2005) |

| | | | | | | | | |
|---|---|---|---|---|---|---|---|---|
| | | | | | | ME software. NPRS implemented in r8s | | |
| | | South American *Hypericum* (Hypericaceae): Páramo clade (sect. *Brathys*) | 63 | 3.83 (2.26–5.62) | 1.098 (0.748–1.860) | Fossil and secondary calibration | Magallon & Sanderson approach | (Nürk *et al.*, 2013) |
| | **(2) Tumbes–Choco–Magdalena** | Bromeliaceae; *Ronnbergia* | 26 | 3.5 (2.9–4.1) | 0.68 (0.38–1.03) | Secondary calibration using BEAST | Geiger; BAMM | (Aguirre-Santoro *et al.*, 2019) |
| | **(3) Atlantic Forest** | Bromeliaceae; Brazilian *Wittmackia* | ~ 24 | 3.2 (2.6–3.8) | 1.01 (0.68–1.41) | Secondary calibration using BEAST | Geiger; BAMM | (Aguirre-Santoro *et al.*, 2019) |
| | | Asteraceae; Brazilian Eupatorieae | ~ 247 | <7 | – | Based on the BEAST calibration on the common ancestor with their sister clade in another study (Tippery et al. 2014) | – | (Rivera *et al.*, 2016) |
| | **(4) Cerrado** | Asteraceae; Brazilian Eupatorieae | ~ 247 | <7 | – | Based on the BEAST calibration on the common ancestor with their sister clade in | – | (Rivera *et al.*, 2016) |

| | | | | | | | | |
|---|---|---|---|---|---|---|---|---|
| | | | | | | another study (Tippery et al. 2014) | | |
| | **(5) Chilean Winter Rainfall and Valdivian Forests** | – | – | – | – | – | – | – |
| **North and Central America** | **(6) Caribbean Islands** | Bromeliaceae; Caribbean *Wittmackia* | ~ 17 | 1 (0.8–1.3) | 1.16 (0.61–2.07) | BEAST | Geiger; BAMM | (Aguirre-Santoro *et al.*, 2019) |
| | **(7) Mesoamerica** | Zamiaceae; *Zamia* | ~ 75 (~ 20 in Mesoamerica) | 9.54 (9.0–10.62) for *Zamia* crown group 3.9 (2.54–5.34) for Mesoamerican clade | ~ 0.39 (ε = 0) in *Zamia* crown group and 0.60 (ε = 0) in Mesoamerican clade | Fossil calibration using BEAST | Geiger; BAMM | (Calonje *et al.*, 2019) |
| | **(8) Madrean Pine–Oak Woodlands** | – | – | – | – | – | – | – |
| | **(9) North American Coastal Plain** | – | – | – | – | – | – | – |
| | **(10) California Floristic Province** | Rhamnaceae; *Ceanothus* | ~ 53 | 12.9 (4.2–22.1) | – | Fossil calibration using BEAST | – | (Calsbeek *et al.*, 2003; Burge *et al.*, 2011) |
| **Africa** | **(11) Cape Floristic Region** | Rhamnaceae; Phylica | ~ 150 | ~ 7.22 | – | Island age using NPRS | – | (Richardson *et al.*, 2001) |
| | | Ericaceae; Erica | ~ 800 (~ 690 endemic) | 19.31 (12–27) for *Erica* 9.15 (11–6) for Cape clade | 0.39–0.97 (ε = 0) in Cape *Erica* | Fossil and secondary calibration using BEAST | Geiger | (Richardson *et al.*, 2001; Linder, 2003; Linder & Hardy, 2004) |

| | | | | | | | | |
|---|---|---|---|---|---|---|---|---|
| | **(12) Succulent Karoo** | Aizoaceae; Core Ruschioideae | 1,563 | 3.8–8.7 | 0.77–1.75 | NPRS implemented in TreeEdit | R = (ln n1 − ln n0)/t | (Klak *et al.*, 2004) |
| | **(13) Maputaland–Pondoland–Albany** | The dwarf chameleons (Chamaeleonidae ; Bradypodion) | 15 | 14.8 (8.5–23.2) | – | Fossil calibration using Bayesian relaxed molecular clock | LTT plots; Symmetree (Chan & Moore, 2005) | (Tolley *et al.*, 2008) |
| | **(14) Madagascar and Indian Ocean Islands** | Paussus (ant–nest beetles): Malagacy radiation | >86 | 2.6 (1.7–3.6) | – | Fossil calibration using BEAST | LTT plots; MEDUSA | (Moore & Robertson, 2014) |
| | | *Coffea* subgenus *Coffea* L. (Rubiaceae) | ~ 95 (~ 58 in Madagascar) | 0.1–0.46 | – | Fossil and Island age | – | (Anthony *et al.*, 2010) |
| | **(15) Eastern African montane** | Zosteropidae; *Zosterops* | ~ 100 species (~ 15 taxa in the hotspot) | 1.55 (0.97–2.5) | – | Island age using BEAST | – | (Cox *et al.*, 2014) |
| | | Hypericaceae; *Hypericum* | ~ 20 | 1.7–7.9 | – | Fossil calibration using NPRS implemented in TreeEdit and BEAST | – | (Meseguer *et al.*, 2013) |
| | **(16) Guinean Forests of West Africa** | *Coffea* subgenus *Coffea* L. (Rubiaceae) | ~ 95 (~ 17 in Guinea) | 0.1–0.46 | – | Fossil and Island age calibration | – | (Anthony *et al.*, 2010) |
| | **(17) Horn of Africa** | Gekkonidae; *Hemidactylus* African radiation | ~ 90 (~ 22 in African radiation) | 9.8 (6.5–13.6) | – | Secondary calibration using BEAST, and fossil cross-validation method | – | (Smíd *et al.*, 2013) |

| | | | | | | | | |
|---|---|---|---|---|---|---|---|---|
| | **(18) Coastal Forests of Eastern Africa** | Coffea subgenus *Coffea* L. (Rubiaceae) | ~ 95 (~ 11 in the hotspot) | 0.1–0.46 | – | Fossil and Island age calibration | – | (Anthony *et al.*, 2010) |
| **Europe** | **(19) Mediterranean Basin** | Caryophyllaceae; *Dianthus* | ~ 300 (> 200 in Eurasian radiation) | 1.9–7.0 | 2.21–7.55 | Fossil calibration using BEAST and PATHd8 | Geiger | (Valente *et al.*, 2010) |
| **West Asia** | **(20) Irano–Anatolian** | **Caryophyllaceae; *Gypsophila*** | **~ 149 (>100 in the hotspot)** | **5.31 (3.02–7.95)** | **0.81 (ε = 0)<br>0.51 (ε = 0.9)** | **Fossil calibration using BEAST** | **Geiger; RPANDA; BAMM** | **This study** |
| | **(21) Caucasus** | **Caryophyllaceae; *Gypsophila*** | **~149 (>20 in the hotspot)** | **5.31 (3.02–7.95)** | **0.81 (ε = 0)<br>0.51 (ε = 0.9)** | **Fossil calibration using BEAST** | **Geiger; RPANDA; BAMM** | **This study** |
| **Central and East Asia** | **(22) Mountains of Central Asia** | Saxifragaceae; *Saxifraga* section *Ciliatae* | ~175 | 26.21 (19.95–34.00) | min= 0.455<br>max= 0.486 | Fossil calibration using BEAST | BAMM | (Ebersbach *et al.*, 2017a,b) |
| | **(23) Western Ghats and Sri Lanka** | Bufonidae; Adenominae radiation | ~ 40 | 22.42 (17.14–31.05) | – | Fossil calibration using BEAST | – | (Van Bocxlaer *et al.*, 2009) |
| | **(24) Mountains of Southwest China** | Poaceae; Bambusoideae; Arundinarieae; *Fargesia* and *Yushania* (Alpine bamboos) | ~ 90 and ~80 | 7.36 (4.7–11.91) | 0.75 (0.60–0.89) | Secondary calibration using BEAST | BAMM | (Ye *et al.*, 2019) |
| | **(25) Himalaya** | Ranunculaceae; *Delphinium* subg. *Delphiniastrum* | ~ 300 | 6.19–13.56 | 0.37–0.81 | Fossil calibration using BEAST | Geiger | (Jabbour & Renner, 2012) |
| | **(26) Japan** | Gobiidae; *Luciogobius* | 11 | 7.67 (5.52–12.27) | – | Secondary calibration using r8s | – | (Yamada *et al.*, 2009) |
| | **(27) Indo–Burma** | Musaceae; *Musa* | ~83 | 37.9 (24.5–50.5) | 0.15 species **Ma–1** (95% HPD: 0.19–0.52) | Fossil calibration using BEAST | BAMM | (Janssens *et al.*, 2016) |

| | | | | | | | | |
|---|---|---|---|---|---|---|---|---|
| **Pacific Asia, Australia, and Pacific Islands** | **(28) Sundaland** | Gekkonidae; *Cyrtodactylus* | 150 | 52 (65–40) | – | Fossil calibration using BEAST | – | (Wood *et al.*, 2012) |
| | **(29) Wallacea** | Corvides (Crows and Jays, Birds–of–paradise, Vangas, and allies) | ~837 | 25.5 (25.06–25.99) | – | Fossil calibration using BEAST | – | (McCullough *et al.*, 2022) |
| | **(30) Philippines** | Soricidae; *Crocidura* Southeast Asian shrews | >200 (~ 9 in Philippines) | ~ 4.4 (colonization of Philippines and Sundaland) | – | Two plausible substitution rates using PAUP | Maximum likelihood approachimplemented in R package LASER; LTT in GENIE | (Esselstyn *et al.*, 2009) |
| | **(31) Southwest Australia** | Proteaceae; *Adenanthos* | 31 (29 in the hotspot) | 24 (15.3–33.2) | – | Fossil calibration using BEAST | LTT plots | (Nge *et al.*, 2021) |
| | **(32) Forests of East Australia** | Pygopodoid geckos (families: Carphodactylidae, Diplodactylidae, and Pygopodidae) | ~ 180 Carphodactylidae ~30 Diplodactylidae >100 Pygopodida 46 | 57 (50–64) Carphodactylidae: ~30 Diplodactylidae: ~30 Pygopodidae: ~25 | 0.0938 (BAMM estimate) | Fossil and secondary calibration using BEAST | BAMM; TreePar, LASER | (Brennan & Oliver, 2017) |
| | **(33) East Melanesian Islands** | *Exocelina* diving beetles (Coleoptera, Dytiscidae, Copelatinae) | ~200 (>150 in New Guinea) | *Exocelina:* 15.35 (11.20–19.92) New Guinean *Exocelina*: 5.37 (3.74–7.07) | Speciation rate in the New Guinean radiation = 0.785 the global *Exocelina* radiation rate = 0.497 | Fossil calibration using BEAST | BAMM | (Toussaint *et al.*, 2015) |

| | | | | | | | | |
|---|---|---|---|---|---|---|---|---|
| | **(34) New Caledonia** | *Exocelina* diving beetles (Coleoptera, Dytiscidae, Copelatinae) | ~200 (~40 in New Caledonia) | *Exocelina:* 15.35 (11.20–19.92) New Caledonia *Exocelina*: 7.21 (4.31–10.19) | Speciation rate in the New Guinean radiation = 0.320 the global *Exocelina* radiation rate = 0.497 | Fossil calibration using BEAST | BAMM | (Toussaint *et al.*, 2015) |
| | **(35) New Zealand** | New Zealand rockcresses (Brassicaceae; *Pachycladon*) | 11 | 0.82 | Speciation rate = 1.99 (0.35–2.19) Extinction rate = 0.005 (0–0.92). | Synonymous substitution estimate obtained from 256 genes and secondary calibration using BEAST | LASER | (Joly *et al.*, 2009, 2014) |
| | **(36) Polynesia–Micronesia (including Hawaiian Islands)** | Silversword alliance | 30 | 5.2 (4.4–6.0) | The Kendall/Moran estimate of diversification rate: Ŝ= 0.56 ± 0.17 species per million years | Fossil evidence paleoclimatic data usng r8s | R = (ln n1 − ln n0)/t and The Kendall/Moran estimator | (Baldwin & Sanderson, 1998) |

**Table S2.** Taxa used in this study, voucher information, and GenBank accession numbers for ITS, *rps16*, and *mat*K.
Herbaria: FI = Museo di Storia Naturale dell'Universita`, Firenze, Italy; GE = Universita` di Genova, Genova, Italy; RNG = University of Reading Herbarium; JACA = Instituto Pirenaico de Ecologı´a, Jaca, Spain; US = United States National Herbarium, Smithsonian Institution; S = Swedish Museum of Natural History, Stockholm, Sweden; BM = British Museum of Natural History; GB = University of Gothenburg herbarium; VPI = Massey Herbarium, Virginia Polytechnic Institute and State University; UPS = Museum of Evolution, Uppsala University; IBSC = South China Botanical Garden; B = The herbarium of the Botanic Garden and Botanical Museum Berlin-Dahlem; BOCH = Ruhr-Universität Bochum; L = Leiden herbarium; HMK = Herbarium Martin Kemler; C = Herbarium Copenhagen; UBT = The University of Beyreuth Herbarium, within the Ecological-Botanical Garden of the University of Bayreuth; AAU = Aarhus University Herbarium; LD = Lund University Botanical Museum Herbarium; MSB = Ludwig-Maximilians-Universität; NY = The New York Botanical Garden; KUN = Kunming Institute of Botany, Chinese Academy of Sciences; M = Botanische Staatssammlung München; K = Royal Botanic Gardens, Kew; HSNU = the herbarium of East China Normal University; SYKO = the Herbarium of the Institute of Biology, Komi Scientific Center; G = Conservatory and Botanical Garden of the City of Geneva; MJG = Johannes Gutenberg-Universität; NMW = National Museum Wales; TUR = University of Turku; YNUH = Yeungnam University; TMRC = Shahid Beheshti University of Medical Sciences; UPA = University of Patras; W = Naturhistorisches Museum Wien.

| | | **matK** | | | ***rps16*** | | | **ITS** | | |
|---|---|---|---|---|---|---|---|---|---|---|
| **species name** | **tribe** | **source** | **voucher informati on** | **GenBank accession number** | **source** | **voucher information (herbarium)** | **accession number** | **source** | **voucher informati on** | **accession number** |
| *Acanthophyllum acerosum* Sosn. | Caryophylleae | - | - | - | GenBank | Zarre & al. 41900 (TUH) | KF924732 | GenBank | Zarre & al. 41900 (TUH) | KF924680 |
| *Acanthophyllum bracteatum* Boiss. | Caryophylleae | - | - | - | GenBank | Pirani & Moazzeni 2104 (TMRC) | KF924726 | GenBank | Pirani & Moazzeni 2104 (TMRC) | KF924674 |
| *Acanthophyllum bungei* (Boiss.) Trautv. | Caryophylleae | - | - | - | GenBank | Nydegger 19519 (MSB) | MF401169 | GenBank | Nydegger 19519 (MSB) | MF401121 |
| *Acanthophyllum caespitosum* Boiss. | Caryophylleae | - | - | - | GenBank | Zarre & al. 41903 (TUH) | KF924725 | GenBank | Zarre & al. 41903 (TUH) | KF924673 |

| | | | | | | | | | | |
|---|---|---|---|---|---|---|---|---|---|---|
| *Acanthophyllum cerastioides* (D.Don) Madhani & Zarre | Caryophylleae | - | - | - | GenBank | Rechinger 30724 (M) | MF401168 | GenBank | Rechinger 30724 (M) | MF401122 |
| *Acanthophyllum honigbergeri* (Fenzl) Barkoudah | Caryophylleae | - | - | - | GenBank | K. H. Rechinger 35371 (B) | MF401176 | GenBank | K. H. Rechinger 35371 (B) | MF401125 |
| *Acanthophyllum kermanense* (Bornm.) A.Pirani & Rabeler | Caryophylleae | GenBank | 100591988 (B) | KX183873 | - | - | - | GenBank | 10059198 8 (B) | KX184037 |
| *Acanthophyllum kurdicum* Boiss. & Hausskn. ex Boiss. | Caryophylleae | - | - | - | GenBank | Hamzehee & Lashkarbolooki 1756 (TARI) | KF924712 | GenBank | Hamzehee & Lashkarbo looki 1756 (TARI) | KF924660 |
| *Acanthophyllum mucronatum* C.A.Mey. | Caryophylleae | - | - | - | GenBank | Optima Iter XI/2050 (M) | MF401170 | GenBank | Assadi & Olfat 68668 (TARI) | KF924652 |
| *Acanthophyllum paniculatum* Regel & Herder | Caryophylleae | GenBank | T.A. Haebkob (NY) | JN589209 | - | - | - | GenBank | T.A. Haebkob s.n. (NY) | JN589016 |
| *Acanthophyllum sordidum* Bunge ex Boiss. | Caryophylleae | GenBank | Kiseleva & Proskuriak ova s.n. (FI) | AY936324 | GenBank | Pirani & Moazzeni 2147 (TMRC) | KF924697 | GenBank | Pirani & Moazzeni 2147 (TMRC) | KF924644 |
| *Acanthophyllum versicolor* Fisch. & C.A.Mey. | Caryophylleae | GenBank | Nydegger 435976 (FI) | AY936323 | GenBank | Nydegger 43597b (MSB) | KF924687 | GenBank | Nydegger 43597b (MSB) | KF924633 |
| *Agrostemma githago* L. | Sileneae | GenBank | Minuto s.n. (GE) | AY936315 | | | | | | |
| *Amaranthus caudatus* L. | Outgroup(Ama ranthaceae Juss.) | GenBank | N/A | AY514809 | | | | | | |
| *Arenaria digyna* Willd. Ex D.F.K.Schltdl. | Arenarieae | GenBank | N/A | AY936304 | | | | | | |

| | | | | | | | | | | |
|---|---|---|---|---|---|---|---|---|---|---|
| *Arenaria nevadensis* Boiss. & Reut. | Arenarieae | GenBank | Quer & Cuartrec s.n. (S) | AY936303 | | | | | | |
| *Arenaria serpyllifolia* L. | Arenarieae | GenBank | Fior s.n. (GE) | AY936302 | | | | | | |
| *Balkana spergulifolia* (Griseb.) Madhani & Zarre | Caryophylleae | - | - | - | GenBank | Kalheber 04-1558 (M) | MF401185 | GenBank | Kalheber 04-1558 (M) | MF401126 |
| *Bolanthus cherlerioides* (Bornm.) Barkoudah | Caryophylleae | - | - | - | GenBank | K.P. & E. Buttler 19986 (M) | MF401183 | GenBank | K.P. & E. Buttler 19986 (M) | MF401128 |
| *Bolanthus confertifolius* ( Hub.-Mor.) Madhani & Heubl | Caryophylleae | - | - | - | - | - | - | GenBank | Özkan Eren 4362 (B) | KX834007 |
| *Bolanthus filicaulis* (Boiss.) Barkoudah | Caryophylleae | - | - | - | GenBank | Fragman-Sapir s.n. (UPA) | MN995821 | GenBank | Fragman-Sapir s.n. (UPA) | MN990420 |
| *Bolanthus frankenioides* (Boiss.) Barkoudah | Caryophylleae | - | - | - | - | - | - | GenBank | N/A | KY406148 |
| *Bolanthus fruticulosus* (Bory & Chaub.) Barkoudah | Caryophylleae | - | - | - | GenBank | Rechinger 19439 (M) | MF401180 | - | - | - |
| *Bolanthus graecus* (Schreb.) Barkoudah | Caryophylleae | - | - | - | GenBank | Merxmüller & Podlech 31173 (MSB) | MF401181 | GenBank | Merxmüller & Podlech 31173 (MSB) | KX834004 |
| *Bolanthus hirsutus* (Labill.) Barkoudah | Caryophylleae | - | - | - | GenBank | Fragman-Sapir s.n. (UPA) | MT063100 | GenBank | Fragman-Sapir s.n. (UPA) | MN990421 |
| *Bolanthus huber-morathii* C.Simon | Caryophylleae | - | - | - | GenBank | Nydegger 15138 (MSB) | MF401184 | GenBank | Nydegger 15138 (MSB) | KX834006 |
| *Bolanthus laconicus* (Boiss.) Barkoudah | Caryophylleae | - | - | - | GenBank | Zografidis 579 (UPA) | MN995820 | - | - | - |
| *Bolanthus mevlanae* Aytaç | Caryophylleae | - | - | - | - | - | - | GenBank | N/A | KY406145 |

| | | | | | | | | | | |
|---|---|---|---|---|---|---|---|---|---|---|
| *Bolanthus minuartioides* (Jaub. & Spach) Hub.-Mor. | Caryophylleae | - | - | - | - | - | - | GenBank | Walter 201 (B) | KX834005 |
| *Bolanthus ortegioides* (Fisch. & C.A.Mey.) Madhani & Rabeler | Caryophylleae | - | - | - | GenBank | Zarre 42 (MSB) | MF401182 | GenBank | Zarre 42 (MSB) | KX834008 |
| *Bolanthus spergulifolius* (Jaub. & Spach) Hub.-Mor. | Caryophylleae | - | - | - | - | - | - | GenBank | N/A | KY406150 |
| *Bolanthus stenopetalus* Hartvig & Å.Strid | Caryophylleae | - | - | - | - | - | - | GenBank | N/A | KY406147 |
| *Bolanthus thymifolius* Sm. (Phitos), | Caryophylleae | - | - | - | - | - | - | GenBank | Constantinidis s.n. (UPA) | MN99042 |
| *Bolanthus thymoides* Hub.-Mor. | Caryophylleae | - | - | - | - | - | - | GenBank | N/A | KY406152 |
| *Bolanthus turcicus* Koç & Hamzaoglu | Caryophylleae | - | - | - | - | - | - | GenBank | N/A | KY406146 |
| *Bufonia macropetala* subsp. *wilkommiana* (Boiss.) Amich | Sagineae | GenBank | Zubizarreta 14916 (FI) | AY936290 | | | | | | |
| *Bufonia tenuifolia* L. | Sagineae | GenBank | Montserrat 4964/69 (FI) | AY936289 | | | | | | |
| *Cerastium arvense* L. | Alsineae | GenBank | Minuto & Fior s.n. (GE) | AY936295 | | | | | | |
| *Cerastium fontanum* Baumg. | Alsineae | GenBank | Alanko 58340 (FI) | AY936296 | | | | | | |
| *Cherleria laricifolia* (L.) lamonico | Sclerantheae | GenBank | Martini s.n., (FI) | AY936294 | | | | | | |
| *Corrigiola litoralis* subsp. *foliosa* (Pérez Lara) Devesa | Corrigioleae | GenBank | Lewalle 10564 (FI) | AY936331 | | | | | | |
| *Cyathophylla chlorifolia* (Poir.) Bocquet & Strid | Caryophylleae | GenBank | 100208060 (B) | KX183864 | GenBank | Ulrich s.n. (M) | MF401164 | GenBank | 100208060 (B) | KX184028 |

| | | | | | | | | | | |
|---|---|---|---|---|---|---|---|---|---|---|
| *Cyathophylla viscosa* (C.A.Mey.) Madhani & Rabeler | Caryophylleae | GenBank | 254384 (L) | KX183924 | GenBank | Optima Iter XI/1846 (M) | MF401165 | GenBank | Optima Iter XI/1846 (M) | MF401117 |
| *Dianthus andrzejowskianus* (Zapal.) Kulcz. | Caryophylleae | GenBank | V. Tichomirov 7462 (NY) | JN589251 | - | - | - | GenBank | V. Tichomirov 7462 (NY) | JN589032 |
| *Dianthus armeria* L. | Caryophylleae | GenBank | R. Rabeler 1431 (MICH) | FJ404832 | GenBank | R. Rabeler 1431 (MICH) | FJ404903 | GenBank | Steven R. Hill 32454 (NY) | JN589087 |
| *Dianthus barbatus* L. | Caryophylleae | GenBank | C. Aedo et al. 2201 (NY) | JN589237 | - | - | - | GenBank | Valente, LM 314LV07 | GU440783 |
| *Dianthus benearnensis* Loret | Caryophylleae | GenBank | N/A | KX183892 | - | - | - | GenBank | Morales & al 485RM (MA 45883) | GU440784 |
| *Dianthus broteri* Boiss. & Reut. | Caryophylleae | GenBank | A. Schinini, G. Aviles Olmos, & S. Sanchez Garcia 32461 (NY) | JN589180 | - | - | - | GenBank | A. Schinini, G. Aviles Olmos, & S. Sanchez Garcia 32461 (NY) | JN589117 |
| *Dianthus candicus* (P.W.Ball & Heywood) Madhani & Heubl | Caryophylleae | - | - | - | GenBank | Greuter 7679 (M) | MF401178 | - | - | - |
| *Dianthus capitatus* J.St.-Hil. | Caryophylleae | GenBank | E. Marin 633 (NY) | JN589204 | - | - | - | GenBank | Vargas, P. et al 147PV06 | GU440792 |
| *Dianthus carthusianorum* L. | Caryophylleae | GenBank | R.K. Rabeler 710 (NY) | JN589238 | GenBank | N/A | EF674194 | GenBank | Aldasoro 8729 (MA 727515) | GU440794 |

| | | | | | | | | | | |
|---|---|---|---|---|---|---|---|---|---|---|
| *Dianthus caryophyllus* L | Caryophylleae | GenBank | F. Casas 598 (NY) | JN589207 | GenBank | N/A | NC039650 | GenBank | F. Casas 598 (NY) | JN589053 |
| *Dianthus caucaseus* Sims | Caryophylleae | GenBank | J. Reveal 8688 (NY) | JN589184 | - | - | - | GenBank | J. Reveal 8688 (NY) | JN589024 |
| *Dianthus chinensis* L. | Caryophylleae | GenBank | H.T. Beck 1414 (NY) | JN589242 | - | - | - | GenBank | Kim, 2005-0815 | GU440796 |
| *Dianthus cretaceus* Adams | Caryophylleae | GenBank | O. Abdaladze, M. Chiboshvili, & T. Siukaev 295 (NY) | JN589232 | - | - | - | GenBank | Aedo & al 11669 (MA 743655) | GU440804 |
| *Dianthus crinitus* Sm. | Caryophylleae | GenBank | Atha, et al. 3157 (NY) | JN589188 | - | - | - | GenBank | Muñoz Garmendia & al. 4661 (MA 688938) | GU440806 |
| *Dianthus cyri* Fisch. & C.A.Mey. | Caryophylleae | GenBank | T. Gviniashvili, L. Jinjolia, 458, NY | JN589246 | - | - | - | GenBank | Medina et al 2462 (MA742743) | GU440808 |
| *Dianthus deltoides* L. | Caryophylleae | GenBank | Steven R. Hill 32613 (NY) | JN589299 | - | - | - | GenBank | Christenhusz MJM 4326 (TUR) | GU440809 |
| *Dianthus furcatus* Balb. | Caryophylleae | GenBank | Minuto & Fior s.n. (GE) | AY936320 | - | - | - | GenBank | Minuto & Fior s.n. (GE) | AY936268 |
| *Dianthus gracilis* Sm. | Caryophylleae | GenBank | T. Constantinidis 10216 (NY) | JN589276 | - | - | - | GenBank | Wilkinson, S. SW83 | GU440822 |
| *Dianthus gratianopolitanus* Vill. | Caryophylleae | GenBank | Faulconer 42 | MF349921 | - | - | - | GenBank | UNKAR3620 | KU722885 |

| | | | | | | | | | | |
|---|---|---|---|---|---|---|---|---|---|---|
| *Dianthus haematocalyx* Boiss. & Heldr. | Caryophylleae | GenBank | E. Horandl & F.F. Hadacek 7664 (NY) | JN589197 | - | - | - | GenBank | Vargas, P. 87PV08 | GU440824 |
| *Dianthus illyricus* (Ard.) Fassou, N.Korotkova, Dimop. & Borsch | Caryophylleae | GenBank | 272414 (L) | KX183925 | - | - | - | GenBank | 170952 (L) | KX184016 |
| *Dianthus imereticus* (Rupr.) Schischk. | Caryophylleae | GenBank | M. Khutsishvili 751 (NY) | JN589168 | - | - | - | GenBank | A. Groeger & W. Lobin UNKAR7911 | KU722886 |
| *Dianthus inamoenus* Schischk. | Caryophylleae | GenBank | N. Lachashvili & M. Khutsishvili 439 (NY) | JN589179 | - | - | - | - | - | - |
| *Dianthus knappii* (Pant.) Asch. & Kanitz ex Borbás | Caryophylleae | GenBank | T.G. Lammers 8600 (NY) | JN589284 | - | - | - | GenBank | Christenhusz 4316 TUR | GU440828 |
| *Dianthus kuschakewiczii* Regel & Schmalh. | Caryophylleae | GenBank | N/A | MF158641 | - | - | - | - | - | - |
| *Dianthus longicalyx* Miq. | Caryophylleae | GenBank | UNKAR8506 | KU722873 | GenBank | N/A | KM668208 | GenBank | 20100029 (YNUH) | KF954520 |
| *Dianthus lusitanus* Brot. | Caryophylleae | GenBank | W. Lippert 25368 (NY) | JN589224 | - | - | - | GenBank | S. Wilkinson SW5 | GU440838 |
| *Dianthus microlepis* Boiss. | Caryophylleae | GenBank | P.M. Uribe-Echebarria 226 (NY) | JN589171 | - | - | - | GenBank | P.M. Uribe-Echebarria 226 (NY) | JN589034 |
| *Dianthus micropetalus* Ser. | Caryophylleae | - | - | - | - | - | - | GenBank | Retief en Germishuizen 231 K | GU440841 |

| | | | | | | | | | | |
|---|---|---|---|---|---|---|---|---|---|---|
| *Dianthus monspessulanus* L. | Caryophylleae | GenBank | E. Horandl & F.F. Hadacek 6028 (NY) | JN589181 | - | - | - | GenBank | N/A | KC293977 |
| *Dianthus moravicus* Kovanda | Caryophylleae | GenBank | N/A | LN877396 | GenBank | N/A | LN877396 | GenBank | N/A | KC293978 |
| *Dianthus nudiflorus* Griff. | Caryophylleae | GenBank | H.M.P. N. Rec.It. 1302 (FI) | AY936322 | - | - | - | GenBank | H.M.P. N. Rec.It. 1302 (FI) | AY936269 |
| *Dianthus orientalis* Adams | Caryophylleae | GenBank | A.A. Donmez 11044 (NY) | JN589265 | - | - | - | GenBank | Nisa et al 1023(MA 689355) | GU440847 |
| *Dianthus recticaulis* Ledeb. | Caryophylleae | - | - | - | GenBank | Fayvush & al. OPTIMA Iter XI/2199 (M) | MF401177 | - | - | - |
| *Dianthus repens* Willd. | Caryophylleae | GenBank | N.A. Brummitt & R.K. Brummitt 174 (NY) | JN589229 | - | - | - | GenBank | N.A. Brummitt & R.K. Brummitt 174 (NY) | JN589124 |
| *Dianthus seguieri* Vill. | Caryophylleae | GenBank | A. Gharpin & R. Salanan AC19645 (NY) | JN589247 | - | - | - | GenBank | Molero J & Vicens J 3ITDG7878 (MA 632669) | GU440856 |
| *Dianthus serotinus* Waldst. & Kit. | Caryophylleae | GenBank | J. Walter 7975 (NY) | JN589269 | - | - | - | GenBank | Christenhusz 4324 (TUR) | GU440857 |
| *Dianthus serrulatus* Desf. | Caryophylleae | - | - | - | - | - | - | GenBank | Lafkih et al 695 (Reading 19 2006 60) | GU440860 |
| *Dianthus squarrosus* M.Bieb. | Caryophylleae | GenBank | V.A. Sagalaev, V.D. Bochkin, | JN589182 | - | - | - | GenBank | V.A. Sagalaev, V.D. Bochkin, | JN589065 |

| | | | | | | | | | | |
|---|---|---|---|---|---|---|---|---|---|---|
| | | | & N.G. Il’minskikh 1993 (NY) | | | | | | & N.G. Il’minskikh 1993 (NY) | |
| *Dianthus subulosus* Conrath & Freyn | Caryophylleae | GenBank | D.E. Atha, H. Stevens, & M. Khutsishvili 2782 (NY) | JN589234 | - | - | - | GenBank | Tribsch, 11172 | GU440865 |
| *Dianthus superbus* L. | Caryophylleae | GenBank | D.E. Boufford, J.H. Chen, K. Fujikawa, S.L. Kelley, R.H. Ree, H. Sun, J.P. Yue, D.C. Zhang, & Y.H. Zhang 34287 (NY) | JN589216 | - | - | - | GenBank | RBG Edinburgh, Living Coll., FBI 34 | GU440867 |
| *Dianthus sylvestris* Wulfen | Caryophylleae | GenBank | MIB:ZPL: 03122 | HE967400 | - | - | - | GenBank | P. Vargas 120PV08 | GU440870 |
| *Dianthus tunicoides* (Ser.) Madhani & Heubl | Caryophylleae | - | - | - | GenBank | Lüdtke 581 (M) | MF401179 | GenBank | Lüdtke 581 (M) | MF401129 |
| *Dianthus turkestanicus* Preobr. | Caryophylleae | GenBank | UNKAR3 671 | KU722876 | GenBank | ZXQ-30 | KX158434 | GenBank | ZXQ-30 | KX158323 |
| *Drypis spinosa* L | Sagineae | GenBank | Moggi, Luccioli & Tosi s.n. (FI) | AY936293 | - | - | - | | - | - |
| *Eremogone picta* (Sm.) Dillenb. & Kadereit | Eremogoneae | GenBank | H.M.P. N.Rec.It. | AY936319 | - | - | - | - | - | - |

| | | | | | | | | | | |
|---|---|---|---|---|---|---|---|---|---|---|
| | | | 1587/91 (FI) | | | | | | | |
| *Eremogone procera* (Spreng.) Rchb. | Eremogoneae | GenBank | Bochkin, Klinkova & Sogalaev s.n. (FI) | AY936318 | - | - | - | - | - | - |
| *Eremogone saxatilis* (L.) Ikonn. | Eremogoneae | GenBank | Tichomirov & Anoheeva s.n. (FI) | AY936317 | - | - | - | - | - | - |
| *Facchinia lanceolata* (All.) Rchb. | Sagineae | GenBank | Pascale s.n. (FI) | AY936292 | - | - | - | - | - | - |
| *Gypsophila acantholimoides* Bornm. | Caryophylleae | - | - | - | This study | S. Zarre & H. Madhani 34290 (TUH) | OR801022* | This study | S. Zarre & H. Madhani 34290 (TUH) | OR731872* |
| *Gypsophila acutifolia* Fisch. | Caryophylleae | - | - | - | GenBank | Quasdorf 67 (B) | MF401156 | GenBank | Quasdorf 67 (B) | MF401100 |
| Gypsophila sp. | Caryophylleae | - | - | - | This study | H. Madhani 34298 (TUH) | OR801032* | This study | H. Madhani 34298 (TUH) | OR735441* |
| *Gypsophila altissima* L. | Caryophylleae | GenBank | M. Chase 8852 (K) | AY042597 | This study | A. Skvortsx s.n. (M) | OR801037* | This study | A. Skvortsx s.n. (M) | OR735440* |
| *Gypsophila arabica* Barkoudah | Caryophylleae | - | - | - | - | - | - | This study | A. Danin et al. 35.036?! (B) | MF401082 |
| *Gypsophila aretioides* Boiss. | Caryophylleae | - | - | - | This study | H. Madhani & N. Madhani 47116 (TUH) | OR801028* | This study | H. Madhani & N. Madhani 47116 (TUH) | OR735460* |

| | | | | | | | | | | |
|---|---|---|---|---|---|---|---|---|---|---|
| *Gypsophila arrostii* Guss. | Caryophylleae | GenBank | C. Aedo et al. 15625SC (NY) | JN589167 | GenBank | Nydegger 13410 (MSB) | MF401155 | This study | Nydegger 13410 (MSB) | OR735452* |
| *Gypsophila aucheri* Boiss. | Caryophylleae | GenBank | A.A. Donmez & H. Ascantas 4871 (NY) | JN589200 | GenBank | Nydegger 18633 (MSB) | MF401147 | GenBank | Nydegger 18633 (MSB) | MF401098 |
| *Gypsophila bazorganica* Rech.f. | Caryophylleae | - | - | - | This study | H. Madhani 45392 (TUH) | OR801026* | This study | H. Madhani 45392 (TUH) | OR735457* |
| *Gypsophila bermejoi* G.López | Caryophylleae | - | - | - | - | - | - | GenBank | Ladero & Casaseca 12107 (B) | MF401106 |
| *Gypsophila bicolor* (Freyn & Sint.) Grossh. | Caryophylleae | GenBank | N. Lachashvili 533 (NY) | JN589208 | GenBank | Zarre, Mashayekhi, Taeb, Pirani & Moazzeni 35136 (MSB) | MF401149 | This study | Zarre, Mashayekhi, Taeb, Pirani & Moazzeni 35136 (MSB) | OR735454* |
| *Gypsophila bitlisensis* Barkoudah | Caryophylleae | - | - | - | - | - | - | This study | Raus 4199 (B) | OR735464* |
| *Gypsophila bucharica* B.Fedtsch. | Caryophylleae | GenBank | G. Kinzikaeva & T. Koczkareva 1988 ( NY) | JN589190 | GenBank | Kinzikaeva & Koczkareva 6663 (M) | MF401162 | GenBank | Kinzikaeva & Koczkareva 6663 (M) | MF401102 |
| *Gypsophila capillaris* (Forssk.) C.Chr. | Caryophylleae | - | - | - | GenBank | Podlech 50067 (MSB) | MF401135 | GenBank | Podlech 50067 (MSB) | MF401092 |
| *Gypsophila capitata* M.Bieb. | Caryophylleae | - | - | - | GenBank | Tzvelev, Czerepanov, Bobrov & Dogadova 7559 (B) | MF401161 | GenBank | Tzvelev, Czerepanov, Bobrov & Dogadova 7559 (B) | MF401103 |

| | | | | | | | | | | |
|---|---|---|---|---|---|---|---|---|---|---|
| *Gypsophila capituliflora* Rupr. | Caryophylleae | GenBank | S. Ikonnikov 4365 (NY) | JN589211 | GenBank | Ikonnikov 4365 (M) | MF401157 | GenBank | Ikonnikov 4365 (M) | MF401111 |
| *Gypsophila caricifolia* Boiss. | Caryophylleae | - | - | - | This study | H. Madhani 34294 (TUH) | OR801023 | This study | H. Madhani 34294 (TUH) | OR735459* |
| Gypsophila *cephalotes* (Schrenk) F.N.Williams | Caryophylleae | GenBank | N. Kaletkina 6662 (NY) | JN589176 | GenBank | Anders 7613 (MSB) | MF401158 | GenBank | Anders 7613 (MSB) | MF401105 |
| *Gypsophila curvifolia* Fenzl | Caryophylleae | GenBank | A.A. Donmez, B Mutlu and T. Agar 13674 (NY) | JN589186 | GenBank | P. Hein 52-2 (B) | MF401159 | GenBank | P. Hein 52-2 (B) | MF401099 |
| *Gypsophila davurica* Turcz. ex Fenzl | Caryophylleae | GenBank | N/A | MK534848 | - | - | - | GenBank | N/A | MK555271 |
| *Gypsophila elegans* M.Bieb. | Caryophylleae | GenBank | Hillyihugseanov s.n. (FI) | AY936327 | This study | H. Madhani 45395 (TUH) | OR801018* | This study | H. Madhani 45395 (TUH) | OR735465* |
| *Gypsophila eriocalyx* Boiss. | Caryophylleae | - | - | - | This study | Nydegger 43821 (MSB) | OR801007* | GenBank | N/A | OR396946 |
| *Gypsophila fastigiata* L. | Caryophylleae | GenBank | J. Walter 116 (NY) | JN589262 | GenBank | Kalheber 88-2892 (M) | MF401152 | GenBank | Kalheber 88-2892 (M) | MF401097 |
| *Gypsophila globulosa* Steven. | Caryophylleae | - | | - | - | - | - | GenBank | Köhler (61) Bm 4306210 (B) | MF401108 |
| *Gypsophila graminifolia* Barkoudah | Caryophylleae | - | - | - | This study | H. Madhani 45397 (TUH) | OR801027* | This study | H. Madhani 45397 (TUH) | OR735458* |
| *Gypsophila gypsophiloides* (Fenzl) Blakelock | Caryophylleae | - | - | - | GenBank | Rechinger 48149 (M) | MF401138 | GenBank | Rechinger 48149 (M) | MF401086 |

| | | | | | | | | | | |
|---|---|---|---|---|---|---|---|---|---|---|
| *Gypsophila heteropoda* Freyn | Caryophylleae | GenBank | V. Vasak 1985 (NY) | JN589302 | - | - | - | GenBank | V. Vasak s.n. (B) | MF401085 |
| *Gypsophila huashanensis* Tsui & D.Q.Lu | Caryophylleae | GenBank | N/A | MK435786 | - | - | - | GenBank | TianXH033 | MH808292 |
| *Gypsophila laricina* Schreb. | Caryophylleae | - | - | - | This study | F. Sorger 77-101-4 (M) | OR801038* | This study | F. Sorger 77-101-4 (M) | OR735435* |
| *Gypsophila leioclada* Rech.f. | Caryophylleae | - | - | - | This study | H. Madhani 34293 (TUH) | OR801014* | GenBank | Podlech & Zarre 55219 (MSB) | MF401104 |
| *Gypsophila libanotica* Boiss. | Caryophylleae | - | - | - | This study | Güven Görk, Per Hartvig & Arne strid 10 0630519 (B) | OR801036* | This study | Güven Görk, Per Hartvig & Arne strid 10 0630519 (B) | OR735436* |
| *Gypsophila linearifolia* (Fisch. & C.A.Mey.) Boiss. | Caryophylleae | - | - | - | This study | R. Hand & C. S. Christodoulou 5454 (B) | OR801021* | GenBank | Akhani 8509 (MSB) | MF401091 |
| *Gypsophila melampoda* Bien. ex Boiss. | Caryophylleae | - | - | - | This study | S. Zarre & H. Madhani s.n. (TUH) | OR801012* | This study | S. Zarre & H. Madhani s.n. (TUH) | OR735446* |
| *Gypsophila nabelekii* Schischk. | Caryophylleae | - | - | - | GenBank | K. H. Rechinger 48847 (B) | MF401142 | GenBank | K. H. Rechinger 48847 (B) | MF401088 |
| *Gypsophila nana* Bory & Chaub. | Caryophylleae | - | - | - | This study | N. Bohling 8749 (B) | OR801010* | This study | N. Bohling 8749 (B) | OR735450* |
| *Gypsophila nodiflora* (Boiss.) Barkoudah | Caryophylleae | - | | - | - | - | - | This study | Davis & Hedge D.28883 (B) | OR735445* |
| *Gypsophila oblanceolata* Barkoudah | Caryophylleae | - | - | - | GenBank | Hagemann, Binder & | MF401160 | GenBank | Hagemann, Binder & | MF401115 |

| | | | | | | | | | | |
|---|---|---|---|---|---|---|---|---|---|---|
| | | | | | | Schwarz 2144 (B) | | | Schwarz 2144 (B) | |
| *Gypsophila oldhamiana* Miq. | Caryophylleae | GenBank | s.n. 20151013 (HSNU) | MF064036 | - | - | - | GenBank | 20151013 (HSNU) | MF063499 |
| *Gypsophila pacifica* Kom. | Caryophylleae | - | - | - | - | - | - | GenBank | N/A | JX274528 |
| *Gypsophila pallasii* Ikonn. (G. glomerata Pall. ex M.Bieb) | Caryophylleae | - | - | - | - | - | - | This study | V. Vasak 10 0630523 (B) | OR735439* |
| *Gypsophila pallida* Stapf | Caryophylleae | - | - | - | - | - | - | This study | K. H. Rechinger , 48028 (B) | OR735444* |
| *Gypsophila paniculata* L. | Caryophylleae | GenBank | H. Hess, K. Allen, N. Stoynoff, K. Cherwin, & K. Dreisilker 9201 (NY) | JN589231 | This study | H. Merxmüller & W. Sauer 31544 (M) | OR801030* | This study | H. Merxmüller & W. Sauer 31544 (M) | OR735451* |
| *Gypsophila patrinii* Ser. | Caryophylleae | GenBank | A. Cronquist 12157 (NY) | JN589271 | GenBank | Raab-Straube 020105 (B) | MF401150 | GenBank | Raab-Straube 020105 (B) | MF401110 |
| *Gypsophila perfoliata* L. | Caryophylleae | GenBank | ZXQ-31 (?) | KX158361 | GenBank | Doring, Parolly & Tolimir 7438 (B) | MF401139 | GenBank | Doring, Parolly & Tolimir 7438 (B) | MF401114 |
| *Gypsophila persica* Barkoudah | Caryophylleae | - | - | - | This study | K. H. Rechinger 47133 (B) | OR801029* | This study | K. H. Rechinger 47133 (B) | OR735447* |
| *Gypsophila sp.* | Caryophylleae | - | - | - | This study | H. Madhani 34292 (TUH) | OR801009* | This study | H. Madhani 34292 (TUH) | OR735449* |

| | | | | | | | | | | |
|---|---|---|---|---|---|---|---|---|---|---|
| *Gypsophila petraea* (Baumg.) Rchb. | Caryophylleae | - | - | - | GenBank | Buttler & Dietrich 8953 (B) | MF401151 | GenBank | Buttler & Dietrich 8953 (B) | MF401095 |
| *Gypsophila pilosa* Huds. | Caryophylleae | - | - | - | This study | S. Zarre, H. Madhani 34287 (TUH) | OR801019* | GenBank | S. Zarre, H. Madhani 34287 (TUH) | MF401094 |
| *Gypsophila pilulifera* Boiss. & Heldr. | Caryophylleae | GenBank | A.A. Donmez, B Mutlu, & T. Agar 13754 (NY) | JN589173 | This study | S. Zarre 103 (MSB) | OR801015* | This study | S. Zarre 103 (MSB) | OR735453* |
| *Gypsophila pinifolia* Boiss. & Hausskn. | Caryophylleae | GenBank | C. Aedo et al. 4668 (NY) | JN589288 | GenBank | Buttler 5774 (M) | MF401163 | GenBank | Buttler 5774 (M) | MF401116 |
| *Gypsophila polyclada* Fenzl ex Boiss. | Caryophylleae | - | - | - | This study | H. Madhani 45399 (TUH) | OR801013* | This study | H. Madhani 45399 (TUH) | OR735448* |
| *Gypsophila pulvinaris* Rech.f. | Caryophylleae | - | - | - | This study | H. Madhani 34297 (TUH) | OR801025* | This study | H. Madhani 34297 (TUH) | OR735455* |
| *Gypsophila repens* L. | Caryophylleae | GenBank | T. Flynn 6777 (NY) | JN589281 | GenBank | Podlech 38401 (MSB) | MF401153 | GenBank | Podlech 38401 (MSB) | MF401101 |
| *Gypsophila ruscifolia* Boiss. | Caryophylleae | - | - | - | This study | H. Madhani 34295 (TUH) | OR801034* | GenBank | H. Madhani 34295 (TUH) | OR735443* |
| *Gypsophila saponarioides* Bornm. & Gauba | Caryophylleae | - | - | - | - | - | - | This study | H. Madhani & N. Madhani 47425 (TUH) | OR735461* |

| | | | | | | | | | | |
|---|---|---|---|---|---|---|---|---|---|---|
| *Gypsophila scorzonerifolia* Ser. | Caryophylleae | GenBank | A.W. Cusick 33241 (NY) | JN589195 | - | - | - | GenBank | A.W. Cusick 33241 (NY) | JN589100 |
| *Gypsophila silenoides* Rupr. | Caryophylleae | GenBank | L.H. Bailey Hortorium 11239 (NY) | JN589290 | This study | Fayvush et al., OPTIMA Iter XI/0330 (M) | OR801017* | This study | Fayvush et al., OPTIMA Iter XI/0330 (M) | OR735463* |
| *Gypsophila simulatrix* Bornm. & Woronow | Caryophylleae | - | - | - | This study | M. Nydegger 19307 (MSB) | OR801035* | This study | M. Nydegger 19307 (MSB) | OR735442* |
| *Gypsophila sp. (ortegioiges!)* | Caryophylleae | - | - | - | - | - | - | GenBank | Manissadjian 1165 (B) | MF401087 |
| *Gypsophila stevenii* Fisch. Ex Schrank | Caryophylleae | GenBank | N. Lachashvili 488 (NY) | JN589273 | - | - | - | GenBank | N. Lachashvili 488 (NY) | JN589022 |
| *Gypsophila struthium* subs. *hispanica* (Willk.) G.López (G. hispanica) | Caryophylleae | - | - | - | - | P. Montserrat 10 0630517 (B) | OR801016* | This study | P. Montserrat 10 0630517 (B) | OR735438* |
| *Gypsophila szovitsii* Fisch. & C.A.Mey. ex Fenzl | Caryophylleae | - | - | - | This study | A. Yzossheim 7487 (B) | OR801031* | - | - | - |
| *Gypsophila takhtadzhanii* Schischk. Ex Ikonn. | Caryophylleae | GenBank | A. Takhtadjan & S. Czerepanov 7563 (NY) | JN589289 | - | - | - | - | - | - |
| *Gypsophila tenuifolia* M.Bieb. | Caryophylleae | - | - | - | This study | H. Ch. Friedrich s.n. (M) | OR801020* | This study | H. Ch. Friedrich s.n. (M) | OR735462* |

| | | | | | | | | | | |
|---|---|---|---|---|---|---|---|---|---|---|
| *Gypsophila tomentosa* L. | Caryophylleae | - | - | - | GenBank | Molero 30SWJ90 (33) (M) | MF401146 | GenBank | Molero 30SWJ90 (33) (M) | MF401113 |
| *Gypsophila tuberculosa* Hub.-Mor. | Caryophylleae | - | - | - | This study | Nydegger 19242 (MSB) | OR801008* | This study | Nydegger 19242 (MSB) | OR735437* |
| *Gypsophila uralensis* Less. | Caryophylleae | - | - | - | - | - | - | GenBank | s.n. (SYKO) | KJ755071 |
| *Gypsophila vaccaria* (L.) Sm. | Caryophylleae | GenBank | H.M.P. N.Rec. It. 0267 (FI) | AY936328 | GenBank | Liujq-fjj-0060 (KUN) | MK397906 | GenBank | A. Debolt 1462 (NY) | JN589056 |
| *Gypsophila venusta* Fenzl | Caryophylleae | - | - | - | GenBank | Nydegger 16995 (B) | MF401154 | GenBank | Nydegger 16995 (B) | MF401096 |
| *Gypsophila virgata* Boiss. | Caryophylleae | - | - | - | This study | K. H. Rechinger 43614 (B) | OR801033* | GenBank | K. H. Rechinger 43614 (B) | MF401107 |
| *Gypsophila sp.* | Caryophylleae | - | - | - | This study | H. Madhani 34291 (TUH) | OR801011* | This study | H. Madhani 34291 (TUH) | ?????? (114- G. virgata2) |
| *Gypsophila viscosa* Murray | Caryophylleae | - | - | - | GenBank | Eren Bircan & Gerald Parolly 110 (B) | MF401137 | GenBank | Eren Bircan & Gerald Parolly 110 (B) | MF401084 |
| *Gypsophila wilhelminae* Rech.f. | Caryophylleae | - | - | - | This study | H. Madhani 45393 (TUH) | OR801024 | This study | H. Madhani 45393 (TUH) | OR735456* |
| *Heterochroa desertorum* Bunge | Caryophylleae | GenBank | S. Timokhina & L. Daniljuk 6371 (NY) | JN589275 | GenBank | Timokhina & Daniljuk 6371 (M) | MF401171 | GenBank | Timokhina & Daniljuk 6371 (M) | MF401118 |
| *Heterochroa violacea* Fenzl | Caryophylleae | GenBank | T.S. Elias & D. Murray 11471 (NY) | JN589294 | - | - | - | GenBank | T.S. Elias & D. Murray 11471 (NY) | JN589068 |

| | | | | | | | | | | |
|---|---|---|---|---|---|---|---|---|---|---|
| *Holosteum umbellatum* L. | Alsineae | GenBank | Hoste 8 (FI) | AY936297 | | | | | | |
| *Loeflingia hispanica* L. | Polycarpaeae | GenBank | H.M.P. N.Rec.It. 1121/88 (FI) | AY936288 | | | | | | |
| *Mcneillia graminifolia* (Ard.) Dillenb. & Kadereit | Eremogoneae | GenBank | Skvortsov & Schanzer s.n. (FI) | AY936316 | | | | | | |
| *Minuartia dichotoma* L. | Sagineae | GenBank | Mejías & Silvestre SS15/93 (RNG) | KF737645 | | | | | | |
| *Moehringia muscosa* L | Arenarieae | GenBank | Minuto & Casazza s.n. (GE) | AY936306 | | | | | | |
| *Mollugo verticillata* L. | for rooting the tree (Molluginaceae Bartl.) | GenBank | Barchiesi 0202 (FI) | AY936330 | | | | | | |
| *Odontostemma Pogonanthum* (W.W.Sm.) Sadeghian & Zarre | Alsineae | GenBank | Forrest 27217 (S) | AY936300 | | | | | | |
| *Odontostemma trichophorum* (Franch.) Sadeghian & Zarre | Alsineae | GenBank | Maire 6687 (S) | AY936301 | | | | | | |
| *Ortegia hispanica* L. | Polycarpaeae | GenBank | Valdes & Bermejo s.n. (FI) | AY936286 | | | | | | |
| *Paronychia echinulata* Chater | Paronychieae | GenBank | Selvi s.n. (FI) | AY936285 | | | | | | |
| *Paronychia kapela* A.Kern. | Paronychieae | GenBank | Minuto & Fior s.n. (GE) | AY936284 | | | | | | |
| *Petroana montserratii* (Fern.Casas) Madhani & Zarre | Caryophylleae | GenBank | A. Ortiz 1978 (NY) | JN589280 | GenBank | Casas s.n. (B) | MF401166 | GenBank | Casas s.n. (B) | MF401120 |

| | | | | | | | | | | |
|---|---|---|---|---|---|---|---|---|---|---|
| *Petrocoptis pyrenaica* (Bergeret) A.Braun ex Walp | Sileneae | GenBank | Montserrat & Villar s.n. (JACA) | AY936314 | | | | | | |
| *Petrorhagia dubia* (Raf.) G.López & Romo | Caryophylleae | - | - | - | - | - | - | GenBank | KEW ID 8853 | AY857974 |
| *Petrorhagia nanteuilii* (Burnat) P.W.Ball & Heywood | Caryophylleae | GenBank | 7427 (NMW) | MK925686 | - | - | - | GenBank | 7429 (NMW) | KX165484 |
| *Petrorhagia prolifera* (L.) P.W.Ball & Heywood | Caryophylleae | GenBank | N/A | KJ746192 | - | - | - | GenBank | N/A | GU440883 |
| *Petrorhagia saxifraga* (L.) Link | Caryophylleae | GenBank | L. Faivre 00199087 (G) | KJ204516 | GenBank | S. Downie 1069 (ILL) | FJ404930 | GenBank | Moore 1068 (MJG 011028) | KF737440 |
| *Petrorhagia thessala* (Boiss.) P.W.Ball & Heywood | Caryophylleae | - | - | - | - | - | - | GenBank | N/A | GU440885 |
| *Polycarpon tetraphyllum* (L.) L. | Polycarpaeae | GenBank | Minuto s.n. (GE) | AY936287 | | | | | | |
| *Psammophiliella muralis* (L.) Ikonn. | Caryophylleae | GenBank | F. Kummert 1996 (NY) | JN589218 | GenBank | E. Dörr s.n. (M) | MF401186 | GenBank | E. Dörr s.n. (M) | MF401127 |
| *Psammosilene tunicoides* W.C. Wu & C.Y. Wu | Caryophylleae | GenBank | D.E. Boufford, B. Bartholomew, C.Y. Chen, M.J. Donoghue, R.H. Ree, H. Sun, S.K. Wu 28888 (NY) | JN589222 | GenBank | Gang Yao YGYN2015071101(IBSC) | MN136196 | GenBank | D.E. Boufford, B. Bartholomew, C.Y. Chen, M.J. Donoghue, R.H. Ree, H. Sun, S.K. Wu 28888 (NY) | JN589122 |

| | | | | | | | | | | |
|---|---|---|---|---|---|---|---|---|---|---|
| *Rabelera holostea* (L.) M.T.Sharples & E.A.Tripp | Alsineae | GenBank | N/A | JN589257 | GenBank | D. Podlech 05-1265 (MSB) | MH243549 | - | - | - |
| *Rhodalsine geniculata* (Poir.) F.N.Williams | Sperguleae | GenBank | H.M.P. N.Rec.It. 151/88b (FI) | AY936307 | | | | | | |
| *Sagina pilifera* (DC.) Fenzl | Sagineae | GenBank | Foggi s.n. (FI) | AY936291 | | | | | | |
| *Sagina procumbens* L. | Sagineae | GenBank | H. Schaefer 2008/375 (BM) | HM850782 | | | | | | |
| *Saponaria atocioides* Boiss. | Caryophylleae | GenBank | year 1862 (C) | KX183847 | - | - | - | GenBank | year 1862 (C) | KX183962 |
| *Saponaria bellidifolia* Sm. | Caryophylleae | GenBank | 20006 (BOCH) | KX183901 | - | - | - | GenBank | 20006 (BOCH) | KX183981 |
| *Saponaria bodeana* Boiss. | Caryophylleae | GenBank | 621437 (UBT) | KX183852 | - | - | - | GenBank | 621437 (UBT) | KX183967 |
| *Saponaria caespitosa* DC. | Caryophylleae | GenBank | 20004 (BOCH) | KX183898 | - | - | - | GenBank | 20004 (BOCH) | KX183978 |
| *Saponaria calabrica* Guss. | Caryophylleae | GenBank | B100591971 (B) | KX183860 | - | - | - | GenBank | B100591971 (B) | KX184023 |
| *Saponaria cypria* Boiss. | Caryophylleae | GenBank | B100534453 (B) | KX183866 | - | - | - | GenBank | B100534453 (B) | KX184030 |
| *Saponaria glutinosa* M.Bieb. | Caryophylleae | GenBank | B100351591 (B) | KX183868 | GenBank | Meierott and Gregor GR09/143 (M) | MT152166 | GenBank | B100351591 (B) | KX184031 |
| *Saponaria griffithiana* Boiss. | Caryophylleae | GenBank | B100052225 (B) | KX183870 | GenBank | Schloeder & Jacobs 1757 (M) | MF401133 | GenBank | Schloeder & Jacobs 1757 (M) | MF401080 |
| *Saponaria haussknechtii* Simmler | Caryophylleae | GenBank | Botanical garden Bayreuth fresh material | KX183882 | - | - | - | GenBank | Botanical garden Bayreuth fresh material | KX183972 |
| *Saponaria intermedia* Simmler | Caryophylleae | GenBank | B100591985 (B) | KX183869 | - | - | - | GenBank | B100591985 (B) | KX184032 |

| | | | | | | | | | | |
|---|---|---|---|---|---|---|---|---|---|---|
| *Saponaria intricata* Freyn | Caryophylleae | GenBank | 1045254 (LD) | KX183890 | - | - | - | GenBank | 1045254 (LD) | KX183973 |
| *Saponaria kotschyi* Boiss. | Caryophylleae | GenBank | B1003888 18 (B) | KX183871 | - | - | - | GenBank | B1003888 18 (B) | KX184035 |
| *Saponaria lutea* L. | Caryophylleae | GenBank | (BOCH) | KX183912 | - | - | - | GenBank | (BOCH) | KX183993 |
| *Saponaria mesogitana* Boiss. | Caryophylleae | GenBank | B1002704 62 (B) | KX183875 | - | - | - | GenBank | B1002704 62 (B) | KX184040 |
| *Saponaria ocymoides* L. | Caryophylleae | GenBank | D. Podlech 22945 (MSB) | MT135047 | GenBank | Šída & Vagnerová 3658 (M) | MF401130 | GenBank | Šída & Vagnerová 3658 (M) | MF401077 |
| *Saponaria officinalis* L. | Caryophylleae | GenBank | Minuto s.n. (GE) | AY936325 | GenBank | Sugawara 2080906 (M) | MF401131 | GenBank | Sugawara 2080906 (M) | MF401078 |
| *Saponaria pamphylica* Boiss. & Heldr. | Caryophylleae | GenBank | B1005918 21 (B) | KX183881 | - | - | - | GenBank | B1005918 21 (B) | KX184045 |
| *Saponaria prostrata* Willd. | Caryophylleae | GenBank | 418584 (L) | KX183922 | GenBank | Zarre 122 (MSB) | MF401132 | GenBank | Zarre 122 (MSB) | MF401079 |
| *Saponaria pumila* Janch. | Caryophylleae | GenBank | 016037 (HMK) | KX183921 | - | - | - | GenBank | 016037 (HMK) | KX184001 |
| *Saponaria pumilio* Boiss. | Caryophylleae | GenBank | 20007 (BOCH) | KX183900 | - | - | - | GenBank | 20007 (BOCH) | KX183980 |
| *Saponaria sicula* Raf. | Caryophylleae | GenBank | B1005919 89 (B) | KX183877 | GenBank | N/A | Z83153 | GenBank | B1005919 89 (B) | KX184041 |
| *Saponaria stenopetala* Rech.f. | Caryophylleae | GenBank | 35882 (AAU) | KX183894 | - | - | - | GenBank | 35882 (AAU) | KX183974 |
| *Saponaria subrosularis* Rech.f. | Caryophylleae | GenBank | 29886 (C) | KX183895 | - | - | - | GenBank | 29886 (C) | KX183975 |
| *Saponaria zapateri* Pau | Caryophylleae | GenBank | s.n 052035 (L) | KX183923 | - | - | - | GenBank | s.n 052035 (L) | X184007 |
| *Schiedea ligustrina* Cham. & Schltdl. | Sclerantheae | GenBank | N/A | DQ907813 | | | | | | |
| *Scleranthus annuus* L. | Sclerantheae | GenBank | W. L. Wagner 6862 (US) | DQ267196 | | | | | | |
| *Silene campanula* Pers. | Sileneae | GenBank | Minuto s.n. (GE) | AY936311 | - | - | - | - | - | - |

| | | | | | | | | | | |
|---|---|---|---|---|---|---|---|---|---|---|
| *Silene flos-jovis* (L.) Greuter & Burdet | Sileneae | GenBank | Minuto & Casazza s.n. (GE) | AY936313 | | | | | | |
| *Silene gallica* L. | Sileneae | GenBank | D. Sloan 002 (VPI) | FJ589528 | GenBank | Bengt Oxelman and Lars Tollsten 603 (GB) | LC423974 | GenBank | Bengt Oxelman 2286 (GB) | X86847 |
| *Silene italica* (L.) Pers. | Sileneae | GenBank | Minuto s.n. (GE) | AY936312 | GenBank | Zeynep Aydın 49 (GB) | LC423805 | GenBank | Eslami 29564 (TUH) | LC424095 |
| *Silene vulgaris* (Moench) Garcke | Sileneae | GenBank | Mats Thulin 5717 (UPS) | FJ376828 | GenBank | Mats Thulin 5717 (UPS) | FN821317 | GenBank | Mats Thulin 5717 (UPS) | FN821149 |
| *Spergula arvensis* L. | Sperguleae | GenBank | Baldini s.n. (FI) | AY936310 | | | | | | |
| *Spergularia marina* (L.) Besser | Sperguleae | GenBank | Minuto s.n. (GE) | AY936309 | | | | | | |
| *Spergularia rubra* (L.) J. Presl & C. Presl | Sperguleae | GenBank | Minuto & Fior s.n. (GE) | AY936308 | | | | | | |
| *Stellaria media* (L.) Vill. | Alsineae | GenBank | Minuto s.n. (GE) | AY936299 | | | | | | |
| *Stellaria nemorum* L. | Alsineae | GenBank | Luccioli & Padovani s.n. (FI) | AY936298 | | | | | | |
| *Yazdana shirkuhensis* A.Pirani & N oroozi | Caryophylleae | - | - | - | GenBank | 4003-1 Jalil Noroozi's collection in W | MN417289 & MK651077 | GenBank | 4003-1 Jalil Noroozi's collection in W | MN381230 & MN381233 |

**Table S3** Age estimates for major clades within Caryophylleae using secondary calibration (**on the Caryophylleae node**) of ***rps16*** and **ITS** datasets by BEAST; Net diversification rate estimates Magallon & Sanderson (2001) under low (ε= 0), medium (ε= 0.5), and high (ε= 0.9) extinction rates (ε) implemented in geiger package in R; speciation and extinction rates estimated by (BAMM).

| ***rps16*** | | | | | | |
|---|---|---|---|---|---|---|
| **Major clades** | **Ages estimates & 95% HPD (Ma)** | **Net diversification rate (r) Magallon & Sanderson (2001)** | | | **BAMM analysis** | |
| | | **r (ε= 0)** | **r (ε= 0.5)** | **r (ε= 0.9)** | **Speciation rate mean (95% HPD)** | **Extinction rate mean (95% HPD)** |
| ***Gypsophila*-stem group** (*Gypsophila* + *Saponaria*) n = 182 | 12.74 (6.07–19.6) | 0.41 | 0.35 | 0.23 | 1.46 (0.91–2.79) | 0.8 (0.10–2.49) |
| ***Gypsophila*-crown group** n = 149 | 5.83 (2.57–9.73) | 0.74 | 0.69 | 0.47 | 1.67 (1.05–2.84) | 0.89 (0.07–2.51) |
| **Core *Gypsophila*** n = 114 | 4.48 (1.93–7.39) | 0.92 | 0.85 | 0.56 | 1.92 (1.29 –2.92) | 0.79 (0.06–2.17) |
| ***Dianthus*-stem group** (*Dianthus* + *Petrorhagia* + *Graecobolanthus*) | 5.49 (2.49–8.83) | 1.07 | 0.94 | 0.64 | 1.69 (0.96 –2.97) | 0.8 (0.03 –2.56) |
| ***Dianthus*-crown + *Velezia*** n = 300 | 4.17 (1.81–6.88) | 1.21 | 1.13 | 0.81 | 1.83 (1.02 –3.11) | 0.84 (0.02 –2.59) |
| ***Saponaria*** n = 30 | 3.21 (0.96–6.09) | 0.85 | 0.76 | 0.41 | 0.95 (0.24 –2.77) | 0.6 (0.01–2.50) |
| ***Acanthophyllum*** n = 100 | 4.99 (1.73–8.85) | 0.78 | 0.72 | 0.46 | 1.35 (0.61–2.85) | 0.7 (0.03–2.52) |
| **ITS** | | | | | | |
| **Major clades** | **Ages estimates & 95% HPD (Ma)** | **Net diversification rate (r) Magallon & Sanderson (2001)** | | | **BAMM analysis** | |
| | | **r (ε= 0)** | **r (ε= 0.5)** | **r (ε= 0.9)** | **Speciation rate mean (95% HPD)** | **Extinction rate mean (95% HPD)** |
| ***Gypsophila*-Stem group** (*Gypsophila* + all genera except *Psammosilene*) n = 182 | 19.10 (12.62–25.65) | 0.34 | 0.30 | 0.22 | 0.84 (0.65–1.22) | 0.29 (0.06–0.82) |
| ***Gypsophila*-Crown group** n = 149 | 9.70 (5.21–14.68) | 0.45 | 0.42 | 0.28 | 0.83 (0.62–1.27) | 0.31 (0.03–0.94) |
| ***Dianthus*-Stem group** (*Dianthus*+*Petrorhagia*+*Graecobolanthus*+*Bolanthus*) n = 350 | 7.70 (4.26–11.24) | 0.76 | 0.67 | 0.46 | 1.38 (1.04 –1.92) | 0.42 (0.05–1.07) |
| ***Dianthus*-Crown + *Velezia*** n = 300 | 4.40 (2.21–6.80) | 1.14 | 1.08 | 0.77 | 2.4 (1.75–3.36) | 0.68 (0.06–1.92) |

| | | | | | | |
|---|---|---|---|---|---|---|
| ***Saponaria***<br>n = 30 | 8.06 (3.92–12.55) | 0.34 | 0.30 | 0.16 | 0.56<br>(0.37–1.04) | 0.2<br>(0.004–0.79) |
| ***Acanthophyllum***<br>n = 100 | 6.70 (3.38–10.50) | 0.58 | 0.54 | 0.34 | 0.61<br>(0.39–1.25) | 0.24<br>(0.00–0.94) |

**Table S4** Model comparison for the BiSSE analysis of habitat: non-montane vs. montane. (sf: sampling fraction).

| **Habitat: 0 = non-montane (sf: 6/~36); 1 = montane (sf: 49/~113)** | | | | | | | | | | | | |
|---|---|---|---|---|---|---|---|---|---|---|---|---|
| Models | NP | **lnLik** | **AICc** | **ΔAIC** | **ChiSq** | **p value** | **$\lambda_0$** | **$\lambda_1$** | **$\mu_0$** | **$\mu_1$** | **$q_{01}$** | **$q_{10}$** |
| 1: ($\lambda_0 \neq \lambda_1$; $\mu_0 \neq \mu_1$; $q_{01} \neq q_{10}$)<br>Full model | 6 | -77.63 | 167.26 | 1.71 | 11.93 | 0.008 | 0.52 | 2.3 | 2e-9 | 1.01 | 0.19 | 0.16 |
| 2: ($\lambda_0 = \lambda_1$; $\mu_0 \neq \mu_1$; $q_{01} \neq q_{10}$) | 5 | -80.88 | 171.75 | 7.46 | 4.18 | 0.12 | 1.78 | 1.78 | 1.47 | 0.26 | 0.07 | 0.17 |
| 3: ($\lambda_0 \neq \lambda_1$; $\mu_0 = \mu_1$; $q_{01} \neq q_{10}$) | 5 | -78.13 | 166.27 | 1.84 | 9.81 | 0.007 | 0.5 | 1.75 | 7e-7 | 7e-7 | 0.13 | 0.2 |
| 4: ($\lambda_0 \neq \lambda_1$; $\mu_0 \neq \mu_1$; $q_{01} = q_{10}$) | 5 | -77.65 | 165.29 | 1.26 | 10.39 | 0.005 | 0.51 | 2.24 | 6e-7 | 0.89 | 0.17 | 0.17 |
| 5: ($\lambda_0 = \lambda_1$; $\mu_0 = \mu_1$; $q_{01} \neq q_{10}$) | 4 | -82.57 | 173.14 | 9.38 | 0.27 | 0.60 | 2.56 | 2.56 | 2.01 | 2.01 | 0.17 | 0.16 |
| **6: ($\lambda_0 \neq \lambda_1$; $\mu_0 = \mu_1$; $q_{01} = q_{10}$)** | **4** | **-78.29** | **164.57** | **0** | **9.65** | **0.002** | **0.5** | **1.73** | **1e-7** | **1e-7** | **0.15** | **0.15** |
| 7: ($\lambda_0 = \lambda_1$; $\mu_0 \neq \mu_1$; $q_{01} = q_{10}$) | 4 | -81.22 | 170.43 | 6.37 | 3.27 | 0.07 | 1.99 | 1.99 | 1.67 | 0.76 | 0.1 | 0.1 |
| 8: ($\lambda_0 = \lambda_1$; $\mu_0 = \mu_1$; $q_{01} = q_{10}$)<br>Null model (minimal) | 3 | -78.13 | 166.27 | 7.65 | - | - | 2.56 | 2.56 | 2.01 | 2.01 | 0.16 | 0.16 |

**Table S5** Model comparison for the BiSSE analysis of life strategy: annual vs. perennial. (sf: sampling fraction).

| Life strategy: 0 = annual (sf: 6/~17); 1 = perennial (sf: 49/~132) | | | | | | | | | | | | |
|---|---|---|---|---|---|---|---|---|---|---|---|---|
| Models | NP | **lnLik** | **AICc** | **ΔAIC** | **ChiSq** | **p value** | $\lambda_0$ | $\lambda_1$ | $\mu_0$ | $\mu_1$ | $q_{01}$ | $q_{10}$ |
| 1: ($\lambda_0 \neq \lambda_1$; $\mu_0 \neq \mu_1$; $q_{01} \neq q_{10}$) Full model | 6 | -78.56 | 169.13 | 1.39 | 10.47 | 0.01 | 0.46 | 2.58 | 8e-7 | 1.40 | 0.27 | 0.07 |
| 2: ($\lambda_0 = \lambda_1$; $\mu_0 \neq \mu_1$; $q_{01} \neq q_{10}$) | 5 | -82.50 | 175.00 | 7.26 | 2.60 | 0.27 | 2.02 | 2.02 | 1.84 | 0.68 | 0.09 | 0.08 |
| 3: ($\lambda_0 \neq \lambda_1$; $\mu_0 = \mu_1$; $q_{01} \neq q_{10}$) | 5 | -79.61 | 169.23 | 1.49 | 8.38 | 0.01 | 0.39 | 1.73 | 4e-8 | 4e-8 | 0.16 | 0.10 |
| 4: ($\lambda_0 \neq \lambda_1$; $\mu_0 \neq \mu_1$; $q_{01} = q_{10}$) | 5 | -79.67 | 169.34 | 1.6 | 8.26 | 0.02 | 0.40 | 2.01 | 4e-8 | 0.43 | 0.13 | 0.13 |
| 5: ($\lambda_0 = \lambda_1$; $\mu_0 = \mu_1$; $q_{01} \neq q_{10}$) | 4 | -83.66 | 175.31 | 7.57 | 0.29 | 0.59 | 2.80 | 2.80 | 2.30 | 2.30 | 0.33 | 0.10 |
| **6: ($\lambda_0 \neq \lambda_1$; $\mu_0 = \mu_1$; $q_{01} = q_{10}$)** | **4** | **-79.87** | **167.74** | **0** | **7.86** | **5e-3** | **0.39** | **1.75** | **1e-5** | **1e-5** | **0.13** | **0.13** |
| 7: ($\lambda_0 = \lambda_1$; $\mu_0 \neq \mu_1$; $q_{01} = q_{10}$) | 4 | -82.51 | 173.01 | 5.27 | 2.59 | 0.11 | 1.99 | 1.99 | 1.82 | 0.62 | 0.08 | 0.08 |
| 8: ($\lambda_0 = \lambda_1$; $\mu_0 = \mu_1$; $q_{01} = q_{10}$) Null model (minimal) | 3 | -83.80 | 173.60 | 5.86 | - | - | 2.79 | 2.79 | 2.29 | 2.29 | 0.09 | 0.09 |

**Table S6** Model comparison for the BiSSE analysis of life form: herbaceous vs. woody. (sf: sampling fraction).

| Life form: 0 = herbaceous (sf: 26/~69); 1 = woody (sf: 29/~80) | | | | | | | | | | | | |
|---|---|---|---|---|---|---|---|---|---|---|---|---|
| Models | NP | **lnLik** | **AICc** | **ΔAIC** | **ChiSq** | **p value** | **$\lambda_0$** | **$\lambda_1$** | **$\mu_0$** | **$\mu_1$** | **$q_{01}$** | **$q_{10}$** |
| 1e: ($\lambda_0 \neq \lambda_1$; $\mu_0 \neq \mu_1$; $q_{01} \neq q_{10}$) Full model | 6 | -98.59 | 209.18 | 1.27 | 12.53 | 0.01 | 1.9e-6 | 2.87 | 2.44 | 0.19 | 2.4e-6 | 2.4 |
| 2: ($\lambda_0 = \lambda_1$; $\mu_0 \neq \mu_1$; $q_{01} \neq q_{10}$) | 5 | -104.67 | 219.35 | 11.44 | 0.36 | 0.83 | 2.79 | 2.79 | 1.39 | 3.05 | 2.5e+6 | 2.1e+6 |
| **3: ($\lambda_0 \neq \lambda_1$; $\mu_0 = \mu_1$; $q_{01} \neq q_{10}$)** | **5** | **-98.96** | **207.91** | **0** | **11.8** | **0.003** | **2e-8** | **3.59** | **1.71** | **1.71** | **0.12** | **1.72** |
| 4: ($\lambda_0 \neq \lambda_1$; $\mu_0 \neq \mu_1$; $q_{01} = q_{10}$) | 5 | -102.13 | 214.26 | 6.35 | 5.45 | 0.07 | 3e-8 | 4.72 | 0.66 | 3.77 | 1.28 | 1.28 |
| 5: ($\lambda_0 = \lambda_1$; $\mu_0 = \mu_1$; $q_{01} \neq q_{10}$) | 4 | -104.67 | 217.35 | 9.44 | 0.36 | 0.55 | 2.79 | 2.79 | 2.29 | 2.29 | 4.5e+5 | 3.8e+5 |
| 6: ($\lambda_0 \neq \lambda_1$; $\mu_0 = \mu_1$; $q_{01} = q_{10}$) | 4 | -103.46 | 214.92 | 7.01 | 2.79 | 0.09 | 5.1e-7 | 3.49 | 1.87 | 1.87 | 2.37 | 2.37 |
| 7: ($\lambda_0 = \lambda_1$; $\mu_0 \neq \mu_1$; $q_{01} = q_{10}$) | 4 | -104.66 | 217.33 | 9.42 | 0.38 | 0.54 | 2.77 | 2.77 | 4.29 | 0.59 | 10.33 | 10.33 |
| 8: ($\lambda_0 = \lambda_1$; $\mu_0 = \mu_1$; $q_{01} = q_{10}$) Null model (minimal) | 3 | -104.85 | 215.71 | 7.80 | - | - | 2.78 | 2.78 | 2.28 | 2.28 | 19.04 | 19.04 |

**Table S7** Model comparison for BiSSE analysis of calyx shape: tubiform vs. campanulate/turbinate. (sf: sampling fraction).

| Calyx shape: 0 = tubiform (sf: 6/~16); 1 = campanulate/turbinate (sf: 49/133) | | | | | | | | | | | | |
|---|---|---|---|---|---|---|---|---|---|---|---|---|
| Models | NP | **lnLik** | **AICc** | **ΔAIC** | **ChiSq** | **p value** | **$\lambda_0$** | **$\lambda_1$** | **$\mu_0$** | **$\mu_1$** | **$q_{01}$** | **$q_{10}$** |
| 1: ($\lambda_0 \neq \lambda_1$; $\mu_0 \neq \mu_1$; $q_{01} \neq q_{10}$) Full model | 6 | -82.7 | 177.4 | 0.52 | 9.24 | 0.03 | 1e-8 | 3 | 0.37 | 2.37 | 0.35 | 0.16 |
| 2: ($\lambda_0 = \lambda_1$; $\mu_0 \neq \mu_1$; $q_{01} \neq q_{10}$) | 5 | -84.22 | 178.44 | 1.56 | 6.2 | 0.05 | 2.67 | 2.67 | 5.36 | 1.73 | 6e-7 | 0.47 |
| 3: ($\lambda_0 \neq \lambda_1$; $\mu_0 = \mu_1$; $q_{01} \neq q_{10}$) | 5 | -84.62 | 179.23 | 2.35 | 5.41 | 0.07 | 0.36 | 1.68 | 4e-8 | 4e-8 | 0.24 | 0.15 |
| 4: ($\lambda_0 \neq \lambda_1$; $\mu_0 \neq \mu_1$; $q_{01} = q_{10}$) | 5 | -84.73 | 179.47 | 2.59 | 5.17 | 0.08 | 0.37 | 1.95 | 1.5e-7 | 0.44 | 0.19 | 0.19 |
| 5: ($\lambda_0 = \lambda_1$; $\mu_0 = \mu_1$; $q_{01} \neq q_{10}$) | 4 | -85.34 | 178.68 | 1.8 | 3.95 | 0.05 | 2.79 | 2.79 | 2.29 | 2.29 | 3.05 | 0.41 |
| 6: ($\lambda_0 \neq \lambda_1$; $\mu_0 = \mu_1$; $q_{01} = q_{10}$) | 4 | -84.92 | 177.84 | 0.96 | 4.8 | 0.03 | 0.36 | 1.71 | 1.2e-7 | 1e-7 | 0.18 | 0.18 |
| **7: ($\lambda_0 = \lambda_1$; $\mu_0 \neq \mu_1$; $q_{01} = q_{10}$)** | **4** | **-84.44** | **176.88** | **0** | **5.76** | **0.02** | **2.7** | **2.7** | **4.78** | **1.86** | **0.42** | **0.42** |
| 8: ($\lambda_0 = \lambda_1$; $\mu_0 = \mu_1$; $q_{01} = q_{10}$) Null model (minimal) | 3 | -87.32 | 180.64 | 3.76 | - | - | 2.79 | 2.79 | 2.29 | 2.29 | 0.1 | 0.1 |

**Table S8** Model comparison for the BiSSE analysis of elevation: low vs. high. (sf: sampling fraction).

| Elevation: 0 = low (sf: 19/~56); 1 = high (sf: 36/~93) | | | | | | | | | | | | |
|---|---|---|---|---|---|---|---|---|---|---|---|---|
| Models | NP | **lnLik** | **AICc** | **ΔAIC** | **ChiSq** | **p value** | **$\lambda_0$** | **$\lambda_1$** | **$\mu_0$** | **$\mu_1$** | **$q_{01}$** | **$q_{10}$** |
| 1: ($\lambda_0 \neq \lambda_1$; $\mu_0 \neq \mu_1$; $q_{01} \neq q_{10}$) Full model | 6 | -96.01 | 204.02 | 1.77 | 12.6 | 0.01 | 8.6e-7 | 2.8 | 2.12 | 0.91 | 5.7e-7 | 1.55 |
| 2: ($\lambda_0 = \lambda_1$; $\mu_0 \neq \mu_1$; $q_{01} \neq q_{10}$) | 5 | -98.41 | 206.83 | 4.58 | 7.79 | 0.02 | 2.38 | 2.38 | 4.82 | 8e-6 | 2.2e-7 | 2.02 |
| **3: ($\lambda_0 \neq \lambda_1$; $\mu_0 = \mu_1$; $q_{01} \neq q_{10}$)** | **5** | **-96.13** | **202.25** | **0** | **12.37** | **0.002** | **4.7e-9** | **3.12** | **1.59** | **1.59** | **0.1** | **1.26** |
| 4: ($\lambda_0 \neq \lambda_1$; $\mu_0 \neq \mu_1$; $q_{01} = q_{10}$) | 5 | -97.7 | 205.4 | 3.15 | 9.22 | 0.01 | 3.8e-7 | 3.83 | 0.65 | 2.89 | 0.96 | 0.96 |
| 5: ($\lambda_0 = \lambda_1$; $\mu_0 = \mu_1$; $q_{01} \neq q_{10}$) | 4 | -101.68 | 211.37 | 9.12 | 1.25 | 0.26 | 2.77 | 2.77 | 2.27 | 2.27 | 1.99 | 1.33 |
| 6: ($\lambda_0 \neq \lambda_1$; $\mu_0 = \mu_1$; $q_{01} = q_{10}$) | 4 | -98.71 | 205.42 | 3.17 | 7.19 | 0.01 | 2.4e-7 | 3.06 | 1.68 | 1.68 | 1.57 | 1.57 |
| 7: ($\lambda_0 = \lambda_1$; $\mu_0 \neq \mu_1$; $q_{01} = q_{10}$) | 4 | -100.99 | 209.98 | 7.73 | 2.63 | 0.1 | 2.71 | 2.71 | 3.29 | 1.57 | 1.56 | 1.56 |
| 8: ($\lambda_0 = \lambda_1$; $\mu_0 = \mu_1$; $q_{01} = q_{10}$) Null model (minimal) | 3 | -102.31 | 210.62 | 8.37 | - | - | 2.77 | 2.77 | 2.27 | 2.27 | 0.96 | 0.96 |

**Table S9** Model comparison for GeoSSE analyses of the geographical distribution using *rps16* dataset. (sf: sampling fraction).

Distribution: 0 = both regions/widespread (sf: 13/~23); 1 = Endemic to Irano-Anatolian and Caucasus biodiversity hotspots (sf: 33/~84); 2 = Endemic to other regions (sf: 9/~42)

| Models | NP | **lnLik** | **AICc** | **ΔAIC** | **ChiSq** | **p value** | $\lambda_A$ | $\lambda_B$ | $\lambda_{AB}$ | $\mu_A$ | $\mu_B$ | $d_A$ | $d_B$ |
|---|---|---|---|---|---|---|---|---|---|---|---|---|---|
| 1: ($\lambda_0 \neq \lambda_1$; $\mu_0 \neq \mu_1$; $q_{01} \neq q_{10}$) Full model | 7 | -112.92 | 239.84 | 3.83 | 3.36 | 0.34 | 2.12 | 1.83 | 8e-9 | 1.71 | 2.36 | 0.84 | 0.46 |
| 2: ($\lambda_A$=$\lambda_B$) | 6 | -112.93 | 237.86 | 1.85 | 3.33 | 0.19 | 2.08 | 2.08 | 1e-8 | 1.66 | 2.73 | 0.93 | 0.42 |
| 3: ($\mu_A$=$\mu_B$) | 6 | -112.95 | 237.91 | 1.90 | 3.28 | 0.19 | 2.19 | 1.48 | 5e-9 | 1.78 | 1.78 | 0.7 | 0.53 |
| 4: ($d_A$=$d_B$) | 6 | -113 | 238 | 1.99 | 3.19 | 0.2 | 2.22 | 1.48 | 5e-8 | 1.85 | 1.76 | 0.65 | 0.65 |
| 5: ($\lambda_A$=$\lambda_B$; $\mu_A$=$\mu_B$) | 5 | -114.01 | 238.02 | 2.01 | 1.17 | 0.28 | 2.17 | 2.17 | 2e-9 | 2.21 | 2.21 | 0.53 | 1.3 |
| 6: ($\lambda_A$=$\lambda_B$; $d_A$=$d_B$) | 5 | -113.16 | 236.32 | 0.31 | 2.88 | 0.09 | 2.11 | 2.11 | 2e-7 | 1.82 | 2.57 | 0.76 | 0.76 |
| **7: ($\mu_A$=$\mu_B$; $d_A$=$d_B$)** | **5** | **-113** | **236.01** | **0** | **3.19** | **0.07** | **2.21** | **1.54** | **3e-8** | **1.85** | **1.85** | **0.66** | **0.66** |
| 8: ($\lambda_A$=$\lambda_B$; $\mu_A$=$\mu_B$; $d_A$=$d_B$) Null model (minimal) | 4 | -114.6 | 237.19 | 1.18 | - | - | 2.04 | 2.04 | 1e-7 | 1.88 | 1.88 | 0.62 | 0.62 |
| 9: ($\lambda_A$= 0) | 6 | -117.78 | 247.56 | 11.55 | -6.37 | 1 | 0 | 1e-8 | 21.37 | 2.84 | 5.12 | 4.76 | 8.04 |
| 10: ($\lambda_B$= 0) | 6 | -117.78 | 247.56 | 11.55 | -6.37 | 1 | 2e-8 | 0 | 21.36 | 2.84 | 5.12 | 4.75 | 8.04 |
| 11: ($\lambda_{AB}$= 0) | 6 | -112.92 | 237.84 | 1.83 | 3.36 | 0.19 | 2.12 | 1.83 | 0 | 1.71 | 2.36 | 0.84 | 0.46 |
| 12: ($\mu_A$= 0) | 6 | -114.7 | 241.39 | 5.38 | -0.2 | 1 | 1.5 | 0.69 | 0.23 | 0 | 0.17 | 0.38 | 0.2 |
| 13: ($\mu_B$= 0) | 6 | -114.03 | 240.06 | 4.05 | 1.13 | 0.57 | 2.03 | 0.52 | 6e-8 | 1.1 | 0 | 0.33 | 0.45 |
| 14: ($\mu_A$= 0; $\mu_B$= 0) | 5 | -114.78 | 239.56 | 3.55 | -0.36 | 1 | 1.51 | 0.58 | 0.36 | 0 | 0 | 0.36 | 0.22 |

| **Models** | logL | AICc | ΔAIC | LRT | LRT p-value | λ values or as a function of time $\lambda(t) = \lambda 0 + \alpha t$ (linear) $\lambda(t) = \lambda 0 \times e^{\alpha t}$ (exponential) | μ values or as a function of time $\mu(t) = \mu 0 + \alpha t$ (linear) $\mu(t) = \mu 0 \times e^{\alpha t}$ (exponential) |
|---|---|---|---|---|---|---|---|
| **Caryophylleae (101/630):** | | | | | | | |
| (1) λ constant (no μ) (Yule) | -256.70 | 515.44 | 150.55 | - | - | $\lambda = 0.66$ | - |
| **(2) λ and μ constant** | **-180.38** | **364.89** | **0** | **152.63** | **5e-35** | $\boldsymbol{\lambda = 5.46}$ | $\boldsymbol{\mu = 5.39}$ |
| (3) λ linear and no μ | -228.48 | 461.09 | 96.2 | 56.44 | 6e-14 | $\lambda(t) = \lvert 0.86 - 0.035t \rvert$ | - |
| (4) λ exponential and no μ | -193.52 | 391.16 | 26.27 | 126.37 | 2e-29 | $\lambda(t) = 1.71e^{-0.32t}$ | - |
| (5) λ Linear and μ constant | -179.85 | 365.951 | 1.06 | 153.70 | 4e-34 | $\lambda(t) = \lvert 5.11 - 0.01t \rvert$ | 4.94 |
| (6) λ exponential and μ constant | -179.85 | 365.945 | 1.06 | 153.70 | 4e-34 | $\lambda(t) = 5.104e^{-0.002t}$ | 4.93 |
| (7) λ and μ linear | -228.48 | 465.38 | 100.49 | 56.44 | 3e-12 | $\lambda(t) = \lvert 0.86 - 0.035t \rvert$ | $\mu(t) = \lvert -0.44 - 0.08t \rvert$ |
| (8) λ exponential and μ linear | -180.60 | 369.62 | 4.73 | 152.20 | 9e-33 | $\lambda(t) = 3.93e^{-0.10t}$ | $\mu(t) = \lvert -3.21 + 0.18t \rvert$ |
| (9) λ and μ exponential | -186.05 | 380.52 | 15.63 | 141.29 | 2e-30 | $\lambda(t) = 3.05e^{0.32t}$ | $\mu(t) = 3.06e^{0.32t}$ |
| (10) λ linear and μ exponential | -229.63 | 467.67 | 102.78 | 56.44 | 3e-12 | $\lambda(t) = \lvert 0.866 - 0.035t \rvert$ | $\mu(t) = (2.05e - 7)e^{0.44t}$ |
| (11) λ constant and μ linear | -256.68 | 519.61 | 154.72 | 0.04 | 0.998 | $\lambda = 0.66$ | $\mu(t) = \lvert -0.24 - 0.33t \rvert$ |
| (12) λ constant and μ exponential | -179.85 | 365.96 | 1.07 | 153.69 | 4e-33 | $\lambda = 5.12$ | $\mu(t) = 4.95e^{0.002t}$ |
| ***Gypsophila* (25/152):** | | | | | | | |
| (1) λ constant (no μ) (Yule) | -41.77 | 85.71 | 13.97 | - | - | $\lambda = 1.20$ | - |
| (2) λ and μ constant | -34.02 | 72.59 | 0.85 | 15.50 | 8e-05 | $\lambda = 5.20$ | $\mu = 4.85$ |
| (3) λ linear and no μ | -35.72 | 75.98 | 4.24 | 12.10 | 5e-04 | $\lambda(t) = \lvert 1.81 - 0.34t \rvert$ | - |
| **(4) λ exponential and no μ** | **-33.60** | **71.74** | **0** | **16.35** | **5e-05** | $\boldsymbol{\lambda(t) = 2.72e^{-0.63t}}$ | **-** |
| (5) λ Linear and μ constant | -33.67 | 74.48 | 2.74 | 16.21 | 3e-04 | $\lambda(t) = \lvert 4.22 - 0.25t \rvert$ | $\mu = 3.30$ |

| | | | | | | | |
|---|---|---|---|---|---|---|---|
| (6) λ exponential and μ constant | -33.63 | 74.41 | 2.67 | 16.27 | 3e-04 | $\lambda(t) = 4.05e^{-0.084t}$ | $\mu = 2.99$ |
| (7) λ and μ linear | -35.72 | 81.44 | 9.7 | 12.10 | 0.007 | $\lambda(t) = \lvert 1.81 - 0.34t \rvert$ | $\mu(t) = \lvert -0.0005 - 1.38t \rvert$ |
| (8) λ exponential and μ linear | -33.60 | 77.19 | 5.45 | 16.35 | 0.001 | $\lambda(t) = 2.71e^{-0.625t}$ | $\mu(t) = \lvert -0.16 - 1.78t \rvert$ |
| (9) λ and μ exponential | -33.47 | 76.94 | 5.2 | 16.59 | 9e-04 | $\lambda(t) = 3.56e^{-0.46t}$ | $\mu(t) = 1.66e^{-0.41t}$ |
| (10) λ linear and μ exponential | -33.67 | 77.33 | 5.59 | 16.21 | 0.001 | $\lambda(t) = \lvert 4.22 - 0.24t \rvert$ | $\mu(t) = 3.30e^{0.003t}$ |
| (11) λ constant and μ linear | -41.77 | 90.68 | 18.94 | 0.004 | 1 | $\lambda = 1.20$ | $\mu(t) = \lvert -0.74 - 0.42t \rvert$ |
| (12) λ constant and μ exponential | -33.69 | 74.52 | 2.78 | 16.16 | 0.001 | $\lambda = 4.32$ | $\mu(t) = 3.51e^{0.06t}$ |
| ***Dianthus* (38/300):** | | | | | | | |
| (1) λ constant (no μ) (Yule) | -41.72 | 85.55 | 21.56 | - | - | $\lambda = 2.50$ | - |
| (2) λ and μ constant | -30.08 | 64.50 | 0.51 | 23.28 | 1e-06 | $\lambda = 11.38$ | $\mu = 10.50$ |
| (3) λ linear and no μ | -33.19 | 70.73 | 6.74 | 17.05 | 4e-05 | $\lambda(t) = \lvert 3.56 - 1.13t \rvert$ | - |
| **(4) λ exponential and no μ** | **-29.82** | **63.99** | **0** | **23.79** | **1e-06** | $\boldsymbol{\lambda(t) = 5.42e^{-1.11t}}$ | - |
| (5) λ linear and μ constant | -29.47 | 65.66 | 1.67 | 24.49 | 5e-06 | $\lambda(t) = \lvert 9.12 - 0.85t \rvert$ | $\mu = 7.20$ |
| (6) λ exponential and μ constant | -29.45 | 65.61 | 1.62 | 24.53 | 5e-06 | $\boldsymbol{\lambda(t) = 9.09e^{-0.11t}}$ | $\mu = 7.09$ |
| (7) λ and μ linear | -33.19 | 75.60 | 11.61 | 17.05 | 7e-04 | $\lambda(t) = \lvert 3.56 - 1.13t \rvert$ | $\mu(t) = \lvert -0.22 - 1.42t \rvert$ |
| (8) λ exponential and μ linear | -29.82 | 68.85 | 4.86 | 23.79 | 3e-05 | $\boldsymbol{\lambda(t) = 5.40e^{-1.11t}}$ | $\mu(t) = \lvert -3.81 + 0.13t \rvert$ |
| (9) λ and μ exponential | -29.34 | 67.89 | 3.9 | 24.76 | 2e-05 | $\boldsymbol{\lambda(t) = 8.33e^{-0.68t}}$ | $\mu(t) = 5.20e^{-0.63t}$ |
| (10) λ linear and μ exponential | -32.94 | 75.09 | 11.1 | 17.56 | 5e-04 | $\lambda(t) = \lvert 3.93 - 1.89t \rvert$ | $\mu(t) = 0.001e^{-3.39t}$ |
| (11) λ constant and μ linear | -41.70 | 90.10 | 26.11 | 0.04 | 0.998 | $\lambda = 2.50$ | $\mu(t) = \lvert -1.25 - 1.14t \rvert$ |
| (12) λ constant and μ exponential | -29.49 | 65.68 | 1.69 | 16.16 | 0.001 | $\lambda = 9.36$ | $\mu(t) = 7.63e^{0.09t}$ |

**Table S10** Comparison of different time-dependent diversification models applied on Caryophylleae, *Gypsophila*, and *Dianthus* clades using RPANDA package in R for the *matK* dataset. The best model for each clade is chosen via the corrected Akaike Information Criterion (AICc) and Likelihood Ratio Test (LRT), significant at $P < 0.05$, which is in bold. λ = speciation rate; μ = extinction rate (in events/Ma/lineage

**Table S11** Comparison of different time-dependent diversification models applied on major clades inside Caryophylleae using the RPANDA package in R for the *rps16* dataset. The best model for each clade is chosen via the corrected Akaike Information Criterion (AICc) and Likelihood Ratio Test (LRT), significant at P < 0.05, which is in bold. λ = the speciation rate; μ = the extinction rate (in events/Ma/lineage).

| **Models** | logL | AICc | ΔAIC | LRT | LRT p-value | λ value or as a function of time $\lambda(t) = \lambda 0 + \alpha t$ (linear) $\lambda(t) = \lambda 0 \times e^{\alpha t}$ (exponential) | μ value or as a function of time $\mu(t) = \mu 0 + \beta t$ (linear) $\mu(t) = \mu 0 \times e^{\beta t}$ (exponential) |
|---|---|---|---|---|---|---|---|
| ***Gypsophila* (55/152):** | | | | | | | |
| (1) λ constant (no μ) (Yule) | -74.12 | 150.31 | 14.53 | - | - | $\lambda = 1.16$ | - |
| (2) λ and μ constant | -66.63 | 137.49 | 1.71 | 14.97 | 1e-04 | $\lambda = 2.85$ | $\mu = 2.35$ |
| (3) λ linear and no μ | -67.33 | 138.90 | 3.12 | 13.57 | 2e-04 | $\lambda(t) = \lvert 1.58 - 0.27t \rvert$ | - |
| **(4) λ exponential and no μ** | **-65.78** | **135.78** | **0** | **16.68** | **4e-05** | $\boldsymbol{\lambda(t) = 2.00e^{-0.44t}}$ | - |
| (5) λ Linear and μ constant | -66.02 | 138.51 | 2.73 | 16.19 | 3e-04 | $\lambda(t) = \lvert 2.42 - 0.21t \rvert$ | $\mu = 1.39$ |
| (6) λ exponential and μ constant | -65.79 | 138.05 | 2.27 | 16.65 | 2e-04 | $\boldsymbol{\lambda(t) = 1.98e^{-0.42t}}$ | $\mu = 0.03$ |
| (7) λ and μ linear | -67.33 | 143.47 | 7.69 | 13.57 | 0.004 | $\lambda(t) = \lvert 1.58 - 0.27t \rvert$ | $\mu(t) = \lvert -0.45 - 1.26t \rvert$ |
| (8) λ exponential and μ linear | -65.78 | 140.36 | 4.58 | 16.67 | 8e-04 | $\lambda(t) = 2.00e^{-0.44t}$ | $\mu(t) = \lvert -0.05 - 1.09t \rvert$ |
| (9) λ exponential and μ exponential | -65.75 | 140.30 | 4.52 | 16.73 | 8e-04 | $\lambda(t) = 2.16e^{-0.39t}$ | $\mu(t) = 0.41e^{-0.33t}$ |
| (10) λ linear and μ exponential | -67.93 | 144.66 | 8.88 | 12.38 | 0.006 | $\lambda(t) = \lvert 1.87 - 0.32t \rvert$ | $\mu(t) = 0.45e^{-1.38t}$ |
| (11) λ constant and μ linear | -74.12 | 154.71 | 18.93 | -0.004 | 1 | $\lambda = 1.16$ | $\mu(t) = \lvert -0.45 - 0.62t \rvert$ |
| (12) λ constant and μ exponential | -66.07 | 138.62 | 2.84 | 16.09 | 0.001 | $\lambda = 2.49$ | $\mu(t) = 1.61e^{0.1t}$ |
| ***Dianthus* (10/300):** | | | | | | | |
| (1) λ constant (no μ) (Yule) | -19.64 | 41.77 | 7.09 | - | - | $\lambda = 1.44$ | - |
| (2) λ and μ constant | -14.69 | 35.09 | 0.41 | 9.90 | 0.002 | $\lambda = 15.22$ | $\mu = 14.90$ |
| (3) λ linear and no μ | -15.60 | 36.92 | 2.24 | 8.06 | 0.004 | $\lambda(t) = \lvert 2.44 - 0.59t \rvert$ | - |
| **(4) λ exponential and no μ** | **-14.48** | **34.68** | **0** | **10.31** | **0.001** | $\boldsymbol{\lambda(t) = 4.41e^{-0.92t}}$ | **-** |

| | | | | | | | |
|---|---|---|---|---|---|---|---|
| (5) λ Linear and μ constant | -14.32 | 38.63 | 3.95 | 10.64 | 0.005 | $\lambda(t) = \lvert 9.45 - 0.62t \rvert$ | $\mu = 8.03$ |
| (6) λ exponential and μ constant | -14.32 | 38.65 | 3.97 | 10.63 | 0.005 | $\lambda(t) = 9.46e^{-0.07t}$ | $\mu = 8.00$ |
| (7) λ and μ linear | -15.38 | 46.77 | 12.09 | 8.51 | 0.037 | $\lambda(t) = \lvert 2.60 - 0.62t \rvert$ | $\mu(t) = \lvert -1.5 + 1.05t \rvert$ |
| (8) λ exponential and μ linear | -14.48 | 44.96 | 10.28 | 10.31 | 0.016 | $\lambda(t) = 4.41e^{-0.92t}$ | $\mu(t) = \lvert -1.9 + 0.39t \rvert$ |
| (9) λ exponential and μ exponential | -14.38 | 44.76 | 10.08 | 10.51 | 0.015 | $\lambda(t) = 6.10e^{-0.76t}$ | $\mu(t) = 2.64e^{-0.68t}$ |
| (10) λ linear and μ exponential | -14.32 | 44.63 | 9.95 | 10.64 | 0.014 | $\lambda(t) = \lvert 9.45 - 0.50t \rvert$ | $\mu(t) = 8.04e^{0.02t}$ |
| (11) λ constant and μ linear | -19.63 | 49.27 | 14.59 | 0.004 | 1 | $\lambda = 1.44$ | $\mu(t) = \lvert -0.79 - 0.56t \rvert$ |
| (12) λ constant and μ exponential | -14.31 | 38.62 | 3.94 | 10.65 | 0014 | $\lambda = 9.46$ | $\mu(t) = 8.13e^{0.07t}$ |
| ***Saponaria* (7/30):** | | | | | | | |
| **(1) λ constant (no μ) (Yule)** | **-9.79** | **22.38** | 0 | - | - | $\boldsymbol{\lambda = 0.85}$ | - |
| (2) λ and μ constant | -9.61 | 26.22 | 3.84 | 0.36 | 0.547 | $\lambda = 1.45$ | $\mu = 0.94$ |
| (3) λ linear and no μ | -9.37 | 25.74 | 3.36 | 0.84 | 0.358 | $\lambda(t) = \lvert 1.36 - 0.38t \rvert$ | - |
| (4) λ exponential and no μ | -9.49 | 25.97 | 3.59 | 0.61 | 0.435 | $\lambda(t) = 1.31e^{-0.35t}$ | - |
| (5) λ Linear and μ constant | -9.37 | 32.74 | 10.36 | 0.84 | 0.656 | $\lambda(t) = \lvert 1.36 - 0.38t \rvert$ | 1.78e-06 |
| (6) λ exponential and μ constant | -9.49 | 32.97 | 10.59 | 0.61 | 0.737 | $\lambda(t) = 1.31e^{-0.35t}$ | 2.03e-07 |
| (7) λ and μ linear | -9.11 | 46.21 | 23.83 | 1.37 | 0.713 | $\lambda(t) = \lvert 0.75 + 0.44t \rvert$ | $\mu(t) = \lvert -2.40 + 2.36t \rvert$ |
| (8) λ exponential and μ linear | -9.09 | 46.18 | 23.8 | 1.40 | 0.706 | $\lambda(t) = 0.73e^{0.51.t}$ | $\mu(t) = \lvert -3.07 + 3.05t \rvert$ |
| (9) λ exponential and μ exponential | -9.04 | 46.09 | 23.71 | 1.49 | 0.685 | $\lambda(t) = 0.86e^{0.22t}$ | $\mu(t) = 0.01e^{2.63t}$ |
| (10) λ linear and μ exponential | -9.37 | 46.74 | 24.36 | 0.84 | 0.839 | $\lambda(t) = \lvert 1.36 - 0.38t \rvert$ | $\mu(t = (2.50e - 06)e^{-0.45*t}$ |
| (11) λ constant and μ linear | -9.79 | 33.58 | 11.2 | -2e-4 | 1 | $\lambda = 0.85$ | $\mu(t) = \lvert -0.24 - 0.54t \rvert$ |
| (12) λ constant and μ exponential | -9.00 | 32.01 | 9.63 | 16.09 | 0.001 | $\lambda = 0.99$ | $\mu(t) = (e - 04)e^{4.18t}$ |
| ***Acanthophyllum* (10/95):** | | | | | | | |
| (1) λ constant (no μ) (Yule) | -18.47 | 39.44 | 4.17 | - | - | $\lambda = 0.97$ | - |
| (2) λ and μ constant | -15.39 | 36.50 | 1.23 | 6.15 | 0.013 | $\lambda = 5.09$ | $\mu = 4.90$ |
| (3) λ linear and no μ | -15.20 | 36.12 | 0.85 | 6.53 | 0.011 | $\lambda(t) = \lvert 1.60 - 0.32t \rvert$ | - |
| **(4) λ exponential and no μ** | **-14.78** | **35.27** | **0** | **7.38** | **0.007** | $\boldsymbol{\lambda(t) = 2.57e^{-0.68t}}$ | - |
| (5) λ Linear and μ constant | -14.60 | 39.21 | 3.94 | 7.73 | 0.021 | $\lambda(t) = \lvert 2.70 - 0.54t \rvert$ | $\mu = 1.14$ |
| (6) λ exponential and μ constant | -14.74 | 39.48 | 4.21 | 7.46 | 0.024 | $\lambda(t) = 2.98e^{-0.32t}$ | $\mu = 1.21$ |

| | | | | | | | |
|---|---|---|---|---|---|---|---|
| (7) λ and μ linear | -15.20 | 46.40 | 11.13 | 6.53 | 0.088 | $\lambda(t) = \lvert 1.60 - 0.32t \rvert$ | $\mu(t) = \lvert -1.18 - 0.20t \rvert$ |
| (8) λ exponential and μ linear | -13.97 | 43.94 | 8.67 | 9.00 | 0.029 | $\lambda(t) = 2.21e^{-0.50t}$ | $\mu(t) = \lvert -54.39 + 18.20t \rvert$ |
| (9) λ exponential and μ exponential | -14.48 | 44.96 | 9.69 | 7.98 | 0.046 | $\lambda(t) = 2.79e^{0.99t}$ | $\mu(t) = 2.81e^{1.00t}$ |
| (10) λ linear and μ exponential | -14.56 | 45.13 | 9.86 | 7.81 | 0.050 | $\lambda(t) = \lvert 2.96 - 0.66t \rvert$ | $\mu(t) = 1.59e^{-0.23t}$ |
| (11) λ constant and μ linear | -18.47 | 46.93 | 11.66 | 0.002 | 1 | $\lambda = 0.97$ | $\mu(t) = \lvert -0.30 - 0.61t \rvert$ |
| (12) λ constant and μ exponential | -14.59 | 39.19 | 3.92 | 7.75 | 0.05 | $\lambda = 2.55$ | $\mu(t) = 1.22e^{0.33t}$ |

**Table S12** Comparison of different time-dependent diversification models applied on major clades inside Caryophylleae using the RPANDA package in R for the ITS dataset. The best model for each clade is chosen via the corrected Akaike Information Criterion (AICc) and Likelihood Ratio Test (LRT), significant at P < 0.05, which is in bold. λ = the speciation rate; μ = the extinction rate (in events/Ma/lineage).

| **Models** | logL | AICc | ΔAIC | LRT | LRT p-value | λ value or as a function of time $\lambda(t) = \lambda 0 + \alpha t$ (linear) $\lambda(t) = \lambda 0 \times e^{\alpha t}$ (exponential) | μ value or as a function of time $\mu(t) = \mu 0 + \beta t$ (linear) $\mu(t) = \mu 0 \times e^{\beta t}$ (exponential) |
|---|---|---|---|---|---|---|---|
| ***Gypsophila* (71/152):** | | | | | | | |
| (1) λ constant (no μ) (Yule) | -123.99 | 250.03 | 10.42 | - | - | $\lambda = 0.67$ | - |
| **(2) λ and μ constant** | **-117.72** | **239.61** | **0** | **12.54** | **4e-04** | $\boldsymbol{\lambda = 1.42}$ | $\boldsymbol{\mu = 1.09}$ |
| (3) λ linear and no μ | -118.57 | 241.31 | 1.7 | 10.84 | 0.001 | $\lambda(t) = \|0.89 - 0.09t\|$ | - |
| (4) λ exponential and no μ | -117.82 | 239.82 | 0.21 | 12.33 | 4e-04 | $\lambda(t) = 1.01e^{-0.21t}$ | - |
| (5) λ Linear and μ constant | -117.44 | 241.23 | 1.62 | 13.10 | 0.001 | $\lambda(t) = \|1.28 - 0.05t\|$ | $\mu = 0.76$ |
| (6) λ exponential and μ constant | -117.43 | 241.22 | 1.61 | 13.11 | 0.001 | $\lambda(t) = 1.28e^{-0.05t}$ | $\mu = 0.73$ |
| (7) λ and μ linear | -118.57 | 245.74 | 6.13 | 10.84 | 0.013 | $\lambda(t) = \|0.89 - 0.09t\|$ | $\mu(t) = \|0.0003 - 0.48t\|$ |
| (8) λ exponential and μ linear | -117.21 | 243.02 | 3.41 | 13.56 | 0.004 | $\lambda(t) = 1.51e^{-0.28t}$ | $\mu(t) = \|1.79 - 1.80t\|$ |
| (9) λ exponential and μ exponential | -117.44 | 243.48 | 3.87 | 13.10 | 0.004 | $\lambda(t) = 1.27e^{-0.16t}$ | $\mu(t) = 0.61e^{-0.22t}$ |
| (10) λ linear and μ exponential | -117.43 | 243.47 | 3.86 | 13.11 | 0.004 | $\lambda(t) = \|1.29 - 0.08t\|$ | $\mu(t) = 0.76e^{-0.07t}$ |
| (11) λ constant and μ linear | -123.99 | 254.34 | 14.73 | -0.007 | 1 | $\lambda = 0.67$ | $\mu(t) = \|-0.23 - 0.31t\|$ |
| (12) λ constant and μ exponential | -117.44 | 241.23 | 1.62 | 13.10 | 0.004 | $\lambda = 1.29$ | $\mu(t) = 0.81e^{0.05t}$ |
| ***Dianthus* (40/300):** | | | | | | | |
| (1) λ constant (no μ) (Yule) | -53.21 | 108.52 | 28.39 | - | - | $\lambda = 2.04$ | - |
| (2) λ and μ constant | -39.73 | 83.79 | 3.66 | 26.95 | 2e-07 | $\lambda = 9.31$ | $\mu = 8.68$ |
| (3) λ linear and no μ | -42.61 | 89.55 | 9.42 | 21.19 | 4e-06 | $\lambda(t) = \|2.85 - 0.65t\|$ | - |

| | | | | | | | |
|---|---|---|---|---|---|---|---|
| **(4) λ exponential and no μ** | **-37.90** | **80.13** | **0** | **30.60** | **3e-08** | $\boldsymbol{\lambda(t) = 4.34e^{-0.87t}}$ | **-** |
| (5) λ Linear and μ constant | -38.41 | 83.48 | 3.35 | 29.60 | 4e-07 | $\lambda(t) = \lvert 6.84 - 0.63t \rvert$ | $\mu = 5.11$ |
| (6) λ exponential and μ constant | -37.91 | 82.49 | 2.36 | 30.59 | 2e-07 | $\lambda(t) = 4.37e^{-0.89t}$ | $\mu = 1.31e - 06$ |
| (7) λ and μ linear | -42.61 | 94.37 | 14.24 | 21.19 | 1e-04 | $\lambda(t) = \lvert 2.85 - 0.65t \rvert$ | $\mu(t) = \lvert (6e - 5) - 2.61t \rvert$ |
| (8) λ exponential and μ linear | -37.92 | 84.98 | 4.85 | 30.58 | 1e-06 | $\lambda(t) = 4.34e^{-0.87t}$ | $\mu(t) = \lvert -1.2 - 0.05t \rvert$ |
| (9) λ exponential and μ exponential | -37.81 | 84.75 | 4.62 | 30.80 | 9e-07 | $\lambda(t) = 5.59e^{-0.68t}$ | $\mu(t) = 2.26e^{-0.59t}$ |
| (10) λ linear and μ exponential | -40.60 | 90.34 | 10.21 | 25.21 | 1e-05 | $\lambda(t) = 3.20 - 0.99t$ | $\mu(t) = 0.58e^{-2.99t}$ |
| (11) λ constant and μ linear | -53.20 | 113.08 | | 0.003 | 1 | $\lambda = 2.04$ | $\mu(t) = \lvert -0.71 - 1.39t \rvert$ |
| (12) λ constant and μ exponential | -38.48 | 83.64 | | 16.09 | 0.001 | $\lambda = 7.11$ | $\mu(t) = 5.60e^{0.09t}$ |
| ***Saponaria*:** | | | | | | | |
| **(1) λ constant (no μ) (Yule)** | **-47.895** | **97.97** | **0** | **-** | **-** | $\boldsymbol{\lambda = 0.35}$ | **-** |
| (2) λ and μ constant | -47.892 | 100.36 | 2.39 | 0.006 | 0.938 | $\lambda = 0.36$ | $\mu = 0.02$ |
| (3) λ linear and no μ | -47.889 | 100.35 | 2.38 | 0.011 | 0.915 | $\lambda(t) = \lvert 0.36 - 0.005t \rvert$ | - |
| (4) λ exponential and no μ | -47.887 | 100.35 | 2.38 | 0.016 | 0.9 | $\lambda(t) = 0.36e^{-0.01t}$ | - |
| (5) λ Linear and μ constant | -47.889 | 102.98 | 5.01 | 0.011 | 0.994 | $\lambda(t) = \lvert 0.36 - 0.005t \rvert$ | $\mu = 2.05e - 6$ |
| (6) λ exponential and μ constant | -47.887 | 102.97 | 5.00 | 0.016 | 0.992 | $\lambda(t) = 0.36e^{-0.01t}$ | $\mu = 2.82e - 7$ |
| (7) λ and μ linear | -47.889 | 105.88 | 7.91 | 0.011 | 1 | $\lambda(t) = \lvert 0.36 - 0.004t \rvert$ | $\mu(t) = \lvert -0.29 - 0.02t \rvert$ |
| (8) λ exponential and μ linear | -47.89 | 105.88 | 7.91 | 0.011 | 1 | $\lambda(t) = 0.36e^{-0.01.t}$ | $\mu(t) = \lvert -0.06 - 0.20t \rvert$ |
| (9) λ exponential and μ exponential | -47.89 | 105.88 | 7.91 | 0.011 | 1 | $\lambda(t) = 0.36e^{-0.01t}$ | $\mu(t) = (2.96e - 6)e^{-0.03t}$ |
| (10) λ linear and μ exponential | -47.89 | 105.88 | 7.91 | 0.011 | 1 | $\lambda(t) = 0.36 - 0.004t$ | $\mu(t) = (5.83e - 6)e^{-0.08t}$ |
| (11) λ constant and μ linear | -47.90 | 103.00 | 5.03 | -0.005 | 1 | $\lambda = 0.35$ | $\mu(t) = \lvert -0.07 - 0.13t \rvert$ |
| (12) λ constant and μ exponential | -47.90 | 103.00 | 5.03 | -0.005 | 1 | $\lambda = 0.35$ | $\mu(t) = (6.06e - 6)e^{-0.27t}$ |
| ***Acanthophyllum* (14/95):** | | | | | | | |
| (1) λ constant (no μ) (Yule) | -28.14 | 58.61 | 0.2 | - | - | $\lambda = 0.66$ | - |

| | | | | | | | |
|---|---|---|---|---|---|---|---|
| **(2) λ and μ constant** | **-26.66** | **58.41** | **0** | **2.96** | **0.086** | $\boldsymbol{\lambda = 1.87}$ | $\boldsymbol{\mu = 1.55}$ |
| (3) λ linear and no μ | -26.85 | 58.78 | 0.37 | 2.58 | 0.108 | $\lambda(t) = \lvert 1.00 - 0.14t \rvert$ | - |
| (4) λ exponential and no μ | -26.88 | 58.85 | 0.44 | 2.52 | 0.113 | $\lambda(t) = 1.10e^{-0.24t}$ | - |
| (5) λ Linear and μ constant | -26.65 | 61.69 | 3.28 | 2.98 | 0.225 | $\lambda(t) = \lvert 1.73 - 0.04t \rvert$ | $\mu = 1.29$ |
| (6) λ exponential and μ constant | -26.65 | 61.70 | 3.29 | 2.98 | 0.225 | $\lambda(t) = 1.75e^{-0.02t}$ | $\mu = 1.33$ |
| (7) λ and μ linear | -26.75 | 65.95 | 7.54 | 2.77 | 0.429 | $\lambda(t) = \lvert 1.25 - 0.19t \rvert$ | $\mu(t) = \lvert 1.74 - 2.30t \rvert$ |
| (8) λ exponential and μ linear | -26.88 | 66.20 | 7.79 | 2.52 | 0.472 | $\lambda(t) = 1.10e^{-0.24t}$ | $\mu(t) = \lvert -0.03 - 0.71t \rvert$ |
| (9) λ exponential and μ exponential | -26.48 | 65.40 | 6.99 | 3.32 | 0.345 | $\lambda(t) = 1.39e^{0.34t}$ | $\mu(t) = 1.21e^{0.37t}$ |
| (10) λ linear and μ exponential | -26.03 | 64.50 | 6.09 | 4.23 | 0.238 | $\lambda(t) = \lvert 2.44 - 0.38t \rvert$ | $\mu(t) = 3.42e^{-0.73t}$ |
| (11) λ constant and μ linear | -28.14 | 64.68 | 6.27 | -5e-4 | 1 | $\lambda = 0.66$ | $\mu(t) = \lvert -0.22 - 0.30t \rvert$ |
| (12) λ constant and μ exponential | -26.64 | 61.69 | 3.28 | 2.99 | 0.393 | $\lambda = 1.69$ | $\mu(t) = 1.25e^{0.03t}$ |

**Table S13** Comparison of different environmental-dependent diversification models applied on Caryophylleae, *Gypsophila*, and *Dianthus* clades using the RPANDA package in R for the *matK* dataset. The best model for each clade is chosen via the corrected Akaike Information Criterion (AICc) and Likelihood Ratio Test (LRT), significant at P < 0.05, which is in bold. λ = the speciation rate; μ = the extinction rate (in events/Ma/lineage).

| **Models** | logL | AICc | ΔAIC | LRT | LRT p-value | λ value as a function of temperature $\lambda(t) = \lambda_0 + \alpha t$ (linear) $\lambda(t) = \lambda_0 \times e^{\alpha t}$ (exponential) | μ values as a function of temperature $\mu(t) = \mu_0 + \alpha t$ (linear) $\mu(t) = \mu_0 \times e^{\alpha t}$ (exponential) |
|---|---|---|---|---|---|---|---|
| **Caryophylleae (101/630):** | | | | | | | |
| (1) λ constant (no μ) (Yule) | -256.70 | 515.44 | 159.75 | - | - | $\lambda = 0.66$ | - |
| (2) λ and μ constant | -180.38 | 364.89 | 9.2 | 152.63 | 4.6e-35 | $\lambda = 5.46$ | $\mu = 5.39$ |
| (3) λ linear and no μ | -204.35 | 412.83 | 57.14 | 104.69 | 1.4e-24 | $\lambda(t) = \lvert 1.45 - 0.19t \rvert$ | - |
| (4) λ exponential and no μ | -190.66 | 385.44 | 29.75 | 132.08 | 1.4e-30 | $\lambda(t) = 3.47e^{-0.54t}$ | - |
| (5) λ Linear and μ constant | -179.71 | 365.67 | 9.98 | 153.98 | 3.7e-34 | $\boldsymbol{\lambda(t) = \lvert 4.99 - 0.04t \rvert}$ | $\mu = 4.72$ |
| (6) λ exponential and μ constant | -179.71 | 365.66 | 9.97 | 153.99 | 3.6e-34 | $\lambda(t) = 4.99e^{-0.008t}$ | $\mu = 4.71$ |
| **(7) λ and μ linear** | **-173.63** | **355.69** | **0** | **166.13** | **8.7e-36** | $\boldsymbol{\lambda(t) = \lvert -12.25 + 13.50t \rvert}$ | $\boldsymbol{\mu(t) = \lvert 11.62 - 13.41t \rvert}$ |
| (8) λ exponential and μ linear | -180.53 | 369.48 | 13.79 | 152.34 | 8.2e-33 | $\lambda(t) = 3.05e^{0.1t}$ | $\mu(t) = \lvert -2.28 - 0.59t \rvert$ |
| (9) λ exponential and μ exponential | -181.29 | 371.00 | 15.31 | 150.82 | 1.7e-32 | $\lambda(t) = 6.48e^{-0.19t}$ | $\mu(t) = 5.93e^{-0.18t}$ |
| (10) λ linear and μ exponential | -200.79 | 410.00 | 54.31 | 111.82 | 4.4e-24 | $\lambda(t) = \lvert 1.57 - 0.21t \rvert$ | $\mu(t) = 0.66e^{-0.33t}$ |
| (11) λ constant and μ linear | -179.66 | 365.56 | 9.87 | 154.09 | 3.4e-33 | $\lambda = 4.92$ | $\mu(t) = \lvert -4.65 - 0.04t \rvert$ |
| (12) λ constant and μ exponential | -179.66 | 365.57 | 9.88 | 154.08 | 3.5e-33 | $\lambda = 4.93$ | $\mu(t) = 4.7e^{0.008t}$ |
| ***Gypsophila* (25/152):** | | | | | | | |
| (1) λ constant (no μ) (Yule) | -41.77 | 85.71 | 13.12 | - | - | $\lambda = 1.20$ | - |

| **Models** | logL | AICc | ΔAIC | LRT | LRT p-value | λ value as a function of temperature $\lambda(t) = \lambda_0 + \alpha t$ (linear) $\lambda(t) = \lambda_0 \times e^{\alpha t}$ (exponential) | μ values as a function of temperature $\mu(t) = \mu_0 + \alpha t$ (linear) $\mu(t) = \mu_0 \times e^{\alpha t}$ (exponential) |
|---|---|---|---|---|---|---|---|
| **(2) λ and μ constant** | **-34.02** | **72.59** | **0** | **15.50** | **8.3e-05** | $\boldsymbol{\lambda = 5.20}$ | $\boldsymbol{\mu = 4.85}$ |
| (3) λ linear and no μ | -36.09 | 76.72 | 4.13 | 11.37 | 7.4e-04 | $\lambda(t) = \lvert 2.85 - 0.78t \rvert$ | - |
| (4) λ exponential and no μ | -34.74 | 74.04 | 1.45 | 14.05 | 1.7e-04 | $\lambda(t) = 4.84e^{-0.65t}$ | - |
| (5) λ Linear and μ constant | -33.80 | 74.74 | 2.15 | 15.94 | 3.5e-04 | $\lambda(t) = \lvert 4.73 - 0.20t \rvert$ | $\mu = 3.78$ |
| (6) λ exponential and μ constant | -33.79 | 74.72 | 2.13 | 15.96 | 3.4e-04 | $\lambda(t) = 4.71e^{-0.05t}$ | $\mu = 3.67$ |
| (7) λ and μ linear | -33.66 | 77.33 | 4.74 | 16.21 | 0.001 | $\lambda(t) = \lvert 2.30 + 1.93t \rvert$ | $\mu(t) = \lvert -1.65 - 2.12t \rvert$ |
| (8) λ exponential and μ linear | -33.03 | 76.06 | 3.47 | 17.48 | 5.6e-04 | $\boldsymbol{\lambda(t) = 2.42}e^{0.36t}$ | $\boldsymbol{\mu(t) = \lvert 1.03 - 2.96t \rvert}$ |
| (9) λ exponential and μ exponential | -33.80 | 77.59 | 5 | 15.95 | 0.001 | $\lambda(t) = 4.67e^{-0.09t}$ | $\mu(t) = 3.5e^{-0.03t}$ |
| (10) λ linear and μ exponential | -35.8 | 81.61 | 9.02 | 11.93 | 0.008 | $\lambda(t) = \lvert 2.33 - 0.55t \rvert$ | $\mu(t) = 0.01e^{-1.65t}$ |
| (11) λ constant and μ linear | -33.78 | 74.70 | 2.11 | 15.98 | 0.001 | $\lambda = 4.45$ | $\mu(t) = \lvert -3.49 - 0.22t \rvert$ |
| (12) λ constant and μ exponential | -33.79 | 74.73 | 2.14 | 15.95 | 0.001 | $\lambda = 4.52$ | $\mu(t) = 3.63e^{0.049t}$ |
| ***Dianthus* (38/300):** | | | | | | | |
| (1) λ constant (no μ) (Yule) | -41.72 | 85.55 | 23.62 | - | - | $\lambda = 2.50$ | - |
| (2) λ and μ constant | -30.08 | 64.50 | 2.57 | 23.28 | 1.4e-06 | $\lambda = 11.38$ | $\mu = 10.50$ |
| (3) λ linear and no μ | -35.09 | 74.52 | 12.59 | 13.26 | 2.7e-04 | $\lambda(t) = \lvert 5.18 - 1.51t \rvert$ | - |
| (4) λ exponential and no μ | -33.88 | 72.09 | 10.16 | 15.68 | 7.5e-05 | $\boldsymbol{\lambda(t) = 14.76}e^{-1.08t}$ | - |
| (5) λ Linear and μ constant | -29.81 | 66.32 | 4.39 | 23.82 | 6.7e-06 | $\lambda(t) = \lvert 10.91 - 0.68t \rvert$ | $\mu = 8.68$ |
| (6) λ exponential and μ constant | -29.82 | 66.34 | 4.41 | 23.80 | 6.8e-06 | $\lambda(t) = 11.01e^{-0.07t}$ | $\mu = 8.75$ |

| **Models** | logL | AICc | ΔAIC | LRT | LRT p-value | λ value as a function of temperature $\lambda(t) = \lambda_0 + \alpha t$ (linear) $\lambda(t) = \lambda_0 \times e^{\alpha t}$ (exponential) | μ values as a function of temperature $\mu(t) = \mu_0 + \alpha t$ (linear) $\mu(t) = \mu_0 \times e^{\alpha t}$ (exponential) |
|---|---|---|---|---|---|---|---|
| **(7) λ and μ linear** | **-26.36** | **61.93** | **0** | **30.72** | **9.7e-07** | $\boldsymbol{\lambda(t) = \lvert -29.84 + 31.85t \rvert}$ | $\boldsymbol{\mu(t) = \lvert 29.76 - 31.76t \rvert}$ |
| (8) λ exponential and μ linear | -29.39 | 67.99 | 6.06 | 24.66 | 1.8e-05 | $\lambda(t) = 7.13e^{0.24t}$ | $\mu(t) = \lvert -3.52 - 3.72t \rvert$ |
| (9) λ exponential and μ exponential | -28.80 | 66.80 | 4.87 | 25.85 | 1.0e-05 | $\lambda(t) = 5.34e^{0.61t}$ | $\mu(t) = 4.62e^{0.66t}$ |
| (10) λ linear and μ exponential | -35.09 | 79.40 | 17.47 | 13.25 | 0.004 | $\lambda(t) = 5.20 - 1.51t$ | $\mu(t) = 0.95e^{-3.09t}$ |
| (11) λ constant and μ linear | -29.68 | 66.07 | 4.14 | 24.07 | 2.4e-05 | $\lambda = 9.71$ | $\mu(t) = \lvert -7.1 - 0.87t \rvert$ |
| (12) λ constant and μ exponential | -29.68 | 66.06 | 4.13 | 24.08 | 2.4e-05 | $\lambda = 9.74$ | $\mu(t) = 7.35e^{0.098t}$ |

**Table S14** Comparison of different environmental-dependent diversification models applied on major clades inside Caryophylleae using RPANDA package in R for the *rps16* dataset. The best model for each clade is chosen via the corrected Akaike Information Criterion (AICc) and Likelihood Ratio Test (LRT), significant at $P < 0.05$, which is in bold. λ = the speciation rate; μ = the extinction rate (in events/Ma/lineage).

| **Models** | logL | AICc | ΔAIC | LRT | LRT p value | λ value or as a function of time $\lambda(t) = \lambda0 + \alpha t$ (linear) $\lambda(t) = \lambda0 \times e^{\alpha t}$ (exponential) | μ value or as a function of time $\mu(t) = \mu0 + \beta t$ (linear) $\mu(t) = \mu0 \times e^{\beta t}$ (exponential) |
|---|---|---|---|---|---|---|---|
| ***Gypsophila*: (55/151)** | | | | | | | |
| (1) λ constant (no μ) (Yule) | -74.12 | 150.31 | 16.99 | - | - | $\lambda = 1.16$ | - |
| (2) λ and μ constant | -66.63 | 137.49 | 4.17 | 14.97 | 1e-04 | $\lambda = 2.85$ | $\mu = 2.35$ |
| (3) λ linear and no μ | -68.17 | 140.57 | 7.25 | 11.90 | 6e-04 | $\lambda(t) = \lvert 1.95 - 0.33t \rvert$ | - |
| (4) λ exponential and no μ | -67.51 | 139.24 | 5.92 | 13.22 | 3e-04 | $\lambda(t) = 2.93e^{-0.44t}$ | - |
| (5) λ Linear and μ constant | -66.33 | 139.13 | 5.81 | 15.58 | 8e-05 | $\lambda(t) = \lvert 2.76 - 0.16t \rvert$ | $\mu = 1.76$ |
| (6) λ exponential and μ constant | -66.32 | 139.12 | 5.8 | 15.59 | 8e-05 | $\boldsymbol{\lambda(t) = 2.79e^{-0.07t}}$ | $\mu = 1.73$ |
| **(7) λ and μ linear** | **-62.26** | **133.32** | **0** | **23.50** | **1e-06** | $\boldsymbol{\lambda(t) = \lvert 2.88 - 0.34t \rvert}$ | $\boldsymbol{\mu(t) = \lvert 1.60 - 0.11t \rvert}$ |
| (8) λ exponential and μ linear | -65.12 | 139.05 | 5.85 | 17.99 | 2e-05 | $\lambda(t) = 1.06e^{0.41t}$ | $\mu(t) = \lvert 1.88 - 1.93t \rvert$ |
| (9) λ exponential and μ exponential | -66.18 | 141.15 | 7.83 | 15.88 | 7e-05 | $\lambda(t) = 2.03e^{0.26t}$ | $\mu(t) = 1.64e^{0.31t}$ |
| (10) λ linear and μ exponential | -68.26 | 145.32 | 8.47 | 11.71 | 6e-04 | $\lambda(t) = \lvert 2.17 - 0.54t \rvert$ | $\mu(t) = 0.002e^{0.46t}$ |
| (11) λ constant and μ linear | -66.21 | 138.89 | | 15.82 | 7e-05 | $\lambda = 2.51$ | $\mu(t) = \lvert -1.46 - 0.2t \rvert$ |
| (12) λ constant and μ exponential | -66.25 | 138.97 | | 15.74 | 7e-05 | $\lambda = 2.56$ | $\mu(t) = 1.62e^{0.086t}$ |
| ***Dianthus*: (10/300)** | | | | | | | |
| (1) λ constant (no μ) (Yule) | -19.64 | 41.77 | 7.03 | - | - | $\lambda = 1.44$ | - |
| (2) λ and μ constant | -14.69 | 35.09 | 0.35 | 9.90 | 0.002 | $\lambda = 15.22$ | $\mu = 14.90$ |
| (3) λ linear and no μ | -15.18 | 36.07 | 1.33 | 8.91 | 0.003 | $\lambda(t) = \lvert 3.16 - 0.70t \rvert$ | - |
| **(4) λ exponential and no μ** | **-14.51** | **34.74** | **0** | **10.24** | **0.001** | $\boldsymbol{\lambda(t) = 12.60e^{-1.07t}}$ | **-** |
| (5) λ Linear and μ constant | -14.21 | 38.42 | 3.68 | 10.85 | 0.004 | $\lambda(t) = \lvert 9.88 - 0.80t \rvert$ | $\mu = 7.57$ |
| (6) λ exponential and μ constant | -14.24 | 38.48 | 3.74 | 10.79 | 0.004 | $\lambda(t) = 10.42e^{-0.09t}$ | $\mu = 8.06$ |

| | | | | | | | |
|---|---|---|---|---|---|---|---|
| (7) λ and μ linear | -14.35 | 44.71 | 9.97 | 10.56 | 0.014 | $\lambda(t) = \lvert 6.48 + 2.62t \rvert$ | $\mu(t) = \lvert -4.49 - 3.40t \rvert$ |
| (8) λ exponential and μ linear | -14.42 | 44.83 | 10.09 | 10.44 | 0.015 | $\lambda(t) = 6.74e^{0.13t}$ | $\mu(t) = \lvert -4.04 - 2.11t \rvert$ |
| (9) λ exponential and μ exponential | -14.40 | 44.80 | 10.06 | 10.47 | 0.015 | $\lambda(t) = 13.08e^{-0.92t}$ | $\mu(t) = 2.50e^{-0.61t}$ |
| (10) λ linear and μ exponential | -15.18 | 46.37 | 11.63 | 8.90 | 0.031 | $\lambda(t) = \lvert 3.18 - 0.71t \rvert$ | $\mu(t) = 0.84e^{-2.49t}$ |
| (11) λ constant and μ linear | -14.26 | 38.52 | 3.78 | 10.75 | 0.013 | $\lambda = 9.70$ | $\mu(t) = \lvert -7.61 - 0.74t \rvert$ |
| (12) λ constant and μ exponential | -15.68 | 41.37 | 6.63 | 7.90 | 0.048 | $\lambda = 2.12$ | $\mu(t) = 0.11e^{1.02t}$ |
| ***Saponaria*: (7/30)** | | | | | | | |
| **(1) λ constant (no μ) (Yule)** | **-9.79** | **22.38** | **0** | **-** | **-** | $\boldsymbol{\lambda = 0.85}$ | - |
| (2) λ and μ constant | -9.61 | 26.22 | 3.84 | 0.36 | 0.547 | $\lambda = 1.45$ | $\mu = 0.94$ |
| (3) λ linear and no μ | -9.45 | 25.90 | 3.52 | 0.68 | 0.408 | $\lambda(t) = \lvert 1.64 - 0.37t \rvert$ | - |
| (4) λ exponential and no μ | -9.61 | 26.21 | 3.83 | 0.37 | 0.545 | $\lambda(t) = 1.62e^{-0.31t}$ | - |
| (5) λ Linear and μ constant | -9.45 | 32.90 | 10.52 | 0.68 | 0.71 | $\lambda(t) = \lvert 1.64 - 0.37t \rvert$ | $\mu = 2.14e - 06$ |
| (6) λ exponential and μ constant | -9.58 | 33.15 | 10.77 | 0.43 | 0.807 | $\lambda(t) = 1.63e^{-0.16t}$ | $\mu = 0.54$ |
| (7) λ and μ linear | -8.61 | 45.21 | 22.83 | 2.37 | 0.5 | $\lambda(t) = \lvert -4.58 + 4.21t \rvert$ | $\mu(t) = \lvert 6.24 - 5.17t \rvert$ |
| (8) λ exponential and μ linear | -9.13 | 46.26 | 23.88 | 1.32 | 0.723 | $\lambda(t) = 0.54e^{0.57t}$ | $\mu(t) = \lvert 3.07 - 2.48t \rvert$ |
| (9) λ exponential and μ exponential | -8.75 | 45.50 | 23.12 | 2.08 | 0.556 | $\lambda(t) = 0.22e^{1.04t}$ | $\mu(t) = 0.02e^{1.99t}$ |
| (10) λ linear and μ exponential | -9.45 | 46.90 | 24.52 | 0.68 | 0.877 | $\lambda(t) = \lvert 1.64 - 0.37t \rvert$ | $\mu(t) = 0.0003e^{-3.60t}$ |
| (11) λ constant and μ linear | -9.33 | 32.66 | 10.28 | 0.9 | 0.82 | $\lambda = 1.17$ | $\mu(t) = \lvert 1.25 - 0.93t \rvert$ |
| (12) λ constant and μ exponential | -9.15 | 32.30 | 9.92 | 1.28 | 0.73 | $\lambda = 1.04$ | $\mu(t) = (7e - 04)e^{2.67t}$ |
| ***Acanthophyllum*: (10/95)** | | | | | | | |
| (1) λ constant (no μ) (Yule) | -18.47 | 39.44 | 2.94 | - | - | $\lambda = 0.97$ | - |
| (2) λ and μ constant | -15.39 | 36.50 | 0.45 | 6.15 | 0.013 | $\lambda = 5.09$ | $\mu = 4.90$ |
| **(3) λ linear and no μ** | **-15.17** | **36.05** | **0** | **6.60** | **0.010** | $\boldsymbol{\lambda(t) = \lvert 2.19 - 0.45t \rvert}$ | **-** |
| (4) λ exponential and no μ | -15.17 | 36.06 | 0.01 | 6.59 | 0.010 | $\lambda(t) = 4.90e^{-0.72t}$ | - |
| (5) λ Linear and μ constant | -14.74 | 39.72 | 3.67 | 7.46 | 0.024 | $\lambda(t) = \lvert 3.95 - 0.53t \rvert$ | $\mu = 2.20$ |
| (6) λ exponential and μ constant | -14.78 | 39.56 | 3.51 | 7.38 | 0.025 | $\lambda(t) = 4.36e^{-0.20t}$ | $\mu = 2.28$ |
| (7) λ and μ linear | -14.50 | 45.01 | 8.96 | 7.93 | 0.048 | $\lambda(t) = \lvert 1.28 + 1.21t \rvert$ | $\mu(t) = \lvert 0.68 - 2.00t \rvert$ |

| | | | | | | | |
|---|---|---|---|---|---|---|---|
| (8) λ exponential and μ linear | -14.56 | 45.11 | 9.06 | 7.82 | 0.050 | $\lambda(t) = 2.78e^{0.09t}$ | $\mu(t) = \lvert -0.77 - 1.03t \rvert$ |
| (9) λ exponential and μ exponential | -14.73 | 45.45 | 9.4 | 7.48 | 0.058 | $\lambda(t) = 3.26e^{-0.32t}$ | $\mu(t) = 0.30e^{0.43t}$ |
| (10) λ linear and μ exponential | -14.42 | 44.85 | 8.8 | 8.09 | 0.044 | $\lambda(t) = \lvert 2.09 + 0.60t \rvert$ | $\mu(t) = 1.11e^{0.43t}$ |
| (11) λ constant and μ linear | -14.57 | 39.14 | 3.09 | 7.80 | 0.050 | $\lambda = 2.68$ | $\mu(t) = \lvert -0.42 - 0.80t \rvert$ |
| (12) λ constant and μ exponential | -14.50 | 39.01 | 2.96 | 7.93 | 0.047 | $\lambda = 2.64$ | $\mu(t) = 0.95e^{0.35t}$ |

**Table S15** Comparison of different environmental-dependent diversification models applied on major clades within Caryophylleae using the RPANDA package in R for the ITS dataset. The best model for each clade is chosen via the corrected Akaike Information Criterion (AICc) and Likelihood Ratio Test (LRT), significant at P < 0.05, which is in bold. λ = the speciation rate; μ = the extinction rate (in events/Ma/lineage).

| **Models** | logL | AICc | ΔAIC | LRT | LRT p-value | λ value or as a function of time $\lambda(t) = \lambda 0 + \alpha t$ (linear) $\lambda(t) = \lambda 0 \times e^{\alpha t}$ (exponential) | μ value or as a function of time $\mu(t) = \mu 0 + \beta t$ (linear) $\mu(t) = \mu 0 \times e^{\beta t}$ (exponential) |
|---|---|---|---|---|---|---|---|
| ***Gypsophila*: (71/152)** | | | | | | | |
| (1) λ constant (no μ) (Yule) | -123.99 | 250.03 | 10.46 | - | - | $\lambda = 0.67$ | - |
| (2) λ and μ constant | -117.72 | 239.61 | 0.04 | 12.54 | 4e-04 | $\lambda = 1.42$ | $\mu = 1.09$ |
| (3) λ linear and no μ | -118.13 | 240.44 | 0.87 | 11.71 | 6e-04 | $\lambda(t) = \|1.09 - 0.14t\|$ | - |
| **(4) λ exponential and no μ** | **-117.69** | **239.57** | **0** | **12.58** | **4e-04** | $\boldsymbol{\lambda(t) = 1.40e^{-0.29t}}$ | - |
| (5) λ Linear and μ constant | -117.39 | 241.14 | 1.57 | 13.19 | 0.001 | $\lambda(t) = \|1.34 - 0.08t\|$ | $\mu = 0.66$ |
| (6) λ exponential and μ constant | -117.39 | 241.14 | 1.57 | 13.20 | 0.001 | $\lambda(t) = 1.36e^{-0.09t}$ | $\mu = 0.62$ |
| (7) λ and μ linear | -117.52 | 243.64 | 4.07 | 12.94 | 0.005 | $\lambda(t) = \|0.80 + 0.43t\|$ | $\mu(t) = \|-0.51 - 0.46t\|$ |
| (8) λ exponential and μ linear | -117.46 | 243.52 | 3.95 | 13.06 | 0.004 | $\lambda(t) = 0.99e^{0.20t}$ | $\mu(t) = \|-0.28 - 0.47t\|$ |
| (9) λ exponential and μ exponential | -117.49 | 243.59 | 4.02 | 12.99 | 0.005 | $\lambda(t) = 1.76e^{-0.30t}$ | $\mu(t) = 2.44e^{-1.22t}$ |
| (10) λ linear and μ exponential | -117.39 | 243.39 | 3.82 | 13.19 | 0.004 | $\lambda(t) = \|1.33 - 0.08t\|$ | $\mu(t) = 0.66e^{0.005t}$ |
| (11) λ constant and μ linear | -117.41 | 241.17 | 1.6 | 13.16 | 0.004 | $\lambda = 1.26$ | $\mu(t) = \|-0.66 - 0.076t\|$ |
| (12) λ constant and μ exponential | -117.42 | 241.19 | 1.62 | 13.14 | 0.004 | $\lambda = 1.27$ | $\mu(t) = 0.71e^{0.079t}$ |
| ***Dianthus*: (40/300)** | | | | | | | |
| (1) λ constant (no μ) (Yule) | -53.21 | 108.52 | 25.55 | - | - | $\lambda = 2.04$ | - |
| (2) λ and μ constant | -39.73 | 83.79 | 0.82 | 26.95 | 2e-07 | $\lambda = 9.31$ | $\mu = 8.68$ |
| (3) λ linear and no μ | -43.10 | 90.53 | 7.56 | 20.21 | 7e-06 | $\lambda(t) = \|4.63 - 1.26t\|$ | - |
| (4) λ exponential and no μ | -41.16 | 86.65 | 3.68 | 24.08 | 9e-07 | $\boldsymbol{\lambda(t) = 9.64e^{-0.87t}}$ | **-** |
| (5) λ Linear and μ constant | -42.09 | 90.85 | 7.88 | 22.23 | 1e-05 | $\lambda(t) = \|4.73 - 1.39t\|$ | $\mu = 0.19$ |

| | | | | | | | |
|---|---|---|---|---|---|---|---|
| (6) λ exponential and μ constant | -38.78 | 84.22 | 1.25 | 28.86 | 5e-07 | $\lambda(t) = 8.18e^{-0.09t}$ | $\mu = 5.95$ |
| **(7) λ and μ linear** | **36.92** | **82.97** | **0** | **32.58** | **4e-07** | $\boldsymbol{\lambda(t) = \lvert -14.57 + 16.99t \rvert}$ | $\boldsymbol{\mu(t) = \lvert 15.76 - 17.54t \rvert}$ |
| (8) λ exponential and μ linear | -37.50 | 84.15 | 1.18 | 31.41 | 7e-07 | $\lambda(t) = 4.28e^{0.34t}$ | $\mu(t) = \lvert 0.60 - 4.42t \rvert$ |
| (9) λ exponential and μ exponential | -38.77 | 86.69 | 3.72 | 28.86 | 2e-06 | $\lambda(t) = 7.81e^{-0.05t}$ | $\mu(t) = 5.75e^{0.045t}$ |
| (10) λ linear and μ exponential | -39.09 | 87.33 | 4.36 | 28.23 | 3e-06 | $\lambda(t) = \lvert 7.78 - 2.07t \rvert$ | $\mu(t) = 1.86e^{-0.04t}$ |
| (11) λ constant and μ linear | -38.72 | 84.10 | 1.13 | 28.98 | 2e-06 | $\lambda = 7.36$ | $\mu(t) = \lvert -5.29 - 0.61t \rvert$ |
| (12) λ constant and μ exponential | -38.78 | 84.22 | 1.25 | 28.86 | 2e-06 | $\lambda = 7.57$ | $\mu(t) = 5.75e^{0.08t}$ |
| ***Saponaria*: (24/30)** | | | | | | | |
| **(1) λ constant (no μ) (Yule)** | **-47.89** | **97.97** | **0** | - | - | $\boldsymbol{\lambda = 0.35}$ | - |
| (2) λ and μ constant | -47.89 | 100.36 | 2.39 | 0.006 | 0.94 | $\lambda = 0.36$ | $\mu = 0.02$ |
| (3) λ linear and no μ | -47.89 | 100.36 | 2.39 | 0.002 | 0.96 | $\lambda(t) = \lvert 0.33 + 0.004t \rvert$ | - |
| (4) λ exponential and no μ | -47.89 | 100.36 | 2.39 | 0.002 | 0.96 | $\lambda(t) = 0.33e^{0.012t}$ | - |
| (5) λ Linear and μ constant | -47.83 | 102.86 | 4.89 | 0.13 | 0.94 | $\lambda(t) = \lvert 0.33 + 0.05t \rvert$ | $\mu = 0.22$ |
| (6) λ exponential and μ constant | -47.83 | 102.86 | 4.89 | 0.13 | 0.94 | $\lambda(t) = 0.34e^{0.095t}$ | $\mu = 0.21$ |
| (7) λ and μ linear | -47.72 | 105.54 | 7.57 | 0.35 | 0.95 | $\lambda(t) = \lvert 0.17 + 0.16t \rvert$ | $\mu(t) = \lvert -0.15 - 0.12t \rvert$ |
| (8) λ exponential and μ linear | -47.57 | 105.23 | 7.26 | 0.66 | 0.88 | $\lambda(t) = 0.21e^{0.39t}$ | $\mu(t) = \lvert 0.23 - 0.32t \rvert$ |
| (9) λ exponential and μ exponential | -47.54 | 105.19 | 7.22 | 0.71 | 0.87 | $\lambda(t) = 0.23e^{0.42t}$ | $\mu(t) = 0.29e^{0.36t}$ |
| (10) λ linear and μ exponential | -47.77 | 105.65 | 7.68 | 0.24 | 0.97 | $\lambda(t) = \lvert 0.28 + 0.03t \rvert$ | $\mu(t) = 0.0007e^{1.15t}$ |
| (11) λ constant and μ linear | -47.88 | 102.96 | 4.99 | 0.03 | 0.999 | $\lambda = 0.37$ | $\mu(t) = \lvert -0.14 + 0.04t \rvert$ |
| (12) λ constant and μ exponential | -47.90 | 102.99 | 5.02 | -0.003 | 1 | $\lambda = 0.35$ | $\mu(t) = 0.02e^{-0.097t}$ |
| ***Acanthophyllum*: (14/95)** | | | | | | | |
| (1) λ constant (no μ) (Yule) | -28.14 | 58.61 | 0.2 | - | - | $\lambda = 0.66$ | - |
| **(2) λ and μ constant** | **-26.66** | **58.41** | 0 | **2.96** | **0.086** | $\boldsymbol{\lambda = 1.87}$ | $\boldsymbol{\mu = 1.55}$ |
| (3) λ linear and no μ | -27.08 | 59.26 | 0.85 | 2.11 | 0.15 | $\lambda(t) = \lvert 1.15 - 0.16t \rvert$ | - |
| (4) λ exponential and no μ | -27.09 | 59.27 | 0.86 | 2.10 | 0.15 | $\lambda(t) = 1.39e^{-0.26t}$ | - |

| | | | | | | | |
|---|---|---|---|---|---|---|---|
| (5) λ Linear and μ constant | -26.66 | 61.72 | 3.31 | 2.95 | 0.23 | $\lambda(t) = \lvert 1.89 + 0.006t \rvert$ | $\mu = 1.59$ |
| (6) λ exponential and μ constant | -26.66 | 61.72 | 3.31 | 2.95 | 0.23 | $\lambda(t) = 1.88e^{0.003t}$ | $\mu = 1.58$ |
| (7) λ and μ linear | -26.59 | 65.63 | 7.22 | 3.09 | 0.38 | $\lambda(t) = \lvert 1.10 + 0.70t \rvert$ | $\mu(t) = \lvert -1.08 - 0.67t \rvert$ |
| (8) λ exponential and μ linear | -26.64 | 65.72 | 7.31 | 3.00 | 0.39 | $\lambda(t) = 1.68e^{0.08t}$ | $\mu(t) = \lvert -1.34 - 0.17t \rvert$ |
| (9) λ exponential and μ exponential | -26.72 | 65.88 | 7.47 | 2.84 | 0.42 | $\lambda(t) = 3.78e^{-0.37t}$ | $\mu(t) = 9.60e^{-1.08t}$ |
| (10) λ linear and μ exponential | -26.51 | 65.47 | 7.06 | 3.25 | 0.35 | $\lambda(t) = \lvert 1.55 + 0.51t \rvert$ | $\mu(t) = 1.81e^{0.15t}$ |
| (11) λ constant and μ linear | -27.01 | 62.42 | 4.01 | 2.25 | 0.521 | $\lambda = 1.21$ | $\mu(t) = \lvert -0.82 - 0.02t \rvert$ |
| (12) λ constant and μ exponential | -26.66 | 61.72 | 3.31 | 2.96 | 0.398 | $\lambda = 1.81$ | $\mu(t) = 1.44e^{0.01t}$ |

**Table S16** Comparison of different historical biogeography models for ancestral area estimations of Caryophylleae phylogeny of all three datasets estimated in BIOGEOBEARS. log-likelihood value (lnL), number of parameters (n), rate of range expansion (d), rate of range contraction (e), relative weight of jump dispersal at cladogenesis (j), corrected Akaike's information criteria (AICc), and AICc weight (AICc_wt). The best model for each dataset is shown in bold.

*mat*K:

| Model | Ln L | n | d | e | j | AICc | AICc_wt |
|---|---|---|---|---|---|---|---|
| DEC | -297.5 | 2 | 0.64 | 2.13 | 0 | 599.1 | 0.0014 |
| DEC + j | -297.5 | 3 | 0.67 | 2.25 | 1.0e-05 | 601.2 | 0.0005 |
| DIVALIKE | -315.3 | 2 | 0.20 | 0.47 | 0 | 634.8 | 2.5e-11 |
| DIVALIKE + j | -299.6 | 3 | 0.87 | 2.99 | 1.0e-05 | 605.5 | 5.7e-05 |
| BAYAREALIKE | -294.8 | 2 | 0.40 | 1.54 | 0 | 593.8 | 0.019 |
| **BAYAREALIKE + j** | **-289.9** | **3** | **0.18** | **0.66** | **0.029** | **586** | **0.98** |

*rps16*:

| Model | Ln L | n | d | e | j | AICc | AICc_wt |
|---|---|---|---|---|---|---|---|
| DEC | -231.1 | 2 | 0.11 | 0.27 | 0 | 466.2 | 2.1e-05 |
| DEC + j | -230.4 | 3 | 0.12 | 0.33 | 0.0042 | 467 | 1.4e-05 |
| DIVALIKE | -234.8 | 2 | 0.16 | 0.50 | 0 | 473.8 | 4.8e-07 |
| DIVALIKE + j | -234.8 | 3 | 0.17 | 0.56 | 1.0e-05 | 475.8 | 1.8e-07 |
| BAYAREALIKE | -224.7 | 2 | 0.16 | 0.78 | 0 | 453.6 | 0.011 |
| **BAYAREALIKE + j** | **-219.2** | **3** | **0.080** | **0.32** | **0.018** | **444.7** | **0.99** |

ITS:

| Model | Ln L | n | d | e | j | AICc | AICc_wt |
|---|---|---|---|---|---|---|---|
| DEC | -471.7 | 2 | 0.099 | 0.090 | 0 | 947.5 | 3.1e-18 |
| DEC + j | -463 | 3 | 0.14 | 0.38 | 0.0005 | 932.2 | 6.2e-15 |
| DIVALIKE | -485.3 | 2 | 0.15 | 0.27 | 0 | 974.6 | 3.9e-24 |
| DIVALIKE + j | -479.2 | 3 | 0.22 | 0.74 | 1.0e-05 | 964.5 | 6.0e-22 |
| BAYAREALIKE | -460.6 | 2 | 0.24 | 0.24 | 0 | 925.2 | 2.0e-13 |
| **BAYAREALIKE + j** | **-430.3** | **3** | **0.22** | **0.22** | **0.017** | **866.8** | **1.00** |

**Table S17** ancestral area reconstructions of Caryophylleae and major clades inside it for the *mat*K dataset using three methods implemented in RASP.

| *mat*K | | | | | | | | | |
|---|---|---|---|---|---|---|---|---|---|
| Nodes | BAYAREALIKE + j (Probability) | | | DEC (Probability) | | | BBM (Probability) | | |
| Caryophylleae | ABCDE 5.54 | ABCE 4.73 | ABDE 4.72 | ABCDE 3.52 | ABCE 3.49 | ABDE 3.49 | B 41.63 | C 17.85 | A 9.91 |
| *Gypsophila*-stem group (*Gypsophila* + *Saponaria*) | ABCDE 10.35 | ABDE 5.35 | ACDE 5.21 | ABCDE 45.57 | ABDE 10.99 | ABCE 10.97 | AE 32.92 | A 23.25 | E 16.08 |
| *Gypsophila*-crown group | AB 11.45 | A 10.94 | ABD 7.87 | ABCDE 46.29 | ABDE 12.52 | ABCD 11.36 | A 42.52 | AE 18.48 | AB 16.33 |
| *Dianthus*-stem group (*Dianthus* + *Petrorhagia*) | ABCDE 13.28 | ABDE 10.15 | ADE 8.45 | E 11.86 | ABCDE 9.55 | D 7.51 | ADE 58.43 | ABDE 32.17 | ACDE 2.17 |
| *Dianthus*-crown + *Velezia* | ADE 13.71 | AE 10.39 | ABDE 8.10 | ABCDE 34.19 | ABDE 11.87 | ABCE 10.02 | ABDE 37.64 | ADE 34.11 | ABCDE 8.28 |
| *Saponaria* | E 29.75 | DE 16.00 | AE 10.09 | ABCDE 29.58 | E 11.59 | ABDE 11.02 | E 63.57 | AE 23.47 | DE 6.19 |
| *Acanthophyllum* | B 21.20 | AB 17.26 | A 7.42 | ABCDE 17.29 | B 12.00 | AB 9.80 | AB 65.46 | A 14.58 | B 11.87 |
| Global Cost: | Global Dispersal: 154 Global Vicariance: 22 Global Extinction: 7 | | | Global Dispersal: 136 Global Vicariance: 19 Global Extinction: 16 | | | Global Dispersal: 149 Global Vicariance: 19 Global Extinction: 2 | | |

**Table S18** ancestral area reconstructions for Caryophylleae and major clades inside it for the *rps16* dataset using three methods implemented in RASP.

| *rps16* | | | | | | | | | |
|---|---|---|---|---|---|---|---|---|---|
| | BAYAREALIKE + j | | | DEC | | | BBM | | |
| Caryophylleae | A 15.77 | AE 8.13 | E 7.96 | A 6.25 | AB 4.88 | AC 4.74 | B 12.94 | C 8.70 | A 8.49 |
| *Gypsophila*-stem group (*Gypsophila* + *Saponaria*) | A 43.42 | E 14.23 | AE 12.59 | A 25.14 | ABCDE 17.43 | AE 8.53 | AE 62.58 | A 20.63 | ADE 5.02 |
| *Gypsophila*-crown group | A 80.12 | AE 16.01 | E 1.29 | A 100.00 | - | - | AE 65.48 | A 18.58 | ABE 6.09 |
| *Dianthus*-stem group (*Dianthus* + *Petrorhagia* + *Graecobolanthus*) | AE 49.54 | E 28.66 | ADE 8.33 | E 100.00 | - | - | ADE 87.22 | AE 9.65 | ABDE 1.26 |
| *Dianthus*-crown + *Velezia* | AE 45.77 | E 24.56 | ADE 11.88 | E 86.60 | DE 13.40 | - | ADE 83.61 | AE 13.33 | DE 1.04 |
| *Saponaria* | DE 45.15 | E 18.03 | A 15.056 | ABDE 52.95 | ABCDE 26.37 | ABE 12.82 | AE 47.42 | A 14.81 | ADE 11.80 |
| *Acanthophyllum* | A 71.90 | B 12.75 | AE 5.01 | A 55.92 | AB 44.08 | - | A 66.03 | AB 4.92 | AE 4.42 |
| *Bolanthus* (*Graecobolanthus* clade) | E 99.19 | AE 0.55 | DE 0.19 | E 100.00 | - | - | E 79.74 | DE 12.96 | AE 6.04 |
| *Bolanthus* (*Phrynella* clade) | AE 98.56 | E 0.93 | A 0.15 | AE 55.95 | E 44.05 | - | AE 97.77 | ADE 1.84 | E 0.15 |
| Global Cost: | Global Dispersal: 92<br>Global Vicariance: 20<br>Global Extinction: 0 | | | Global Dispersal: 83<br>Global Vicariance: 14<br>Global Extinction: 0 | | | Global Dispersal: 103<br>Global Vicariance: 23<br>Global Extinction: 0 | | |

**Table S19** ancestral area reconstructions for Caryophylleae and major clades inside it for the ITS dataset using three methods implemented in RASP.

| ITS: | | | | | | | | | |
|---|---|---|---|---|---|---|---|---|---|
| Major clades | BAYAREALIKE + j | | | DEC | | | BBM | | |
| Caryophylleae | AE<br>9.58 | A<br>7.65 | AB<br>5.45 | AB<br>10.55 | AC<br>10.25 | A<br>10.09 | B<br>12.11 | A<br>11.04 | C<br>7.08 |
| *Gypsophila*-Stem group (*Gypsophila* + all genera except *Psammosilene*) | A<br>21.98 | AE<br>18.15 | E<br>5.85 | A<br>57.16 | AE<br>16.30 | E<br>14.01 | AE<br>43.06 | A<br>39.68 | E<br>4.08 |
| *Gypsophila*-Crown group | AE<br>40.99 | A<br>39.00 | ABE<br>3.77 | A<br>60.82 | ABCDE<br>25.76 | AE<br>13.42 | AE<br>82.11 | A<br>9.47 | ABE<br>6.02 |
| *Dianthus*-Stem group (*Dianthus*+*Petrorhagia* +*Graecobolanthus*+*Bolanthus*) | AE<br>68.96 | E<br>14.92 | ADE<br>9.31 | E<br>87.86 | AE<br>12.14 | - | AE<br>69.05 | E<br>18.39 | ADE<br>9.11 |
| *Dianthus*-Crown + *Velezia* | ADE<br>49.11 | AE<br>40.18 | DE<br>5.70 | E<br>49.91 | ABCDE<br>16.53 | AE<br>15.52 | AE<br>58.74 | ADE<br>35.24 | E<br>2.33 |
| *Saponaria* | DE<br>18.31 | AE<br>17.73 | E<br>16.97 | AE<br>31.81 | E<br>23.80 | ABE<br>11.31 | AE<br>41.78 | A<br>33.77 | E<br>8.68 |
| *Acanthophyllum* | A<br>60.24 | AB<br>20.13 | B<br>8.68 | A<br>46.87 | AB<br>46.03 | ABE<br>7.10 | A<br>80.53 | AB<br>14.54 | AE<br>2.42 |
| *Bolanthus* (*Graecobolanthus* clade) | E<br>99.17 | AE<br>0.51 | DE<br>0.14 | E<br>100.00 | - | - | E<br>93.70 | AE<br>3.32 | DE<br>2.68 |
| *Bolanthus* (*Phrynella* clade) | AE<br>98.67 | ADE<br>0.45 | E<br>0.43 | E<br>67.61 | AE<br>32.39 | - | AE<br>93.28 | ADE<br>4.07 | E<br>2.36 |
| *Petrorhagia* | AE<br>51.91 | ADE<br>21.50 | E<br>16.35 | E<br>100.00 | - | - | AE<br>27.68 | ADE<br>26.38 | E<br>23.26 |
| Global Cost: | Global Dispersal: 210<br>Global Vicariance: 31<br>Global Extinction: 2 | | | Global Dispersal: 164<br>Global Vicariance: 21<br>Global Extinction: 3 | | | Global Dispersal: 218<br>Global Vicariance: 26<br>Global Extinction: 0 | | |

**Table S20** Bayes Factor matrix for models with up to 12 diversification rate shifts compared to a null model with 0 shifts as inferred from the *mat*K dataset for Caryophylleae.

| | 0.00 | 1.00 | 2.00 | 3.00 | 4.00 | 5.00 | 6.00 | 7.00 | 8.00 | 9.00 | 10.00 | 12.00 |
|---|---|---|---|---|---|---|---|---|---|---|---|---|
| 0.00 | 1.00 | 0.00 | 0.00 | 0.00 | 0.00 | 0.00 | 0.00 | 0.00 | 0.00 | 0.00 | 0.00 | 0.00 |
| 1.00 | 240.00 | 1.00 | 0.07 | 0.03 | 0.02 | 0.02 | 0.02 | 0.02 | 0.02 | 0.03 | 0.03 | 0.03 |
| 2.00 | 3464.00 | 14.43 | 1.00 | 0.41 | 0.28 | 0.26 | 0.27 | 0.31 | 0.33 | 0.38 | 0.38 | 0.42 |
| 3.00 | 8520.00 | 35.50 | 2.46 | 1.00 | 0.68 | 0.65 | 0.66 | 0.77 | 0.81 | 0.92 | 0.92 | 1.04 |
| 4.00 | 12464.00 | 51.93 | 3.60 | 1.46 | 1.00 | 0.95 | 0.96 | 1.12 | 1.19 | 1.35 | 1.35 | 1.52 |
| 5.00 | 13152.00 | 54.80 | 3.80 | 1.54 | 1.06 | 1.00 | 1.01 | 1.18 | 1.25 | 1.43 | 1.43 | 1.61 |
| 6.00 | 12992.00 | 54.13 | 3.75 | 1.52 | 1.04 | 0.99 | 1.00 | 1.17 | 1.24 | 1.41 | 1.41 | 1.59 |
| 7.00 | 11136.00 | 46.40 | 3.21 | 1.31 | 0.89 | 0.85 | 0.86 | 1.00 | 1.06 | 1.21 | 1.21 | 1.36 |
| 8.00 | 10496.00 | 43.73 | 3.03 | 1.23 | 0.84 | 0.80 | 0.81 | 0.94 | 1.00 | 1.14 | 1.14 | 1.28 |
| 9.00 | 9216.00 | 38.40 | 2.66 | 1.08 | 0.74 | 0.70 | 0.71 | 0.83 | 0.88 | 1.00 | 1.00 | 1.13 |
| 10.00 | 9216.00 | 38.40 | 2.66 | 1.08 | 0.74 | 0.70 | 0.71 | 0.83 | 0.88 | 1.00 | 1.00 | 1.13 |
| 12.00 | 8192.00 | 34.13 | 2.36 | 0.96 | 0.66 | 0.62 | 0.63 | 0.74 | 0.78 | 0.89 | 0.89 | 1.00 |

**Table S21** Bayes Factor matrix for models with up to 15 diversification rate shifts compared to a null model with 0 shifts as inferred from the *rps16* dataset for Caryophylleae.

| | 0.00 | 1.00 | 2.00 | 3.00 | 4.00 | 5.00 | 6.00 | 7.00 | 8.00 | 9.00 | 10.00 | 11.00 | 12.00 | 13.00 | 15.00 |
|---|---|---|---|---|---|---|---|---|---|---|---|---|---|---|---|
| 0.00 | 1.00 | 0.24 | 0.09 | 0.02 | 0.01 | 0.00 | 0.00 | 0.00 | 0.00 | 0.00 | 0.00 | 0.00 | 0.00 | 0.00 | 0.00 |
| 1.00 | 4.12 | 1.00 | 0.39 | 0.10 | 0.02 | 0.01 | 0.00 | 0.00 | 0.00 | 0.00 | 0.00 | 0.00 | 0.01 | 0.01 | 0.01 |
| 2.00 | 10.62 | 2.58 | 1.00 | 0.25 | 0.06 | 0.02 | 0.01 | 0.01 | 0.01 | 0.01 | 0.01 | 0.01 | 0.01 | 0.02 | 0.02 |
| 3.00 | 41.69 | 10.13 | 3.93 | 1.00 | 0.22 | 0.08 | 0.05 | 0.04 | 0.04 | 0.04 | 0.04 | 0.04 | 0.06 | 0.09 | 0.07 |
| 4.00 | 189.54 | 46.06 | 17.86 | 4.55 | 1.00 | 0.36 | 0.22 | 0.17 | 0.17 | 0.18 | 0.20 | 0.19 | 0.27 | 0.40 | 0.30 |
| 5.00 | 524.31 | 127.40 | 49.39 | 12.58 | 2.77 | 1.00 | 0.61 | 0.48 | 0.46 | 0.51 | 0.55 | 0.53 | 0.74 | 1.11 | 0.83 |
| 6.00 | 856.62 | 208.15 | 80.70 | 20.55 | 4.52 | 1.63 | 1.00 | 0.78 | 0.75 | 0.83 | 0.91 | 0.87 | 1.21 | 1.81 | 1.36 |
| 7.00 | 1097.85 | 266.77 | 103.42 | 26.33 | 5.79 | 2.09 | 1.28 | 1.00 | 0.96 | 1.06 | 1.16 | 1.12 | 1.55 | 2.32 | 1.74 |
| 8.00 | 1142.15 | 277.53 | 107.59 | 27.39 | 6.03 | 2.18 | 1.33 | 1.04 | 1.00 | 1.10 | 1.21 | 1.16 | 1.61 | 2.42 | 1.81 |
| 9.00 | 1033.85 | 251.21 | 97.39 | 24.80 | 5.45 | 1.97 | 1.21 | 0.94 | 0.91 | 1.00 | 1.09 | 1.05 | 1.46 | 2.19 | 1.64 |
| 10.00 | 945.23 | 229.68 | 89.04 | 22.67 | 4.99 | 1.80 | 1.10 | 0.86 | 0.83 | 0.91 | 1.00 | 0.96 | 1.33 | 2.00 | 1.50 |
| 11.00 | 984.62 | 239.25 | 92.75 | 23.62 | 5.19 | 1.88 | 1.15 | 0.90 | 0.86 | 0.95 | 1.04 | 1.00 | 1.39 | 2.08 | 1.56 |
| 12.00 | 708.92 | 172.26 | 66.78 | 17.00 | 3.74 | 1.35 | 0.83 | 0.65 | 0.62 | 0.69 | 0.75 | 0.72 | 1.00 | 1.50 | 1.13 |
| 13.00 | 472.62 | 114.84 | 44.52 | 11.34 | 2.49 | 0.90 | 0.55 | 0.43 | 0.41 | 0.46 | 0.50 | 0.48 | 0.67 | 1.00 | 0.75 |
| 15.00 | 630.15 | 153.12 | 59.36 | 15.11 | 3.32 | 1.20 | 0.74 | 0.57 | 0.55 | 0.61 | 0.67 | 0.64 | 0.89 | 1.33 | 1.00 |

**Table S22** Bayes Factor matrix for models with up to 10 diversification rate shifts compared to a null model with 0 shifts as inferred from the ITS dataset for Caryophylleae.

| | 0.00 | 1.00 | 2.00 | 3.00 | 4.00 | 5.00 | 6.00 | 7.00 | 8.00 | 9.00 | 10.00 |
|---|---|---|---|---|---|---|---|---|---|---|---|
| 0.00 | 1.00 | 0.01 | 0.00 | 0.00 | 0.00 | 0.00 | 0.00 | 0.00 | 0.00 | 0.00 | 0.00 |
| 1.00 | 146.00 | 1.00 | 0.09 | 0.01 | 0.01 | 0.01 | 0.01 | 0.01 | 0.01 | 0.02 | 0.02 |
| 2.00 | 1600.00 | 10.96 | 1.00 | 0.16 | 0.10 | 0.10 | 0.12 | 0.15 | 0.15 | 0.22 | 0.20 |
| 3.00 | 10296.00 | 70.52 | 6.44 | 1.00 | 0.67 | 0.62 | 0.79 | 0.95 | 0.94 | 1.44 | 1.26 |
| 4.00 | 15472.00 | 105.97 | 9.67 | 1.50 | 1.00 | 0.93 | 1.19 | 1.42 | 1.41 | 2.16 | 1.89 |
| 5.00 | 16672.00 | 114.19 | 10.42 | 1.62 | 1.08 | 1.00 | 1.28 | 1.53 | 1.51 | 2.33 | 2.04 |
| 6.00 | 12992.00 | 88.99 | 8.12 | 1.26 | 0.84 | 0.78 | 1.00 | 1.19 | 1.18 | 1.81 | 1.59 |
| 7.00 | 10880.00 | 74.52 | 6.80 | 1.06 | 0.70 | 0.65 | 0.84 | 1.00 | 0.99 | 1.52 | 1.33 |
| 8.00 | 11008.00 | 75.40 | 6.88 | 1.07 | 0.71 | 0.66 | 0.85 | 1.01 | 1.00 | 1.54 | 1.34 |
| 9.00 | 7168.00 | 49.10 | 4.48 | 0.70 | 0.46 | 0.43 | 0.55 | 0.66 | 0.65 | 1.00 | 0.88 |
| 10.00 | 8192.00 | 56.11 | 5.12 | 0.80 | 0.53 | 0.49 | 0.63 | 0.75 | 0.74 | 1.14 | 1.00 |

**Table S23** Optimal number shift in diversification rate for the *mat*K dataset calculated by the stepBF function in R (BAMMtools package) using the Bayes factor, as recommended by BAMM authors. We increased the value of the step size to reduce the type I error in the cost of increasing type II error (which translates to inferring fewer shifts).

| | Step.size = 1 (any increase in bayes factor considered a shift) | Step.size = 2 (duplicate increase in bayes factor considered a shift) | Step.size = 3 (duplicate increase in bayes factor considered a shift) | Step.size = 4 (increase by the factor of 4 considered a shift) | Step.size = 14 (increase by the factor of 14 considered a shift) | Step.size = 15 (increase by the factor of 15 considered a shift) |
|---|---|---|---|---|---|---|
| Best model (number of shifts) | 6 | 4 | 3 | 3 | 3 | 2 |
| Second model (number of shifts) | 5 | 3 | 2 | 2 | 2 | 0 |
| Bayes factor | 1.05 | 2.46 | 14.43 | 14.43 | 14.43 | 0 |

**Table S24** Optimal number shift in diversification rate for the *rps16* dataset calculated by the stepBF function in R (BAMMtools package) using Bayes factor, as recommended by BAMM authors. We increased the value of the step size to reduce the type I error in the cost of increasing type II error (which translates to inferring fewer shifts).

| | Step.size = 1 (any increase in bayes factor considered a shift) | Step.size = 2 (duplicate increase in bayes factor considered a shift) | Step.size = 3 (duplicate increase in bayes factor considered a shift) | Step.size = 4 (increase by the factor of 4 considered a shift) | Step.size = 5 (increase by the factor of 5 considered a shift) | Step.size = 6 (increase by the factor of 6 considered a shift) |
|---|---|---|---|---|---|---|
| Best model (number of shifts) | 8 | 5 | 1 | 1 | 0 | 0 |
| Second model (number of shifts) | 7 | 4 | 0 | 0 | 0 | 0 |
| Bayes factor | 1.04 | 2.76 | 4.11 | 4.11 | 0 | 0 |

**Table S25** Optimal number shift in diversification rate for the ITS dataset calculated by the stepBF function in R (BAMMtools package) using Bayes factor, as recommended by BAMM authors. We increased the value of the step size to reduce the type I error in the cost of increasing type II error (which translates to inferring fewer shifts).

| | Step.size = 1 (any increase in bayes factor considered a shift) | Step.size = 2 (duplicate increase in bayes factor considered a shift) | Step.size = 3 (duplicate increase in bayes factor considered a shift) | Step.size = 7 (increase by the factor of 7 considered a shift) | Step.size = 11 (increase by the factor of 11 considered a shift) |
|---|---|---|---|---|---|
| Best model (number of shifts) | 5 | 3 | 3 | 2 | 1 |
| Second model (number of shifts) | 4 | 2 | 2 | 1 | 0 |
| Bayes factor | 1.08 | 6.43 | 6.43 | 10.96 | 0 |

**Table S26** List of all accepted species of *Gypsophila*, based on POWO. The state of each studied trait in diversification analyses is presented for all species.

| All accepted *Gypsophila* species | Herbaceous vs. Woody | Calyx shape (Tubiform vs. Campanulate turbinate) | Montane vs. non-Montane | Habitat | Annual vs. Perennial | Altitude | High elevation vs. Low elevation | Distribution | GeoSSE | Present in our analyses |
|---|---|---|---|---|---|---|---|---|---|---|
| *Gypsophila acantholimoides* Bornm. | W | CT | M | Arid hills | P | 2300 – 2600 | H | Iran | 1 | Yes |
| *Gypsophila acutifolia* Steven ex Spreng. | W | CT | M | Rocky hills, dry stony hills | P | 1000 – 2000? | H | W. Ukraine, Caucasus | 1 | Yes |
| *Gypsophila adenophora* Boiss. & Buhse | H | CT | M | NA | P | NA | NA | SW. Iran | 1 | No |
| *Gypsophila adenophylla* Barkoudah | W | CT | M | Limestone rocks and rocky soil | P | 3000 – 4000 | H | E. Turkey | 1 | No |
| *Gypsophila afghanica* Kandemir & Ghaz. | W | T | M | On granite slopes, and amongst granite outcrops and rocks | P | 3400 – 3600 | H | Afghanistan | 1 | No |
| *Gypsophila altissima* L. | W | CT | M | Calcareous and rocky hills, steppe meadows | P | NA | NA | Ukraine to Mongolia | 0 | Yes |
| *Gypsophila alvandica* Falat., F.Ghahrem. & Assadi | W | CT | M | NA | P | NA | NA | Iran | 1 | No |
| *Gypsophila anatolica* Boiss. & Heldr. | W | CT | N | NA | P | 650 – 1500 | L | Turkey to Syria and Iran | 1 | No |
| *Gypsophila arabica* Barkoudah (This name is a synonym of *G.* | H | CT | N | Desertic sandy soil | P | 10 – 1100 | L | S. Turkey, Syria, E. Iraq, | 1 | Yes |

| | | | | | | | | | | |
|---|---|---|---|---|---|---|---|---|---|---|
| *capillaris* subsp. *confusa*) | | | | | | | | Palestine, and N. Saudi Arabia | | |
| *Gypsophila aretioides* Boiss. | W | CT | M | Calcareous rocks and rocky slope | P | 2000 – 3500 | H | Transcaucasus to Iran and Turkmenistan | 1 | Yes |
| *Gypsophila arrostii* Guss. | H | CT | N | Stony arid calcareous soil | P | 1000> | L | Italy, Sicilia, Turkey | 0 | Yes |
| *Gypsophila arsusiana* (Kotschy ex Boiss.) F.N.Williams | W | T | N | On rocky soil | P | up to 1000 | L | S. Turkey (Hatay) | 1 | No |
| *Gypsophila aucheri* Boiss. | W | CT | M | Shaley slopes and banks | P | 1000 – 2000 (1600>) | H | Lebanon-Syria, Turkey | 1 | Yes |
| *Gypsophila aulieatensis* B.Fedtsch. | W | CT | M | Dry calcareous hills | P | NA | NA | Kazakhstan | 2 | No |
| *Gypsophila australis* (Schltdl.) A.Gray | H | T | N | Sandy soil | A | NA | NA | New South Wales, Victoria, Western Australia | 2 | No |
| *Gypsophila baytopiorum* Kit Tan | NA | NA | NA | NA | P | NA | NA | Turkey | 1 | No |
| *Gypsophila bazorganica* Rech.f. | W | CT | M | Calcareous rocks and rocky slope | P | 2000 – 2500 | H | NW. Iran | 1 | Yes |
| *Gypsophila bellidifolia* Boiss. | H(?) | CT | N(?) | Calcareous Hills | P | 100 – 600 | L | SE. Arabian Peninsula, S. Iran to W. Pakistan | 0 | No |
| *Gypsophila bermejoi* G.López | H | CT | N(?) | Saline soil and ruderal areas in central Spain | P | 1000> | L | N. Central & Central Spain | 2 | Yes |

| | | | | | | | | | | |
|---|---|---|---|---|---|---|---|---|---|---|
| *Gypsophila bicolor* (Freyn & Sint.) Grossh. | H | CT | M | Dry steppe, sandy hills, open forests, fallow fields | P | 1000 – 2000 | H | E. Turkey to Central Asia and Afghanistan | 0 | Yes |
| *Gypsophila bitlisensis* Barkoudah | H | CT | N | Steppes | A | 1800 – 2000 | H | E. Turkey | 1 | Yes |
| *Gypsophila brachypetala* Trautv. | W | CT | M | South-facing slopes in the alpine zone | P | NA | NA | N. & NE. Turkey | 1 | No |
| *Gypsophila briquetiana* Schischk. | W | CT | M | In rocks crevices | P | 2000 – 2500 | H | E. Turkey | 1 | No |
| *Gypsophila bucharica* B.Fedtsch. | H | T | M | Dry hills | P | 2000 | H | Central Asia | 2 | Yes |
| *Gypsophila capillaris* (Forssk.) C.Chr. | H | CT | N | Desertic sandy soil | P | 10 – 1100 | L | SE. Turkey to E. Egypt and NE. Arabian Peninsula | 0 | Yes |
| *Gypsophila capitata* M.Bieb. | H | CT | M | Stony hills, calcareous rocks, gravel | P | 2000? | H | Caucasus | 1 | Yes |
| *Gypsophila capituliflora* Rupr. | W | CT | M | Rocky, stony hills in the alpine zone | P | 3000-4000 | H | Central Asia to W. Mongolia and N. China | 2 | Yes |
| *Gypsophila caricifolia* Boiss. | W | CT | M | Mountain habitats along streams | P | 500-2500 | H | NE. Iraq to W. & Central Iran | 1 | Yes |
| *Gypsophila cephalotes* (Schrenk) F.N.Williams | H | CT | M | Rocky hills, alpine and subalpine meadows and hills | P | 1800-3040 | H | Central Asia | 2 | Yes |

| | | | | | | | | | |
|---|---|---|---|---|---|---|---|---|---|
| *Gypsophila coelesyriaca* (Boiss. & Hausskn.) F.N.Williams | W | T | M | On dry hills and alcareous rocks | P | NA | NA | Syria to Sinai and Iran | 0 | No |
| *Gypsophila collina* Steven ex Ser. | H | CT | M | Stony and dry Hills | P | NA | NA | Central Romania, SW. Ukraine to Krym | 2 | No |
| *Gypsophila curvifolia* Fenzl | W | CT | M | Rocky slopes | P | 1000-3700 (3660?) | H | S. Turkey | 1 | Yes |
| *Gypsophila damascena* Boiss. | W | CT | N | Dry hills, fallow fields | P | NA | NA | Lebanon-Syria | 2 | No |
| *Gypsophila davisii* Barkoudah | W | CT | M | Rocky, stony hills in the alpine zone | P | 1700-2000 | H | SW. Turkey | 1 | No |
| *Gypsophila davurica* Turcz. ex Fenzl | H | CT | M | Steppe grassland, dry stony hills | P | 2500 | H | S. Siberia to N. China | 2 | Yes |
| *Gypsophila diffusa* Fisch. & C.A.Mey. ex Rupr. | H | CT | M | Stony, mostly calcareous hills | P | NA | NA | Caucasus to Central Asia and Central Iran | 0 | No |
| *Gypsophila dumanii* Armagan & E.G.Cakir | W | CT | M | NA | P | 2230 | H | E. Turkey | 1 | No |
| *Gypsophila elegans* M.Bieb. | H | CT | M | Stony hills, gravel banks, and in fields | A | 500-2017 | L(?) | S. Ukraine to W. & N. Iran | 0 | Yes |
| *Gypsophila erikii* Yild. | NA | NA | NA | NA | NA | NA | NA | Turkey | 1 | No |
| *Gypsophila eriocalyx* Boiss. | W | CT | N | Gypsum steppes | P | 800-1000 | L | Turkey | 1 | Yes |
| *Gypsophila farsensis* Falat., Assadi & F.Ghahrem. | H | T | M | In rocky and soily mountains and steppes | P | 1200-1900 | H | Iran, Fars | 1 | No |

| | | | | | | | | | | |
|---|---|---|---|---|---|---|---|---|---|---|
| *Gypsophila fastigiata* L. | H | CT | N | Rocky and calcareous slopes, sandy soil, pine forest | P | NA | NA | Germany to Central Ukraine | 2 | Yes |
| *Gypsophila fedtschenkoana* Schischk. | H | CT | M | Stony hills | P | 1600 | L | Central Asia to NE. Afghanistan | 2 | No |
| *Gypsophila festucifolia* Hub.-Mor. | W | CT | M | Calcareous or limestone slopes | P | 1400-2200 | H | Turkey | 1 | No |
| *Gypsophila floribunda* (Kar. & Kir.) Turcz. ex Fenzl | H | T | M | NA | A | NA | NA | Iran to SW. Siberia and Pakistan | 0 | No |
| *Gypsophila germanicopolitana* Hub.-Mor. | H | CT | M | Gypsum hills | P | 700-800 | L | Turkey | 1 | No |
| *Gypsophila glandulosa* (Boiss.) Walp. | W | CT | M | Stony slopes | P | 1800-2400 | H | North Caucasus, Transcaucasus, Turkey | 1 | No |
| *Gypsophila globulosa* Steven. | W | CT | M | Rocky calcareous hills, rocks | P | 2000> | H | Bulgaria to N. Caucasus | 0 | Yes |
| *Gypsophila glomerata* Pall. ex Adams | W | CT | M | Calcareous rocks, dry stony hills, sandy soil | P | 2000> | H | Bulgaria to N. Caucasus | 0 | Yes |
| *Gypsophila graminifolia* Barkoudak | W | CT | M | Alpine serpentine stone | P | 2700 | H | E. Turkey to NW. Iran | 1 | Yes |
| *Gypsophila guvengorkii* Armagan, Özgökçe & A.Çelik | W | CT | M | Rocks crevices | P | 2000 | H | Turkey | 1 | No |
| *Gypsophila gypsophiloides* (Fenzl) Blakelock | W | T | N | On dry hills, river banks and shallow | P | 700-1400 | L | E. Medit. to Iran | 1 | Yes |

| | | | | | | | | | | |
|---|---|---|---|---|---|---|---|---|---|---|
| | | | | soil, and on calcareous rocks | | | | | | |
| *Gypsophila hakkiarica* Kit Tan | W | CT | M | Alpine serpentine stone | P | 2700-3000 | H | Turkey | 1 | No |
| *Gypsophila heteropoda* Freyn | H | CT | M | Sandy hills, subdesertic sandy soil | A | 1100-1600 | L | Caucasus to W. & Central Asia | 0 | Yes |
| *Gypsophila hispida* Boiss. | H | CT | M | Stony and dry clay hills | P | 2000-2500 | H | Transcaucasus, Turkey | 1 | No |
| *Gypsophila huashanensis* Tsui & D.Q.Lu | H | CT | M | Mountain slopes, roadside grasslands, and rock crevice | P | 600–2600 | H | China | 2 | Yes |
| *Gypsophila imbricata* Rupr. | W | CT | M | Calcareous rocks and rocky slopes | P | 200-2700 | H | Central Caucasus | 1 | No |
| *Gypsophila intricata* Franch. | W | T | M | Clay and calcareous hills, rubble slopes | P | 2000-2400 | H | Central Asia. | 2 | No |
| *Gypsophila iranica* Barkoudah | H | CT | M | Calcareous or limestone slopes | A | 2000 | H | NW. Iran | 1 | No |
| *Gypsophila krascheninnikovii* Schischk. | H | CT | M | Stony hills, sandy steppes, and sandy banks of rivers | P | NA | NA | Kazakhstan, Turkmenistan | 0? | No |
| *Gypsophila laricina* Schreb. | W | CT | M | On dry rocky hills | P | 500-2000 | L? | Central & E. Turkey to NW. Iran | 1 | Yes |

| | | | | | | | | | | |
|---|---|---|---|---|---|---|---|---|---|---|
| *Gypsophila polyclada var. leioclada* Rech.f. | H | CT | M | Calcareous mountain slopes | P | 1500-2500 | H | NW. Iran | 1 | Yes |
| *Gypsophila lepidioides* Boiss. | W | CT | M | Dry gypsum hills | P | 1391 | L | Turkey | 1 | No |
| *Gypsophila leucochleana* Hub.-Mor. | H | CT | M | Limestone hills | P | 1000-1200 | L | Turkey | 1 | No |
| *Gypsophila libanotica* Boiss. | W | CT | M | Rocky calcareous slopes | P | 1000-2500 | H | Iraq, Lebanon-Syria, Turkey | 0 | Yes |
| *Gypsophila licentiana* Hand.-Mazz. | W | CT | M | Dry stony hills, on rocks | P | 1950 | H | NW. & N. China to Mongolia | 2 | No |
| *Gypsophila lignosa* Hemsl. & Lace | W | CT | M | Fissures of calcareous rocks | P | 2000-3000 | H | E. Afghanistan to W. Pakistan | 2 | No |
| *Gypsophila linearifolia* (Fisch. & C.A.Mey.) Boiss. | H | CT | N | Gypsum hills and desertic steppes | A | 350-1250 | L | E. Medit. to Central Asia and Iran | 0 | Yes |
| *Gypsophila litwinowii* Koso-Pol. | W | CT | M | Steppe and hills, calcareous soil | P | NA | NA | European Russia | 2 | No |
| *Gypsophila lurorum* Rech.f. | H | CT | M | On dry rocky hills | A/P | 1200-2300 | H | NW. & W. Iran | 1 | No |
| *Gypsophila macedonica* Vandas | W | CT | NA | NA | P | NA | NA | North Macedonia | 2 | No |
| *Gypsophila melampoda* Bien. ex Boiss. | H | CT* | | Cultivated and Fallow Fields (Calcareous ??????) | A or P | 2000> | H | W. Iran | | Yes |

| | | | | | | | | | | |
|---|---|---|---|---|---|---|---|---|---|---|
| *Gypsophila meyeri* Rupr. | H | CT | M | Calcareous rocks and stony hills | P | NA | NA | NW. Caucasus | 1 | No |
| *Gypsophila modesta* Bornm. | H | CT | NA | NA | A | NA | NA | N. Iran | 1 | No |
| *Gypsophila mongolica* Barkoudah | W | CT | NA | Dry semidesert plains | P | NA | NA | Mongolia | 2 | No |
| *Gypsophila mozaffarianii* Negaresh | W | CT | M | Fissures of calcareous rocks | P | 1600-2500 | H | Iran | 1 | No |
| *Gypsophila mucronifolia* Rech.f. | H | CT | N | Desertic gypsum soil | P | NA | NA | Central Iran | 1 | No |
| *Gypsophila munzurensis* Armagan | H | CT | N | Open fields of oak forest and slopes | A | 1060−1080 | L | E. Central Turkey | 1 | No |
| *Gypsophila nabelekii* Schischk. | W | CT | M | Calcareous, schist, and serpentine rocks | P | 900-3800 | H | E. Turkey to NW. Iran | 1 | Yes |
| *Gypsophila nana* Bory & Chaub. | W | CT | M | Rocks and stony slopes | P | 1500-2500 | H | Greece, Peloponnesus, Crete | 2 | Yes |
| *Gypsophila neoszovitsiana* Lazkov | H | CT | NA | NA | P | NA | NA | Transcaucasus | 1 | No |
| *Gypsophila nodiflora* (Boiss.) Barkoudah | H | T | N | Eroded shaley banks | P | 800-1067 | L | E. Central Turkey | 1 | Yes |
| *Gypsophila oblanceolata* Barkoudah | H | CT | N | Salt Marshes | P | 950-110 | L | Turkey | 1 | Yes |
| *Gypsophila oldhamiana* Miq. | H | CT | N | Rocks and stony slopes | P | 1500> | L | Central China to Korea | 2 | Yes |
| *Gypsophila olympica* Boiss. | W | CT | M | Calcareous rocks | P | 2000-2500 | H | NW. Turkey | 1 | No |

| | | | | | | | | | | |
|---|---|---|---|---|---|---|---|---|---|---|
| *Gypsophila osmangaziensis* Ataslar & Ocak | H | CT | N | Collected only fromOsman gazi University Meselik campus | P | 810 | L | Turkey | 1 | No |
| *Gypsophila pacifica* Kom. | H | CT | M | Rock crevices, rocky slopes, along shores | P | 8-2000 | L | Russian Far East to N. Korea | 2 | Yes |
| *Gypsophila pallasii* Ikonn. | W | CT | N | Rock crevices, rocky slopes | P | 10-? | L | SE. Europe to NW. Caucasus | 0 | Yes |
| *Gypsophila pallida* Stapf | W | CT | M | Rocky, shaley and dry slopes | P | 1200-2700 | H | E. Turkey to W. & Central Iran | 1 | Yes |
| *Gypsophila paniculata* L. | H | CT | N | Sandy and calcareous hills, steppes | P | 0-2100 | L | E. Central Europe to W. Mongolia | 0 | Yes |
| *Gypsophila papillosa* Porta | H | CT | N | Dry, stony hills, calcareous soil | P | 50-200 | L | N. Italy | 2 | No |
| *Gypsophila parva* Barkoudah | H | CT | M | Gypsum hills | P | 800-1000 | L | N. Turkey | 1 | No |
| *Gypsophila patrinii* Ser. | W | CT | M | Dry hills, sandy soil and rocks on river banks | P | 670-2000 | L | Siberia to Russian Far East and N. China | 2 | Yes |
| *Gypsophila perfoliata* L. | H | CT | N | Solonyets, sandy meadows, fallow Fields | P | 0-1000 | L | SE. Europe to Mongolia and Iran | 0 | Yes |

| | | | | | | | | | | |
|---|---|---|---|---|---|---|---|---|---|---|
| *Gypsophila persica* Barkoudak | H | CT | M | Subdesertic dry hills, and mountain slopes | P | 2000-3000 | H | NW. Iran, and E. Iraq | 1 | Yes |
| *Gypsophila peshmenii* Güner | W | CT | M | Limestone slopes, fissures of calcareous rocks | P | 1800-2600 | H | Turkey | 1 | No |
| *Gypsophila petraea* (Baumg.) Rchb. | W | CT | M | On rocks and rocky slopes | P | 1500-2500 | H | Romania (E. & S. Carpathians) to Bulgaria | 2 | Yes |
| *Gypsophila pilosa* Huds. | H | CT | N | Ruderal fields, rocky, shaley and dry slopes | A | 500-2000 | L | Tunisia to W. & Central Asia and W. Himalaya | 0 | Yes |
| *Gypsophila pilulifera* Boiss. & Heldr. | W | CT | N | Along the edges of pine forests and corn fields | P | 2000 | H | SW. Turkey | 1 | Yes |
| *Gypsophila pinifolia* Boiss. & Hausskn. | W | T | M | Calcareous rocks, rocky slopes | P | 1000-1500 | L | E. Central Turkey | 1 | Yes |
| *Gypsophila platyphylla* Boiss. | H | T | M | Schist rocks, and rocky slopes | P | 500-2000 | L | N. & NE. Iraq to W. Iran. | 1 | No |
| *Gypsophila polyclada* Fenzl ex Boiss. | H | CT | N | Stony calcareous slopes, and fallow fields | P | 1000-2500 | H | Iran, and E. Iraq | 1 | Yes |
| *Gypsophila preobrashenskii* Czerniak. | W | CT | M | Rocky hills | P | NA | NA | Central Asia | 2 | No |

| | | | | | | | | | | |
|---|---|---|---|---|---|---|---|---|---|---|
| *Gypsophila pseudomelampoda* Gauba & Rech.f. | H | CT | M | Hills and Mountains | P | 1000-2200 | H | SW. & Central Iran | 1 | No |
| *Gypsophila pseudopallida* Falat., Assadi & F.Ghahrem. | W | CT | M | Rocky slopes | P | 1400-2000 | H | Iran | 1 | No |
| Gypsophila *pulvinaris* Rech.f. | W | CT | M | On rocks and rocky slopes | P | 1400-3000 | H | E. Turkey to Iran | 1 | Yes |
| *Gypsophila repens* L. | W | CT | M | Rocky hills, slopes, clay | P | 2000 | H | Mountains of Central & S. Europe | 2 | Yes |
| *Gypsophila reuteri* (Boiss. & Hausskn.) F.N.Williams | W | T | M | Rocks | P | 1500 | L | S. Turkey | 1 | No |
| *Gypsophila robusta* Grossh. | H | CT | N | River Banks | P | NA | NA | Transcaucasus | 1 | No |
| *Gypsophila rupestris* Kupr. | W | CT | M | NA | P | NA | NA | SE. European Russia to Kazakhstan | 2 | No |
| Gypsophila ruscifolia Boiss. | H | CT | M | Rocky and calcareous hills, dry stony hills and slopes, quercus forests | P | 500-2000 | H | E. Turkey to W. Iran | 1 | Yes |
| *Gypsophila saligna* Schrad. | | | | | | | | | | |
| *Gypsophila sambukii* Schischk. | W | CT | N | Bare patches in the arctic part of siberia | P | 1750 | H | Central & E. Siberia to Central Russian Far East | 2 | No |
| *Gypsophila saponarioides* Bornm. & Gauba | W | CT | M | Calcareous rocks | P | 2500-3000 | H | Iran | 1 | Yes |

| | | | | | | | | | | |
|---|---|---|---|---|---|---|---|---|---|---|
| *Gypsophila scorzonerifolia* Ser. | H | CT | N | Wet sandy places | P | 0-1700 | L | E. Ukraine to Caucasus | 0 | Yes |
| *Gypsophila serpylloides* Boiss. & Heldr. | W | CT | M | High dry mountain pastures | P | 2000-2600 | H | Turkey | 1 | No |
| *Gypsophila silenoides* Rupr. | H | CT | M | Lime screes, rocky banks and slopes, alpine pastures | P | 1400-2200 | H | NE. Turkey to W. Caucasus | 1 | Yes |
| *Gypsophila simonii* Hub.-Mor. | H | CT | M | Gypsum slopes | P | 700 | L | N. Turkey | 1 | No |
| *Gypsophila simulatrix* Bornm. & Woronow | H | CT | M | Crevices of calcareous rocks, rocky slopes | P | 500-2000 | H | NE. Turkey to Caucasus | 1 | Yes |
| *Gypsophila spathulifolia* (Fisch. & C.A.Mey.) Fenzl | H | CT | M | Stony hills | A | NA | NA | Central Asia | 2 | No |
| *Gypsophila spinosa* D.Q.Lu | H | CT | M | Sandy hills | P | 500-1000 | L | N. Xinjiang (Altay Shan) | 2 | No |
| *Gypsophila steupii* Schischk. | H | CT | NA | Rocks | P | NA | NA | W. Caucasus | 1 | No |
| *Gypsophila struthium* Loefl. | W | CT | M | Arid calcareous hills, river banks | P | 1000> | L | Central & SE. Spain, Morocco | 2 | Yes |
| *Gypsophila syriaca* Schischk. | W | CT | M | Calcareous rocks | P | 1500-2500 | H | SE. Turkey | 1 | No |
| *Gypsophila szovitsii* Fisch. & C.A.Mey. ex Fenzl | W | CT | M | Clay Hills | P | 1000-1400 | L | Caucasus | 1 | Yes |
| *Gypsophila takhtadzhanii* Schischk . ex Ikonn. | W | CT | M | Calcareous rocks, rocky slopes | P | 1400-2000 | H | S. Transcaucas us | 1 | Yes |
| *Gypsophila tenuifolia* M.Bieb. | W | CT | M | Rocks, rocky soil, | P | 1400-2700 | H | NE. Turkey to Caucasus | 1 | Yes |

| | | | | | | | | | | |
|---|---|---|---|---|---|---|---|---|---|---|
| | | | | alpine meadows | | | | | | |
| *Gypsophila tomentosa* L. | H | CT | N | Gypsum-rich soils in wet habitats | P | 1000> | L | Central & SE. Spain | 2 | Yes |
| *Gypsophila torulensis* Koç | H | CT | M | Slopes with calcareous rocks | A | 1100 | L | Turkey | 1 | No |
| *Gypsophila transalaica* Ikonn. | NA | NA | NA | NA | P | NA | NA | Central Asia | 2 | No |
| *Gypsophila tschiliensis* J.Krause | W | CT | M | Forest margins, mountain slopes | P | 2000-3000 | H | China | 2 | No |
| *Gypsophila tuberculosa* Hub.-Mor. | W | CT | M | Steppes, dry river banks | P | 1000-2000 | H | E. Turkey | 1 | Yes |
| *Gypsophila tubulifera* Bornm. | H | T | NA | NA | P | NA | NA | Lebanon | 2 | No |
| *Gypsophila tubulosa* (Jaub. & Spach) Boiss. | H | T | M | Stony slopes, and schist | A | 1000> | L | W. Turkey and Greece | 0 | No |
| *Gypsophila turcica* Hamzaoglu | W | CT | M | on Gypsaceous hills | P | 1700-1900 | H | Turkey | 1 | No |
| *Gypsophila umbricola* (J.R.I.Wood ) R.A.Clement | W | CT | M | Wet vertical and subvertical sanstone, limestone, or basalt cliffs | P | 1800-2800 | H | Arabian Peninsula | 2 | No |
| *Gypsophila uralensis* Less. | W | CT | M | Rocky hills, rock crevices | P | 1200-2000 | H | European Russia to W. Siberia | 2 | Yes |
| *Gypsophila vaccaria* (L.) Sm. | H | T | N | Meadows, stepps, rocky areas | A | 0–2400 | H | Macaronesia, Central & E. Europe to Medit. and | 0 | Yes |

| | | | | | | | | | | |
|---|---|---|---|---|---|---|---|---|---|---|
| | | | | | | | | Central Himalaya | | |
| *Gypsophila vedeneevae* Lepeschk. ex Botsch. & Vved. | W | CT? | M? | NA (probably on rocks and rocky slopes) | P | NA | NA | Central Asia | 2 | No |
| *Gypsophila venusta* Fenzl | H | T | N | Fallow fields, cultivated land, and steppe | P | 700-1400 | L | N. Syria and S. Turkey | 0 | Yes |
| *Gypsophila villosa* Barkoudah | H | T | M | On bare gypsum hill | P | 1850 | H | Central Asia | 2 | No |
| *Gypsophila vinogradovii* Safonov | NA | NA | NA | NA | P | NA | NA | Central Caucasus | 1 | No |
| *Gypsophila virgata* Boiss. | W | CT | M | Dry hills of metamorphic rocks | P | 1000-2200 | H | E. Turkey to W. Iran | 1 | Yes |
| *Gypsophila viscosa* Murray | H | CT | N | Sandy valleys, wadi banks, and fallow fields | A | 1700> | L | E. Medit. to Arabian Peninsula | 2 | Yes |
| *Gypsophila volgensis* Krasnova | W | CT | NA | NA | P | NA | NA | E. Europe | 2 | No |
| *Gypsophila wendelboi* Rech.f. | W | CT | M | Fissures of calcareous rocks | P | 1800 | H | E. Afghanistan | 2 | No |
| *Gypsophila wilhelminae* Rech.f. | W | CT | M | Calcareous rocks and rocky slope | P | 2000-2500 | H | NW. Iran | 1 | Yes |
| *Gypsophila xanthochlora* Rech.f. | H | CT | M | Calcareous rocks, and dry slopes | P | 1000-2000 | H | Iran | 1 | No |
| *Gypsophila yazdiana* Falat., F.Ghahrem. & Assadi | H | CT | M | Rocky slopes of a mountainous area | P | 2800-3000 | H | Iran | 1 | No |

| | | | | | | | | | | |
|---|---|---|---|---|---|---|---|---|---|---|
| *Gypsophila yusufeliensis* Budak | W | CT | M | Siliceous rock crevices | P | 1250-1260 | L | Turkey | 1 | No |

**Aguirre-Santoro J, Salinas NR, Michelangeli FA**. **2019**. The influence of floral variation and geographic disjunction on the evolutionary dynamics of Ronnbergia and Wittmackia (Bromeliaceae: Bromelioideae). *Botanical journal of the Linnean Society. Linnean Society of London* **192**: 609–624.

**Anthony F, Diniz LEC, Combes M-C, Lashermes P**. **2010**. Adaptive radiation in Coffea subgenus Coffea L. (Rubiaceae) in Africa and Madagascar. *Plant systematics and evolution = Entwicklungsgeschichte und Systematik der Pflanzen* **285**: 51–64.

**Baldwin BG, Sanderson MJ**. **1998**. Age and rate of diversification of the Hawaiian silversword alliance (Compositae). *Proceedings of the National Academy of Sciences of the United States of America* **95**: 9402–9406.

**Bell CD, Donoghue MJ**. **2005**. Phylogeny and biogeography of Valerianaceae (Dipsacales) with special reference to the South American valerians. *Organisms, diversity & evolution* **5**: 147–159.

**Brennan IG, Oliver PM**. **2017**. Mass turnover and recovery dynamics of a diverse Australian continental radiation. *Evolution; international journal of organic evolution* **71**: 1352–1365.

**Burge DO, Erwin DM, Islam MB, Kellermann J, Kembel SW, Wilken DH, Manos PS**. **2011**. Diversification of Ceanothus (Rhamnaceae) in the California Floristic Province. *International journal of plant sciences* **172**: 1137–1164.

**Calonje M, Meerow AW, Griffith MP, Salas-Leiva D, Vovides AP, Coiro M, Francisco-Ortega J**. **2019**. A Time-Calibrated Species Tree Phylogeny of the New World Cycad Genus Zamia L. (Zamiaceae, Cycadales). *International journal of plant sciences* **180**: 286–314.

**Calsbeek R, Thompson JN, Richardson JE**. **2003**. Patterns of molecular evolution and diversification in a biodiversity hotspot: the California Floristic Province. *Molecular ecology* **12**: 1021–1029.

**Cox SC, Prys-Jones RP, Habel JC, Amakobe BA, Day JJ**. **2014**. Niche divergence promotes rapid diversification of East African sky island white-eyes (Aves: Zosteropidae). *Molecular ecology* **23**: 4103–4118.

**Ebersbach J, Muellner-Riehl AN, Michalak I, Tkach N, Hoffmann MH, Röser M, Sun H, Favre A**. **2017a**. In and out of the

Qinghai-Tibet Plateau: divergence time estimation and historical biogeography of the large arctic-alpine genus*Saxifraga*L. *Journal of biogeography* **44**: 900–910.

**Ebersbach J, Schnitzler J, Favre A, Muellner-Riehl AN**. **2017b**. Evolutionary radiations in the species-rich mountain genus Saxifraga L. *BMC evolutionary biology* **17**: 119.

**Esselstyn JA, Timm RM, Brown RM**. **2009**. Do geological or climatic processes drive speciation in dynamic archipelagos? The tempo and mode of diversification in Southeast Asian shrews. *Evolution; international journal of organic evolution* **63**: 2595–2610.

**Hughes C, Eastwood R**. **2006**. Island radiation on a continental scale: exceptional rates of plant diversification after uplift of the Andes. *Proceedings of the National Academy of Sciences of the United States of America* **103**: 10334–10339.

**Jabbour F, Renner SS**. **2012**. A phylogeny of Delphinieae (Ranunculaceae) shows that Aconitum is nested within Delphinium and that Late Miocene transitions to long life cycles in the Himalayas and Southwest China coincide with bursts in diversification. *Molecular phylogenetics and evolution* **62**: 928–942.

**Janssens SB, Vandelook F, De Langhe E, Verstraete B, Smets E, Vandenhouwe I, Swennen R**. **2016**. Evolutionary dynamics and biogeography of Musaceae reveal a correlation between the diversification of the banana family and the geological and climatic history of Southeast Asia. *The New phytologist* **210**: 1453–1465.

**Joly S, Heenan PB, Lockhart PJ**. **2009**. A Pleistocene inter-tribal allopolyploidization event precedes the species radiation of Pachycladon (Brassicaceae) in New Zealand. *Molecular phylogenetics and evolution* **51**: 365–372.

**Joly S, Heenan PB, Lockhart PJ**. **2014**. Species radiation by niche shifts in New Zealand's rockcresses (Pachycladon, Brassicaceae). *Systematic biology* **63**: 192–202.

**Klak C, Reeves G, Hedderson T**. **2004**. Unmatched tempo of evolution in Southern African semi-desert ice plants. *Nature* **427**: 63–65.

**Lagomarsino LP, Condamine FL, Antonelli A, Mulch A, Davis CC**. **2016**. The abiotic and biotic drivers of rapid diversification in A ndean bellflowers (Campanulaceae). *The New phytologist* **210**: 1430–1442.

**Linder HP**. **2003**. The radiation of the Cape flora, southern Africa. *Biological reviews of the Cambridge Philosophical Society* **78**: 597–638.

**Linder HP, Hardy CR**. **2004**. Evolution of the species-rich Cape flora. *Philosophical transactions of the Royal Society of London. Series B, Biological sciences* **359**: 1623–1632.

**McCullough JM, Oliveros CH, Benz BW, Zenil-Ferguson R, Cracraft J, Moyle RG, Andersen MJ**. **2022**. Wallacean and Melanesian Islands Promote Higher Rates of Diversification within the Global Passerine Radiation Corvides. *Systematic biology* **71**: 1423–1439.

**Meseguer AS, Aldasoro JJ, Sanmartín I**. **2013**. Bayesian inference of phylogeny, morphology and range evolution reveals a complex evolutionary history in St. John's wort (Hypericum). *Molecular phylogenetics and evolution* **67**: 379–403.

**Moore W, Robertson JA**. **2014**. Explosive adaptive radiation and extreme phenotypic diversity within ant-nest beetles. *Current biology: CB* **24**: 2435–2439.

**Nge FJ, Biffin E, Thiele KR, Waycott M**. **2021**. Reticulate Evolution, Ancient Chloroplast Haplotypes, and Rapid Radiation of the Australian Plant Genus Adenanthos (Proteaceae). *Frontiers in Ecology and Evolution* **8**.

**Nürk N, Scheriau C, Madriñán S**. **2013**. Explosive radiation in high Andean Hypericum—rates of diversification among New World lineages. *Frontiers in genetics* **4**.

**Richardson JE, Weitz FM, Fay MF, Cronk QC, Linder HP, Reeves G, Chase MW**. **2001**. Rapid and recent origin of species richness in the Cape flora of South Africa. *Nature* **412**: 181–183.

**Rivera VL, Panero JL, Schilling EE, Crozier BS, Moraes MD**. **2016**. Origins and recent radiation of Brazilian Eupatorieae (Asteraceae) in the eastern Cerrado and Atlantic Forest. *Molecular phylogenetics and evolution* **97**: 90–100.

**Smíd J, Carranza S, Kratochvíl L, Gvoždík V, Nasher AK, Moravec J**. **2013**. Out of Arabia: a complex biogeographic history of multiple vicariance and dispersal events in the gecko genus Hemidactylus (Reptilia: Gekkonidae). *PloS one* **8**: e64018.

**Tolley KA, Chase BM, Forest F**. **2008**. Speciation and radiations track climate transitions since the Miocene Climatic Optimum: a case study of southern African chameleons. *Journal of biogeography* **35**: 1402–1414.

**Toussaint EFA, Hendrich L, Shaverdo H, Balke M**. **2015**. Mosaic patterns of diversification dynamics following the colonization of Melanesian islands. *Scientific reports* **5**: 16016.

**Valente LM, Savolainen V, Vargas P**. **2010**. Unparalleled rates of species diversification in Europe. *Proceedings. Biological* *sciences / The Royal Society* **277**: 1489–1496.

**Van Bocxlaer I, Biju SD, Loader SP, Bossuyt F**. **2009**. Toad radiation reveals into-India dispersal as a source of endemism in the Western Ghats-Sri Lanka biodiversity hotspot. *BMC evolutionary biology* **9**: 131.

**Wood PL Jr, Heinicke MP, Jackman TR, Bauer AM**. **2012**. Phylogeny of bent-toed geckos (Cyrtodactylus) reveals a west to east pattern of diversification. *Molecular phylogenetics and evolution* **65**: 992–1003.

**Yamada T, Sugiyama T, Tamaki N, Kawakita A, Kato M**. **2009**. Adaptive radiation of gobies in the interstitial habitats of gravel beaches accompanied by body elongation and excessive vertebral segmentation. *BMC evolutionary biology* **9**: 145.

**Ye X-Y, Ma P-F, Yang G-Q, Guo C, Zhang Y-X, Chen Y-M, Guo Z-H, Li D-Z**. **2019**. Rapid diversification of alpine bamboos associated with the uplift of the Hengduan Mountains. *Journal of biogeography* **46**: 2678–2689.
